# Regulatory architecture tunes gene expression noise in the plant heat shock response

**DOI:** 10.64898/2026.10.22.760979

**Authors:** Rituparna Goswami, Dora Cano Ramirez, Katie Abley, Mana Afsharinafar, Connor Reynolds, Jiayi Zeng, Marcel Bhattarai, Fiona Jingxuan Zhang, Sandra Cortijo, Anthony Hall, Enrico Sandro Colizzi, James C. W. Locke

**Affiliations:** Sainsbury Laboratory, University of Cambridge, Cambridge, UK; Earlham Institute, Norwich Research Park, Norwich, UK; Max Planck Institute for the Physics of Complex Systems (MPI-PKS), Dresden, Germany; IPSiM, Univ Montpellier, CNRS, INRAE, Institut Agro, Montpellier, France; School of Biological Sciences, University of East Anglia, Norwich, UK; Inria Centre de Lyon, Team Biotic, Villeurbanne, France

**Keywords:** *Arabidopsis thaliana*, Heat stress, Transcriptional noise, Systems biology, Single-cell measurements

## Abstract

Cells within the same tissue can respond differently to the same stress, yet how gene regulatory networks generate and control this variability remains unclear. Here, we show that cell-to-cell variation in the Arabidopsis heat shock response is coordinated across the heat shock regulon and that the distribution of responses is tuned by the underlying regulatory network. Imaging of an *HSP70* promoter reporter revealed basal activity in a subset of root cells and heterogeneous induction after heat shock, with some cells remaining near pre-stress expression levels. Dual reporters showed strong coordination between independently integrated *HSP70* reporter loci, consistent with substantial shared upstream regulatory variation. Single-nucleus RNA sequencing confirmed heterogeneous but coordinated activation of heat shock genes across root cell types. Stronger heat shock states at both ambient and heat stress temperatures were associated with a recurrent thermoprotective programme across root cell populations, whereas weak responders after heat shock were associated with distinct cell-type specific transcriptional programmes, including defence- and jasmonate-associated states in some root cell types. Stochastic modelling suggested that HSFA2-dependent amplification could broaden the distribution of *HSP70* expression across cells. Consistent with this prediction, experiments showed that loss of HSFA2 reduced *HSP70* induction and lowered cell-to-cell variability after controlling for differences in mean expression. Together, these results show that the plant heat shock network controls both the magnitude and distribution of stress-responsive gene expression across cells.

## Introduction

Stress responses are often defined from population averaged measurements, yet this can obscure substantial non-genetic variation between individual cells. In plants, cells remain fixed within structured tissues during stress, sharpening the question of how heterogeneous responses arise between neighbouring cells exposed to a shared environment. Although cell-to-cell variation in plant gene expression has been observed, including in constitutive reporters and auxin-regulated developmental programmes (1–3), how stress response networks reshape this variation during environmental change remains unclear.

The heat shock response provides a well-defined system in which to address these questions. Increases in temperature can destabilise proteins and impair cellular functions (4–6). The canonical heat shock response (HSR) is characterised by the rapid induction of chaperones such as HSP70, HSP101 and small heat shock proteins, which limit protein aggregation and aid refolding (7–12). This response is controlled by a network of heat shock transcription factors (HSFs), which bind heat shock elements in the promoters of target genes (13–17). In Arabidopsis,

HSFA1-family proteins act as central upstream regulators of the heat shock response (18, 19) and are kept inactive under non-stress conditions through interactions with HSP70 and HSP90 chaperones (7, 20). Following heat stress, this repression is relieved, allowing HSFA1-dependent regulation of heat shock response genes, including HSFA2 (13, 19, 21, 22). HSFA2 regulates a partially overlapping set of heat-responsive genes and further amplifies chaperone gene expression (14, 23–25). The heat shock network therefore contains feedback and amplification steps that could, in principle, influence not only mean expression but also the distribution of responses across cells.

Live single-cell transcriptional imaging has shown that plant stress responses can be highly heterogeneous (26, 27). In the heat shock response specifically, imaging of nascent transcription showed that reporters for genes such as *HSP101* are activated heterogeneously across Arabidopsis cells, with tissue-level induction arising largely through changes in the fraction of transcriptionally active nuclei rather than graded increases in transcription rate across all cells (26). Analysis of *HSP101* reporter activity integrated over relatively short timescales during heat stress further showed that intrinsic, allele-specific processes accounted for most of the variability in predicted mRNA production. Complementary single-cell transcriptomic analysis of Arabidopsis roots showed strong induction of canonical heat shock genes across cell types, together with subtler cell-type specific transcriptional responses (28). These studies establish substantial heterogeneity in the plant heat shock response, but leave open how variation in accumulated heat shock gene expression is organised among cells within a tissue: in particular, whether heat shock genes vary coordinately within cell types and with broader transcriptional programmes, and how the regulatory architecture of the heat shock network shapes the distribution of responses.

Here, we combine quantitative live imaging, dual-reporter analysis, single-nucleus transcriptomics, stochastic modelling and genetic perturbation to determine how cell-to-cell heterogeneity in the Arabidopsis heat shock response is organised and regulated. We find that *HSP70* expression heterogeneity predominantly reflects shared upstream differences between cells and is coordinated across the endogenous heat shock regulon and broader transcriptional programmes. We further show that loss of HSFA2 reduces cell-to-cell variability in HSP70 induction after accounting for differences in mean expression, consistent with HSFA2-dependent amplification broadening the response distribution. Together, these results establish cell-to-cell heterogeneity as a regulated feature of the plant heat shock response.

## Results

### *HSP70* reporter expression is heterogeneous across cells under both basal and heat stress conditions

To measure cell-to-cell variability in the heat shock response, we imaged transgenic *Arabidopsis thaliana* seedlings carrying an *HSP70* promoter driving a nuclear-localised mTurquoise2 fluorescent reporter (*pHSP70::2xmTurq-N7*), alongside a constitutive *35S::H2B-RFP* nuclear marker (29). Five- to six-day-old plants were grown and imaged in a sealed imaging chamber that maintained humidity and nutrient access (see Methods, Fig. S1).

Seedlings were first imaged at 22 °C (T0), then exposed to 35 °C for 30 minutes (30 min), 1 hour (1 h), or 2 h, and re-imaged 4 h (T1) and 24 h (T2) after the end of heat treatment. At each time point, we used a large field of view to image the root from the tip to approximately 5–7 mm shootward of the tip, adjusting the imaging position between time points to account for root growth. We imaged the outermost visible root tissues, typically within the first one to two cell layers from the root surface. Because mTurquoise2 is a stable fluorescent protein (30), fluorescence reflects accumulated promoter activity rather than instantaneous transcriptional bursts, allowing us to quantify sustained heterogeneity across individual nuclei. We used the constitutive RFP nuclear marker to identify nuclei and quantified *pHSP70::2xmTurq-N7* fluorescence in each nucleus at each time point (see Methods).

Before stress, detectable *HSP70* reporter fluorescence was restricted to a subset of nuclei in each root (Fig. 1B,D). Heat stress induced reporter activity in a larger fraction of nuclei, but the response remained highly heterogeneous, with some nuclei showing strong activation while others remained weakly induced or close to pre-stress levels (Fig. 1B,D). This effect was already evident after 30 min heat stress and was maintained after longer heat treatments (Fig. 1F,H). Heterogeneity in reporter fluorescence remained evident 24 h after stress. Quantification across roots showed that heat treatment increased both mean reporter expression and the coefficient of variation across nuclei (Fig. 1D,F,H), indicating that heat stress not only increases *HSP70* promoter activity but also broadens the distribution of responses between cells. By contrast, the constitutive *35S::H2B-RFP* nuclear marker showed little change in either variability or mean intensity across the same time course (Fig. 1C,E,G,I), supporting the conclusion that the heterogeneous increase is specific to the *HSP70* reporter rather than a general consequence of imaging or nuclear fluorescence changes. Substantial differences were evident even between neighbouring nuclei, suggesting that local position alone cannot explain the heterogeneous response. Additional independent 1 h heat stress datasets acquired on a second microscope, together with an independent *HSP70* reporter insertion line, reproduced this qualitative behaviour (Figs. S2 and S3). Detailed analyses of the 30 min and 2 h datasets summarised in Fig. 1 are shown in Figs. S4 and S5.

**Fig. 1.**
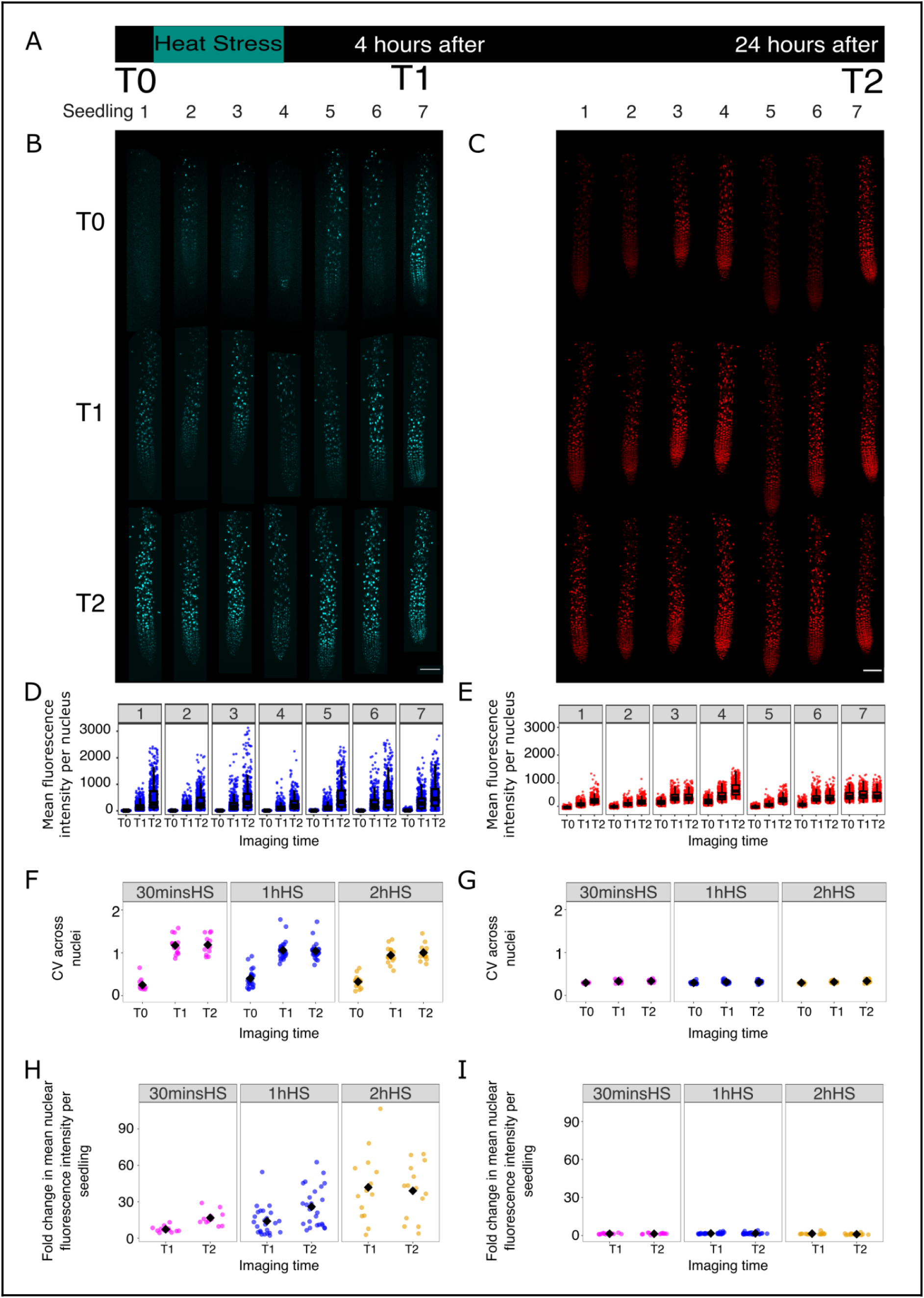
*HSP70* promoter activity is heterogeneous under basal conditions and increases heterogeneously after heat stress. **(A)** Experimental design. Seedlings were imaged before heat stress, exposed to heat stress at 35 °C for 30 min, 1 h, or 2 h, and re-imaged 4 h and 24 h later. **(B, C)** Representative confocal images of roots expressing *pHSP70::2xmTurq-N7* (B) and the constitutive nuclear marker *35S::H2B-RFP* (**C**) before heat stress and at 4 h and 24 h after 1 h heat stress. Low basal *HSP70* reporter activity is visible in a subset of nuclei before stress, whereas heat stress induces a stronger but highly heterogeneous response across cells. Images are shown with adjusted display ranges for visualisation: mTurquoise2, T0, 200–250; T1 and T2, 100–2000; RFP, T0, T1 and T2, 200–1000. Scale bar is 100 micron. **(D, E)** Quantification of nuclear fluorescence intensity for the *pHSP70::2xmTurq-N7* reporter (**D**) and *35S::H2B-RFP* control (**E**) in the seven seedlings shown in **B** and **C**. Each point represents one nucleus. **(F, G)** Cell-to-cell variability, measured as the coefficient of variation across nuclei within each seedling, for the *pHSP70::2xmTurq-N7* reporter (**F**) and *35S::H2B-RFP* control (**G**) after 30 min, 1 h, or 2 h heat stress. **(H, I)** Fold change in mean nuclear fluorescence intensity per seedling relative to the pre-stress time point for the *pHSP70::2xmTurq-N7* reporter (**H**) and *35S::H2B-RFP* control (**I**). Heat stress increases mean *HSP70* reporter fluorescence, whereas the constitutive nuclear marker does not show a comparable increase. For F–I, data are pooled from independent biological repeats: n = 12, 26 and 15 seedlings from 2, 4 and 2 independent imaging experiments for the 30 min, 1 h and 2 h heat stress treatments, respectively. Each point represents one seedling; black diamonds indicate the mean. The heat stress duration datasets were acquired in separate imaging experiments. The 30 min dataset used different mTurq acquisition settings; an acquisition-setting comparison is shown in Fig. S4.

To test whether *HSP70* reporter induction was heat-specific, matched control seedlings were maintained at 22 °C and imaged over 24 hours at the same three time points (T0, T1, T2). Under these conditions, *HSP70* reporter fluorescence remained low and heterogeneous, with no systematic increase in mean intensity or variability over time (Fig. S6).

### *HSP70* reporter heterogeneity arises from upstream regulatory variation

To determine whether the observed cell-to-cell variability in *HSP70 reporter* expression arises primarily from promoter-intrinsic noise or from upstream regulatory differences between cells, we developed a dual-reporter assay. Transgenic Arabidopsis seedlings were generated carrying two independently integrated *HSP70* promoter reporters, *pHSP70::2xmTurq-N7* and *pHSP70::YFP-N7,* together with the constitutive nuclear marker *35S::H2B-RFP* (Fig. 2A,B). This design allowed us to compare *HSP70* reporter levels simultaneously in two fluorescent channels within the same nucleus.

**Fig. 2.**
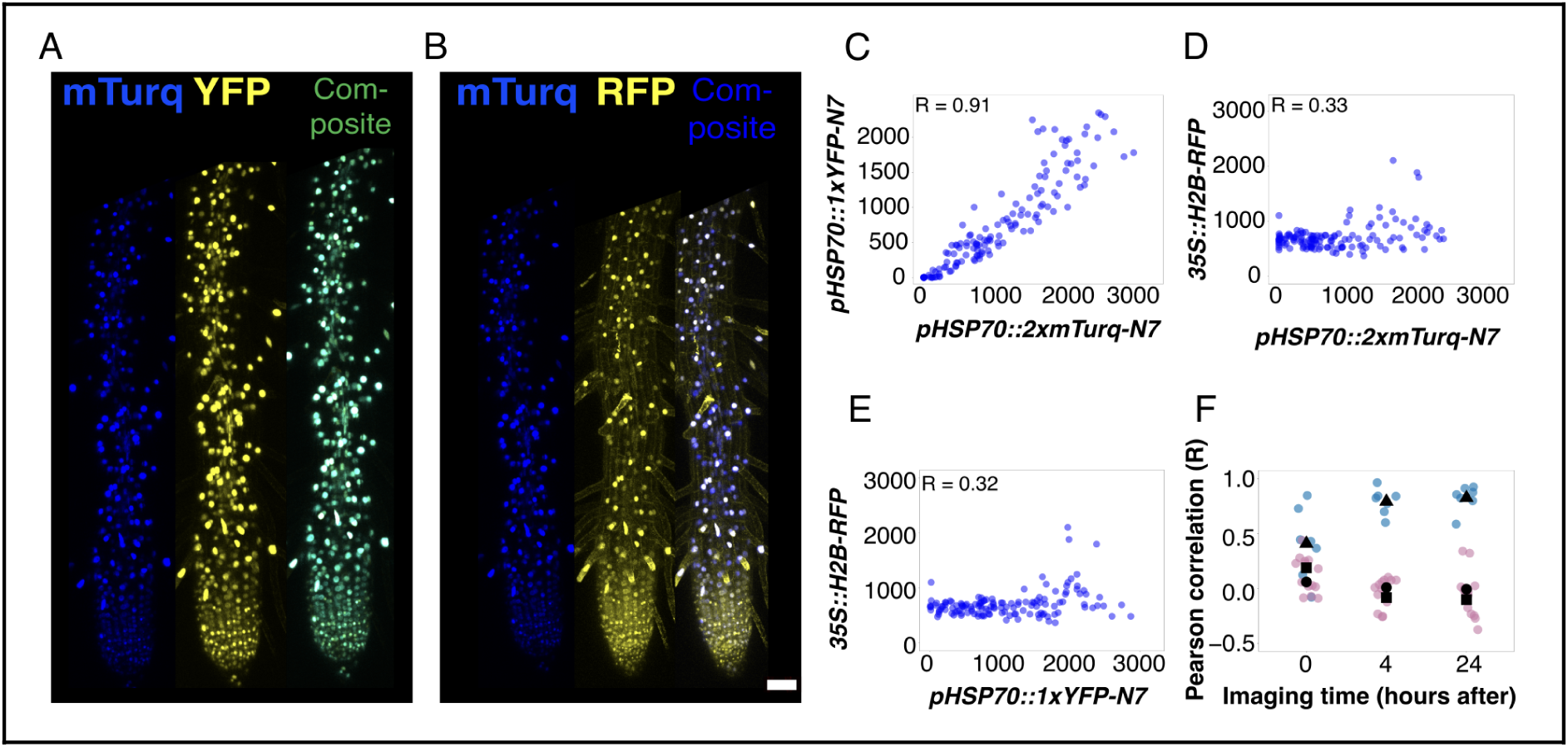
*HSP70* reporter heterogeneity is dominated by upstream regulatory variation. **(A, B)** Representative confocal images of roots 24 h after a 1 h heat stress from T1-generation primary transformants carrying *pHSP70::YFP-N7* in a previously established *pHSP70::2xmTurq-N7; 35S::H2B-RFP* reporter line. (A) mTurquoise2, YFP, and merged image. (B) mTurquoise2, RFP, and merged image. **(C)** Nuclear fluorescence intensities of *pHSP70::2xmTurq-N7* and *pHSP70::YFP-N7* are strongly correlated across individual nuclei, indicating coordinated activation of the two reporters within the same cells. **(D, E)** Nuclear fluorescence intensities of *35S::H2B-RFP* show much weaker correlation with *pHSP70::2xmTurq-N*7 (D) or *pHSP70::YFP-N7* (E), consistent with independent regulation of the constitutive marker **(F)** Pearson correlation coefficients for the paired reporter combinations across imaging time points. Blue dots show *pHSP70::2xmTurq-N7* versus *pHSP70::YFP-N7* correlations; pink dots show correlations between either *HSP70* reporter and *35S::H2B-RFP*. Black symbols indicate the mean for each comparison: triangle, mTurquoise2 reporter versus YFP reporter; circle, mTurquoise2 reporter versus RFP marker; square, YFP reporter versus RFP marker. Each dot represents one Pearson correlation coefficient calculated from one root at the indicated time point (n = 7 independent transformants); reporter pair correlations were calculated from the same roots. In C–E, each dot represents one nucleus and Pearson correlation coefficients (R) are indicated. Images in A and B are shown with adjusted display ranges for visualisation: mTurquoise2, 200–300; YFP, 200–250; RFP, 200–1200.

We imaged these dual-reporter seedlings before stress and again 4 h and 24 h after a 1 h heat stress at 35 °C. Fluorescence intensities of the two *HSP70* reporters were strongly correlated across individual nuclei at all time points after stress (Fig. 2C,F), with Pearson correlation coefficients typically in the range R = 0.80-0.95. By contrast, correlations between either *HSP70* reporter and the constitutive *35S::H2B-RFP* marker were much weaker (Fig. 2D-F), consistent with independent regulation of the control reporter.

These results indicate that heterogeneity in *HSP70* expression is dominated by a shared regulatory state, rather than independent intrinsic fluctuations at each promoter. The coordinated activation of the two *HSP70* reporters within the same nucleus argues that this regulatory state is common to both reporter loci. Together, these data support a model in which cell-to-cell variability in the heat shock response primarily reflects differences in upstream regulatory activity, rather than promoter-intrinsic stochasticity.

### Single-nucleus RNA-seq reveals coordinated yet heterogeneous activation of the endogenous heat shock regulon

To determine whether the heterogeneity observed in reporter lines is also evident in endogenous heat shock gene expression, we performed single-nucleus RNA-seq (snRNA-seq) on Arabidopsis root tips under matched control (22 °C) and heat stress (35 °C for 1 h) conditions, using a protocol optimised for high-depth transcript detection (see Methods). The main expression analyses used two fresh tissue biological replicates per condition (33,664 nuclei across the fresh samples), with an additional liquid nitrogen frozen replicate (12,147 nuclei) generated to test whether the observed heterogeneity was sensitive to sample handling. Nuclei from fresh and frozen samples were integrated for clustering and annotation (Fig. 3A), and marker gene expression supported the assignment of major root cell populations (Fig. S7, Supplementary Table. S5, see methods). We then quantified heat shock gene expression across individual nuclei in the fresh-tissue samples.

**Fig. 3.**
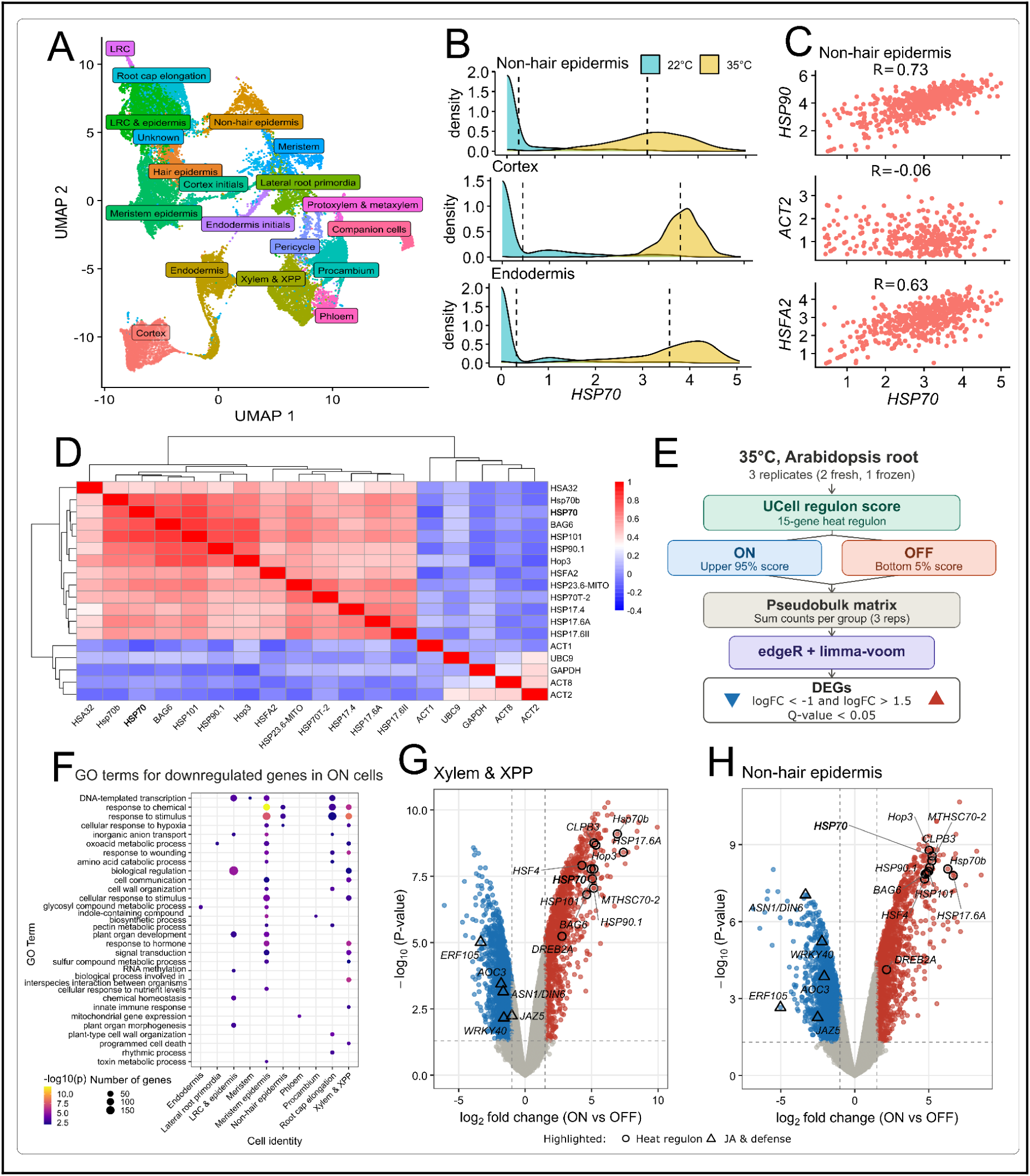
The endogenous heat shock response is activated heterogeneously but coordinately across Arabidopsis root cell types and is associated with broader transcriptional programmes. **(A)** UMAP of single nuclei from *Arabidopsis* roots, coloured by annotated cell type. Major root cell populations are represented across the dataset. **(B)** Distribution of *HSP70* expression in selected cell types in control (22 °C) and heat shock (35 °C) samples. Heat stress shifts the distribution towards higher *HSP70* expression in all cell types shown, but substantial heterogeneity remains within each population. Dashed lines indicate the mean expression level for each condition. **(C)** Single-nucleus expression correlations between *HSP70* and selected genes in non-hair epidermis nuclei from the fresh 35 °C heat stress samples. *HSP70* expression is strongly correlated with other heat shock genes, including *HSP90* and *HSFA2*, but only weakly correlated with the control gene *ACT2*. Pearson correlation coefficients (R) are indicated. **(D)** Pairwise gene-gene correlation heatmap for heat shock genes and control genes across non-hair epidermis nuclei from the fresh 35 °C heat shock samples. Colour indicates the Pearson correlation coefficient between each gene pair, with higher values indicating stronger positive correlation. Heat shock genes show coordinated positive correlation, whereas housekeeping/control genes show weaker correlations. **(E)** Workflow for the 35 °C Heat Shock Response (HSR)-score analysis. Nuclei from three biological replicates, comprising two fresh and one frozen sample, were scored using a 15-gene heat shock response (HSR) UCell signature. The lowest-scoring 5% of nuclei were designated OFF and compared with the remaining upper 95%, designated ON. Pseudobulk counts (3 replicates) were compared by edgeR and limma-voom to identify DEGs (Q < 0.05; log2FC < –1 or > 1.5) (Supplementary Datasets S1 and S2). **(F)** GO enrichment of genes relatively enriched in OFF compared with ON nuclei across annotated root cell types at 35 °C. Enriched processes differ between cell types and include defence-, jasmonate- and wound-associated responses. **(G,H)** Differential expression between ON and OFF nuclei in the combined Xylem & XPP population (**G**) and non-hair epidermis (**H**) at 35 °C. Positive log2 fold change indicates higher expression in ON nuclei and negative values indicate higher expression in OFF nuclei. Selected heat-response genes and other stress-associated genes are highlighted.

In the control (22 °C) samples, *HSP70* expression was detected within an average of 21.8% of nuclei between replicates across all root cell-types. This basal heterogeneity was consistent with the low-level reporter activity observed in some nuclei in non-stressed seedlings (Fig. 1B,D,F). Among all the nuclei with detectable *HSP70* transcripts, the mean coefficient of variation (CV) was 2.0 (+/- 0.84), compared with the mean CV of the global gene background of 0.33 (+/- 0.30) . Thus, heterogeneous endogenous *HSP70* expression was already evident under ambient conditions.

Upon heat stress, HSP70 expression increased significantly for each cell-type (EdgeR pseudobulk differential expression Q-value < 0.00001), and was detected within an average of 98.6% of nuclei. Despite this widespread induction, expression levels remained broadly distributed within cell-types, (Fig. 3B; Fig. S8), with some cells showing low expression similar to uninduced. The mean CV among nuclei with detectable *HSP70* transcripts was 0.95 ± 0.14, compared with a mean CV of 0.35 ± 0.33 for the global gene background. Heat treatment therefore increased the prevalence and abundance of *HSP70* expression with a broad expression distribution qualitatively similar to that observed in confocal measurements. Similar broad expression distributions were observed for *HSP90*, whereas *ACT2* did not show the marked heat-dependent induction observed for *HSP70* and *HSP90* (Fig. S8).

To test whether this heterogeneity reflected coordinated cell-to-cell differences in activation of the heat shock regulon, we examined correlations in gene expression across nuclei within individual cell types at 22 and 35 °C (Figs. S12 and S13). Heat shock genes showed coordinated positive correlations across nuclei in multiple cell types at both temperatures, with the pattern particularly pronounced following heat stress, whereas control genes generally showed weaker correlations (Fig. 3D; Figs. S12 and S13). Consistent with this, single-nucleus *HSP70* expression was positively correlated with other heat shock genes, including *HSP90* and *HSFA2*, but only weakly correlated with the control gene *ACT2* (Fig. 3C; Figs. S9, S10, S12 and S13). Similar broad *HSP70* expression distributions and heat shock gene covariation were observed in the frozen replicate, indicating that these patterns were not specific to the fresh-nuclei isolation procedure (Fig. S11). Together, these data show that cell-to-cell heterogeneity in the heat shock response is coordinated across the regulon and support the use of *HSP70* expression as a readout of heat shock response state at the nuclear level.

### Heat shock expression variation is associated with broader transcriptional programmes

We next asked whether variation in heat shock gene expression was associated with broader transcriptional differences. Nuclei were scored using a 15-gene heat shock response (HSR) UCell signature, and score group differential expression was examined separately for each annotated root cell type using all three biological replicates per condition. The workflow shown in Fig. 3E illustrates the 35 °C analysis. At 22 °C, the highest-scoring 5% of nuclei were designated ON and compared with the remaining 95% (OFF). At 35 °C, the lowest-scoring 5% were designated OFF and compared with the remaining upper 95% (ON). These asymmetric contrasts were chosen to capture the relatively high-scoring basal tail and the relatively low-scoring post-heat tail, respectively.

Higher HSR scores were associated with a recurrent heat response and protein folding programme across root cell types at both temperatures (Supplementary Datasets S1-S5). In control samples, 41 genes were upregulated in ON nuclei across at least 14 cell-types, including 28 which were not used to establish the UCell scores (which relied on *HSP70* and 14 *HSP70-*associated genes). After heat-stress, 63 significant differential genes were upregulated in ON nuclei across at least 14 cell-types, 50 of which were not found within the gene shortlist used to calculate UCell scores. Amongst these upregulated genes included *HSP70-2T*, *HSP70-15*, and other known HSPs, HSFs, or genes known to be heat-responsive.

By contrast, expression associated with the OFF group was more population specific (Fig. 3F–H). At 35 °C, genes relatively enriched in OFF nuclei in the combined Xylem & XPP population were enriched for defence-, jasmonate- and wound-response annotations. Jasmonate- and wound-response enrichment was also detected in meristem epidermis, but these programmes were not shared across all root cell types. The volcano plots illustrate gene-level differences associated in Xylem & XPP and non-hair epidermis (Fig. 3G,H). Together, these analyses show that variation in HSR activity is associated with a recurrent heat-response programme and with population specific differences in broader transcriptional programmes.

### Stochastic modelling and perturbation experiments identify regulators of heterogeneous heat shock gene induction

To test whether the observed transcriptional heterogeneity in the heat shock response could arise from the architecture of the underlying gene regulatory network, we developed a simplified stochastic model of the Arabidopsis heat shock response gene circuit (Fig. 4A, Fig. S15). The model focuses on a well characterised regulatory network linking heat induced protein misfolding to HSF activation and HSP70/HSP90 induction.

**Figure 4.**
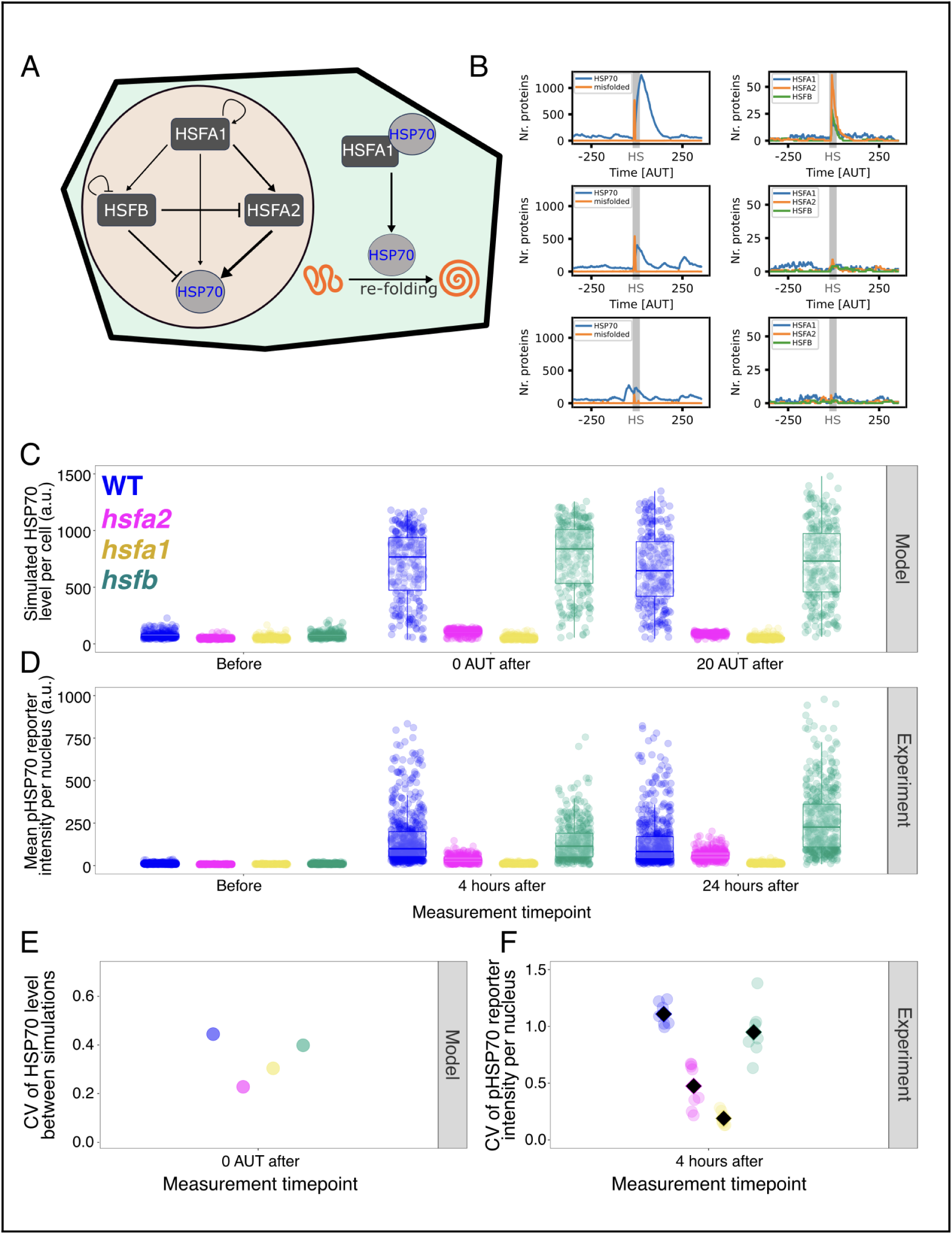
Stochastic modelling and perturbation experiments identify regulators of heterogeneous heat shock gene induction. **(A)** Schematic of the regulatory network captured in the model. Heat stress generates misfolded proteins, which are refolded by HSP70. HSP70 expression is regulated by a small network of heat shock transcription factors including HSFA1, HSFA2 and HSFB. **(B)** Representative single-cell stochastic simulations of the model under heat stress. Left, HSP70 and misfolded protein levels. Right, HSFA1, HSFA2 and HSFB levels. The grey shaded region indicates the heat stress period. Example trajectories illustrate heterogeneous HSP70 induction despite identical external input. **(C)** Model summary showing simulated HSP70 levels in WT and HSFA1-, HSFA2- or HSFB-perturbed simulations before heat stress, immediately after heat stress, and 20 arbitrary time units (AUT) after heat stress. HSFA1 perturbation strongly reduces simulated HSP70 induction, HSFA2 perturbation retains a lower and more compressed response, and HSFB perturbation largely preserves heat-induced HSP70 activation under these conditions. **(D)** Representative experimental measurements of nuclear *HSP70* reporter fluorescence in WT and HSF mutant backgrounds before heat stress, 4 h after heat stress, and 24 h after a 1 h heat stress. For compactness, labels indicate the perturbed HSF class: *hsfa1* denotes the *hsfa1abde* quadruple mutant background, and *hsfb* denotes the *hsfb1 hsfb2b* double mutant background. The experimental perturbations qualitatively match the model predictions: *hsfa1* strongly reduces reporter induction, *hsfa2* retains inducible but reduced reporter expression, and *hsfb* has a more limited effect. **(E)** Coefficient of variation (CV) of simulated HSP70 levels across cells in WT and HSFA1-, HSFA2- or HSFB-perturbed simulations immediately after heat stress. HSFA2 perturbation reduced simulated response heterogeneity, whereas HSFB perturbation had a more limited effect under these conditions. **(F)** Experimentally measured CV of nuclear *HSP70* reporter fluorescence across nuclei within individual roots 4 h after a 1 h heat shock. The experimental data showed the same qualitative pattern of altered response heterogeneity across HSF mutant backgrounds as the model.

The regulatory network comprises three heat shock transcription factors (HSFA1, HSFA2, and HSFB), chaperone proteins (HSP70/HSP90), and a pool of misfolded proteins (M) that accumulates under heat stress. In the absence of stress, HSFA1 is sequestered in the cytoplasm through binding to HSP70/HSP90 (7). Heat-induced protein misfolding competes for chaperone binding, releasing HSFA1 to activate expression of downstream targets including *HSP70* and *HSFA2* (13, 21, 22). HSFA2 further amplifies chaperone expression (14, 15, 25), while HSFB acts as a negative regulator (31).

We modelled gene expression using Hill functions and implemented the full dynamics using the Gillespie algorithm (32, 33). Each simulation tracks coarse-grained abundances of HSFA1, HSFA2, HSFB, HSP70/HSP90, folded and misfolded proteins, and associated protein complexes over time. Heat stress was simulated by transiently increasing the protein misfolding rate. Parameter values were chosen to reproduce the qualitative behaviour of the experimental system, including low basal HSP70/HSP90 output and heterogeneous induction after heat stress (see Methods).

Across 250 simulations, representing independent virtual cells, the model recapitulated key features of the data: under control conditions, HSP70 expression was low but exhibited cell-to-cell variability due to stochastic differences in regulatory state (Fig. 4B). Upon simulated heat stress, HSP70 was induced in most simulations, but with wide variation in peak levels and activation timing (Fig. 4B and Fig. S16), matching the heterogeneity observed in both live imaging and snRNA-seq (Fig. 1, Fig. 3 and Fig. S14).

We next used the model to ask how perturbing individual transcription factor components would alter *HSP70* induction. In silico perturbation of HSFA1, HSFA2 and HSFB produced distinct outcomes (Fig. 4C; Figs. S17–S19). Perturbing HSFA1 largely abolished simulated HSP70 induction, reducing expression to near-background levels, consistent with its role as an essential upstream activator of the heat shock response. Perturbing HSFA2 also reduced simulated HSP70 expression, but unlike HSFA1 perturbation, it retained a detectable response while narrowing the distribution of induced expression levels. Perturbing HSFB had a comparatively modest effect on *HSP70* expression and variability under these conditions (Fig. 4C,E; Figs. S17–S19).

We then compared these model predictions with *HSP70* reporter measurements in corresponding transcription factor knock-out backgrounds, using *35S::H2B-RFP* or, where this marker was silenced after crossing, *UBQ10::mScarlet-N7* as a constitutive nuclear marker for segmentation (Methods, Fig S21).

For simplicity, we refer to these backgrounds by the HSF class indicated in Fig. 4: *hsfa1* denotes the *hsfa1abde* quadruple mutant background. Sibling triple-mutant controls and an independent *hsfa1abde* family are shown in Fig. S25 and Fig. S26. *hsfa2* denotes the *hsfa2-1* mutant background, and *hsfb* denotes the *hsfb1-1 hsfb2b-1* double mutant background (15, 18, 19, 24, 31). Consistent with the model, the *hsfa1* background showed little reporter induction after heat stress, whereas the *hsfa2* background retained inducible reporter expression but showed reduced mean expression and lower cell-to-cell variability (Fig. 4D,F). As predicted, the *hsfb* background showed a more limited effect on reporter expression and variability (Fig. 4D,F, and S27). Thus, while *HSFA1* is required for heat shock induction, the transcription factor knock-out experiments revealed *HSFA2* as a candidate regulator of the breadth of *HSP70* responses across cells. We therefore examined the HSFA2-dependent response in more detail.

### HSFA2 tunes the magnitude and heterogeneity of *HSP70* induction

Because *HSFA2* emerged as a candidate regulator of *HSP70* response heterogeneity, we asked whether *HSFA2* and *HSP70* activity covary across individual cells during the heat shock response. snRNA-seq had already shown coordinated endogenous expression of *HSP70* and *HSFA2* across nuclei after 1 h of heat shock (Fig. 3C; Fig. S10). To test this relationship over time using an independent live-imaging reporter assay, we imaged transgenic Arabidopsis seedlings carrying *pHSP70::2xmTurq-N7* and *pHSFA2::YFP-N7* reporters together with the constitutive nuclear marker *35S::H2B-RFP* (Fig. S20A).

Seedlings were imaged before heat stress and again 4 h and 24 h after a 1 h heat stress at 35 °C, allowing us to follow the accumulation of both reporter signals during induction and recovery. Before heat stress, *HSP70*–*HSFA2* correlations were lower and more variable than after induction, consistent with low basal activity of both heat shock reporters and limited dynamic range. After heat shock, the two heat shock reporters became strongly correlated across individual nuclei, with Pearson correlation coefficients predominantly in the range R = 0.7–0.9 , whereas correlations involving the constitutive *35S::H2B-RFP* control remained weaker and less consistent (Pearson R typically ranges 0.003–0.5) (Fig. S20D–F). Together, these data show that the coordinated activation observed across the heat shock regulon includes co-activation of *HSFA2* and *HSP70* at both endogenous transcript and reporter levels.

We next examined how loss of *HSFA2* affects the distribution of *HSP70* responses across heat stress durations. Wild-type and *hsfa2-1* seedlings carrying the *HSP70* reporter were exposed to 35 °C for 30 min, 1 h or 2 h and imaged before heat stress and again 4 h and 24 h after treatment (Fig. 5A–D; Figs. S22, S23). The 1 h condition corresponds to the perturbation comparison introduced in Fig. 4, while the 30 min and 2 h treatments test the duration dependence of the *HSFA2* effect.

**Fig. 5.**
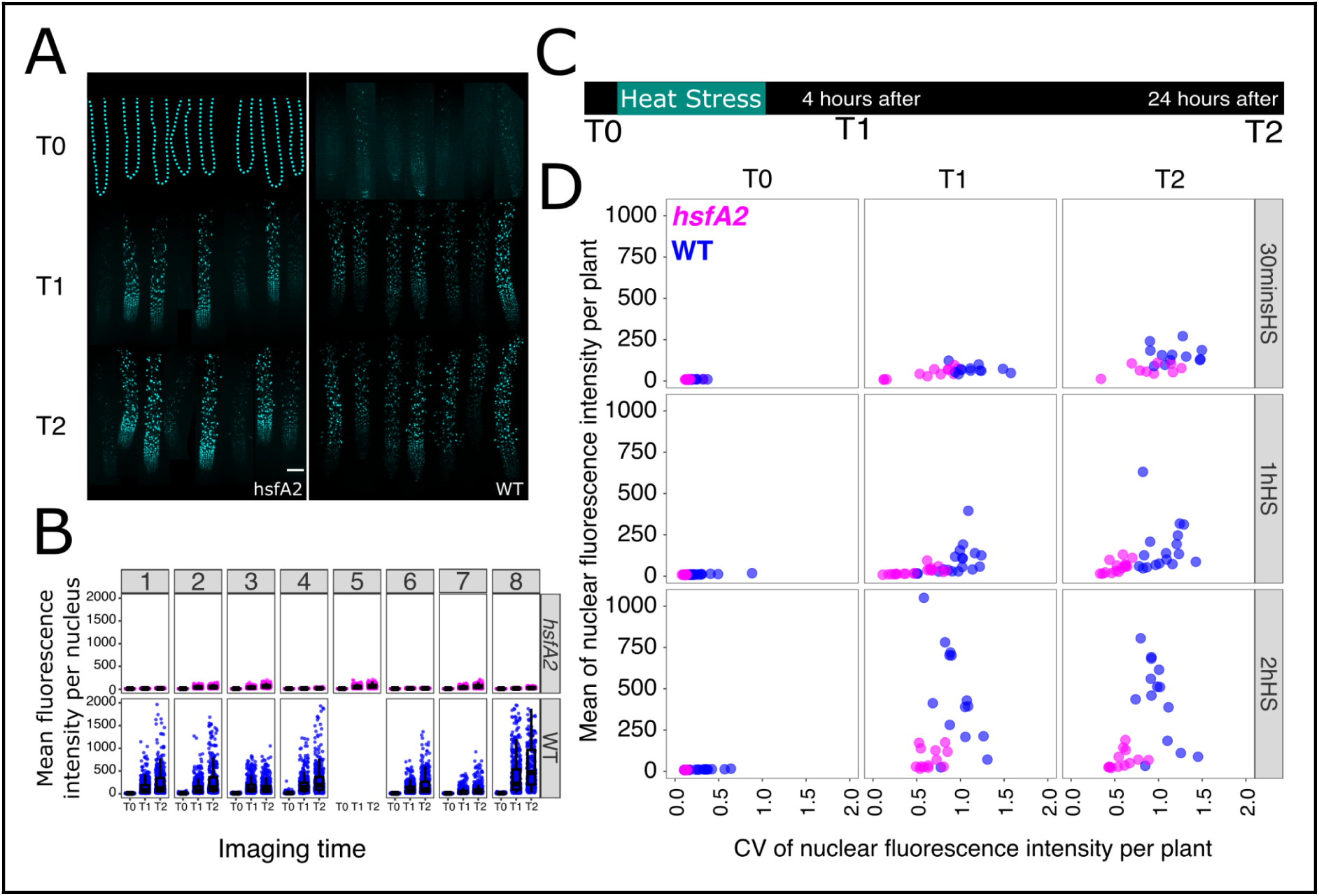
*HSFA2* tunes the magnitude and heterogeneity of *HSP70* reporter induction. **(A)** Representative confocal images of WT and *hsfa2* roots carrying the *HSP70* reporter before heat shock (T0), 4 h after heat shock (T1), and 24 h after heat shock (T2). Images show the 1 h heat stress condition. Basal *HSP70* reporter fluorescence is reduced in *hsfa2* compared with WT. Images are shown with adjusted display ranges for visualisation: *hsfa2*, 200–300; WT, T0, 200–250; WT, T1 and T2, 200–1200. **(B)** Quantification of nuclear mTurquoise2 fluorescence intensity in WT and *hsfa2* roots for the 1 h heat stress condition shown in A. Each dot represents one nucleus. **(C)** Imaging scheme. Seedlings were imaged before heat shock (T0), then re-imaged 4 h (T1) and 24 h (T2) after heat stress treatment. **(D)** Relationship between mean nuclear *HSP70* reporter fluorescence and coefficient of variation across nuclei within individual roots after 30 min, 1 h or 2 h heat stress. Each dot represents one root. *hsfa2* mutants show reduced reporter induction and, among roots with comparable mean activation, reduced expression heterogeneity. For D, data are pooled from matched WT and *hsfa2* imaging experiments. For the 30 min, 1 h and 2 h heat stress treatments, sample sizes were WT, n = 12, 19 and 15 seedlings, respectively, and *hsfa2*, n = 13, 20 and 11 seedlings, respectively, from 2, 3 and 2 independent imaging experiments. At the 24 h time point, the *hsfa2* sample size was n = 10 seedlings for the 30 min and 1 h treatments. The 1 h heat stress condition corresponds to the *hsfa2* data included in Fig. 4D,F. For the 30 min and 2 h comparisons in D, WT datasets are replotted from the experiments summarised in Fig. 1 and detailed in Figs. S4 and S5, respectively; the corresponding full *hsfa2* datasets are shown in Figs. S22 and S23. The 30 min WT and *hsfa2* datasets were acquired using the low-mTurq acquisition settings described in Figs. S4 and S22.

Consistent with our earlier observations, wild-type roots in these experiments showed broad distributions of reporter fluorescence across nuclei after heat stress. In *hsfa2* mutants, reporter induction was retained but reduced, with a narrower distribution of expression levels compared with wild type (Fig. 5A–D; Figs. S22, S23). Basal *HSP70* reporter activity was also reduced in *hsfa2* mutants, indicating that *HSFA2* contributes both to low-level pre-stress activity and to the breadth of stress-induced responses.

Because loss of HSFA2 also reduced mean reporter expression, we asked whether the difference in variability persisted among roots with comparable mean expression. For each root, we calculated mean nuclear fluorescence and the CV across nuclei. At each post heat imaging time, we restricted comparisons to overlapping mean expression ranges for the two genotypes within each experiment (see Methods; Fig. S24). The primary analysis pooled roots across the 30 min, 1 h and 2 h heat treatments and fitted log(CV) as a function of log (mean fluorescence), genotype, their interaction and experiment, separately at 4 h and 24 h after treatment.

Genotype significantly affected the relationship between mean expression and CV at both 4h (P = 3.9 x 10^-5) and 24h (P = 6.9 x 10^-5). At the centre of the shared expression range, *hsfa2* roots had an estimated 28% lower CV than WT at 4h (*hsfa2*/WT CV ratio = 0.72, 95% confidence intervals (CI) 0.62-0.83) and 35% lower CV at 24h (ratio = 0.65, 95% CI 0.56-0.76) (Supplementary Table S4). Thus, the reduced heterogeneity of the *hsfa2* response cannot be explained simply by comparison of roots with different mean reporter levels, supporting a role for HSFA2 in broadening the HSP70 expression response.

To ask whether altered heat responsive transcription more generally was sufficient to compress *HSP70* response heterogeneity, we examined a mechanistically distinct regulatory background using an *arp6* mutant. ARP6 is part of the SWR1 chromatin-remodelling complex required for normal deposition of the histone variant H2A.Z, which has been implicated in thermosensory, temperature-responsive and stress-responsive transcription (22, 34, 35). Loss of *ARP6* did not reproduce the reduced, compressed *HSP70* response observed in *hsfa2* mutants, and cell-to-cell heterogeneity remained evident after heat stress (Fig. S28). This suggests that the narrowing of the response distribution in *hsfa2* is not a general consequence of perturbing heat-responsive transcription, but reflects a more specific role for HSFA2-dependent amplification.

Together, these results show that *HSFA1* and *HSFA2* make distinct contributions to *HSP70* induction. Loss of *HSFA1* strongly compromised heat-induced reporter activation, whereas *hsfa2* mutants retained a reduced but inducible response with a compressed distribution across nuclei. Thus, HSFA2-dependent amplification provides a network mechanism for tuning both the magnitude and heterogeneity of heat shock induction across cells.

## Discussion

Gene expression variability is a pervasive feature of biological systems, but how it is regulated within intact multicellular tissues remains poorly understood. Here, we show that cell-to-cell heterogeneity in the Arabidopsis heat shock response is coordinated across the heat shock regulon and controlled by its regulatory architecture.

A key feature of this variability is that it reflects shared regulatory state within cells. Previous nascent-transcription imaging showed that heat shock reporters are activated heterogeneously across plant cells, with allele-specific stochasticity contributing strongly to variability in nascent reporter output over relatively short time scales (26). Our dual-reporter analysis tested how this heterogeneity appears in accumulated *HSP70* reporter output after heat stress. The strong correlation between independently integrated *HSP70* reporters indicates that sustained cell-to-cell differences are dominated by shared upstream regulatory variation. snRNA-seq provided a complementary view of the endogenous response, showing that heat shock genes covary across nuclei after stress. Together, these findings link *HSP70* heterogeneity to coordinated regulon activation.

Our single-nucleus analysis further linked variation in heat shock gene expression to broader transcriptional differences within cell types (Fig. 3; Datasets S1-S5). After heat stress, lower heat shock gene expression was associated with higher expression of genes linked to defence, jasmonate and wound responses in some cell types. Cells with weaker heat shock responses may therefore be engaged in other stress-response programmes, although the functional consequences of these differences remain to be established. Single-cell transcriptomics has also identified an environmentally regulated transcriptional state shared across Arabidopsis root cell types (36).

Stochastic modelling provided a mechanistic framework for interpreting how coordinated heterogeneity can arise (Fig. 4). In silico perturbations predicted that loss of HSFA1 would largely collapse HSP70 induction, whereas loss of HSFA2 would retain inducibility but compress the distribution of responses. Experimental perturbations were consistent with these predictions (Figs. 4 and 5, Figs. S22-27), identifying HSFA2-dependent amplification, rather than HSFB-mediated repression, as a key component controlling response heterogeneity. Thus, within the broader heat shock regulatory network, heterogeneous induction is shaped by a coherent feed-forward-like amplification branch, in which HSFA1 activates HSP70 directly and indirectly through HSFA2, embedded within negative feedback from HSP70/HSP90-mediated sequestration of HSFA1.

These findings suggest that engineering heat resilience may depend on controlling how stress responses are distributed across cells. Heat stress is a major threat to crop productivity, and HSF–HSP/chaperone networks are central targets for improving thermotolerance in plants (37). However, constitutive or high-level activation of heat shock regulators can also cause growth defects, highlighting a potential cost of uniformly activating the response (23). In microbial systems, gene-expression noise can support bet-hedging by allowing subpopulations to enter different physiological states before stress occurs (38–42). In plant tissues, where cells are fixed within organs, the different transcriptional programmes associated with heat shock activity (Fig. 3F–H) raise the possibility that neighbouring cells maintain different protective states during stress without requiring uniform activation across the tissue. Cells that do not strongly activate *HSP70* may still be protected by basal chaperone pools, alternative protective pathways, or proteins produced in neighbouring cells, including through the movement of HSP70-family chaperones (43, 44). Future experiments combining functional HSP70 translational reporters with measurements of cell survival and root growth after heat stress could test how expression heterogeneity relates to cellular recovery and tissue-level thermotolerance.

Tissue-level responses in plants can involve changes in the distribution of gene expression across individual cells (26, 45–48). Our results identify HSFA2-dependent amplification as a mechanism for tuning both the magnitude and breadth of HSP70 induction across cells, showing that response heterogeneity is a tuneable feature of the plant heat shock response.

## Methods

Plant growth, heat treatments, and imaging are described below. Detailed methods for transgenic-line generation, statistical analysis, single-nucleus RNA-sequencing library preparation and analysis, as well as stochastic modelling are provided in the SI Appendix.

## Plant Growth conditions

Arabidopsis seeds were sterilised by immersing them in 5% bleach for 3 minutes followed by immersion in 70% ethanol for 3 minutes. Seeds were washed with autoclaved water 5-6 times after this. Sterilised seeds were encircled in a droplet of water and were placed evenly on solid ½ Murashige and Skoog (MS) plates. The plates were wrapped in aluminium foil and incubated for 2-3 days at 4°C for stratification. The plates were transferred to Conviron growth chambers with temperature set at 22°C, 65% relative humidity, light intensity of 170 μmoles under 12/12 light dark cycle. Seeds were collected from a single parent for every imaging experiment, repeats have been performed from both the same and different seed bags. To propagate plants for transformation, crossing and bulking P40 trays with F2 (Levington) soil were used. The plants were grown in a Conviron growth chamber with temperature set at 21°C at day time and 17°C at night, 65% relative humidity, light intensity of 170 μmoles under 16/8 light dark cycle.

## Chamber preparation

Small square coverslips were pressed on frosted glass slides with the help of low viscosity non toxic grease to place the whole plant in the same plane and avoid compression of the leaves. A rectangular shape made of low viscosity non toxic grease was made on a glass frosted slide (Epredia) 300-500µL of liquid ½ Murashige and Skoog (MS) media was pipetted on top. The grease barrier was used to hold the liquid in place. 5-6 days old Arabidopsis seedlings were picked up with forceps exerting minimal force (10-14 days for T1-generation experiments where the plants grew with antibiotics selection) and the roots were placed on the square 22X22 mm coverslip. More liquid ½ Murashige and Skoog (MS) media was pipetted on top of the plants according to need. A rectangular 24X50 mm coverslip (Epredia) was placed over gently to seal the chamber.

## Heat Stress treatment

A water bath was heated to 35 °C and the temperature was monitored to ensure a stable stress condition. The slides containing the plants were enclosed in a round plastic petri dish, sealed with parafilm and placed into the waterbath. When the petri dish was taken out of the waterbath it was then placed in a growth chamber with settings matching the normal growth conditions. For mutant comparisons, each independent experiment included matched WT controls, and heat stress treatments were applied simultaneously to all genotypes included in that experiment.

## Confocal microscopy

Confocal imaging was performed using either an inverted Nikon AXR confocal microscope or an upright Zeiss 880 confocal microscope. Seedlings were imaged before heat stress and then re-imaged 4 h and 24 h after the end of heat treatment. Z-stacks were acquired from the first visible optical plane to a depth of 80–140 µm, with a 1 µm step size. Images were acquired at 12-bit depth on the Nikon AXR and 16-bit depth on the Zeiss 880. On the Nikon AXR, fast resonance scanning was used to reduce acquisition time.

For Nikon AXR imaging, excitation wavelengths were 448 nm for mTurquoise2, 514 nm for YFP, and 552 nm for RFP/mScarlet. For Zeiss 880 imaging, mTurquoise2 and RFP were excited using 440 nm and 561 nm laser lines, respectively. In two-channel experiments, emission was collected at 463–541 nm for mTurquoise2 and 581–634 nm for RFP/mScarlet on the Nikon AXR, and at 460–525 nm and 561–624 nm on the Zeiss 880. In three-channel experiments on the Nikon, emission was collected at 463–501 nm for mTurquoise2, 524–544 nm for YFP, and 575–625 nm for RFP using GaAsP detectors. Images were acquired using a 20× Differential Interference Contrast Plan Apo VC objective, NA 0.75, on the Nikon AXR, and a 20× Plan-Apochromat objective, NA 0.8, on the Zeiss 880.

Roots were excluded from analysis if the constitutive nuclear red fluorescent marker was lost at any imaging time point. In analysable roots, this marker was visible as a distinct nuclear signal throughout the imaged root region. Excluded roots showed near complete loss of nuclear red fluorescence, with only faint diffuse residual fluorescence and no discernible nuclei, consistent with loss of tissue viability. This occurred in 45 of 430 roots analysed.

## Image Analysis

Images were processed in Fiji (49) using the FRETENATOR2 plugin (50–52). Confocal Images were segmented using the RFP/mScarlet channel with FRETENATOR segment and ratio v2.0 to segment the recorded nuclei without bias. The settings were kept constant for each experiment after optimising for best segmentation. Otsu thresholding was used, and the difference-of-Gaussian filter sizes were determined empirically for each experiment. A minimum ROI size was set between 20 to 50 to strictly ensure nuclear segmentation.

## Acknowledgements

We thank Dr Fritz Schöffl and Dr Yee-yung Charng for kindly providing us the *hsfa1abde*, *hsfa1ab*, and *hsfa1de* mutant lines and Dr Nadia Radzman for kindly sharing the *UBQ10::mScarlet-N7* construct. We thank Dr Raymond Wightman and Gareth Evans for help with microscopy, and Dr James Rowe for the help with FRETENATOR analysis. ChatGPT (OpenAI, GPT-5) was used for coding assistance and language editing. This work was funded by the BBSRC grant award no. BB/V006088/1, as well as by the Gatsby Charitable Foundation.

## Competing Interest Statement

Authors have no competing interests

## Supporting Information

Additional supporting information is available (see the Supporting Information file and

Supplementary Datasets) including:

Supplemental Methods Figures S1 to S28 Tables S1 to S5

Legends for Datasets S1 to S5 SI References

Supplementary Datasets S1 to S5

## Notes

### Competing Interest Statement

The authors have declared no competing interest.

## References

1. I. S. Araújo, et al., Stochastic gene expression in Arabidopsis thaliana. Nature Communications 8, 2132 (2017).

2. S. Kong, B. Rusnak, M. Zhu, A. H. K. Roeder, Stochastic gene expression in auxin signaling in the floral meristem of Arabidopsis thaliana. Nat Commun 16, 4682 (2025).

3. C. Maranas, S. VanGilder, L. Nguyen, J. Nemhauser, Auxin promotes robust founder cell specification during Arabidopsis lateral root initiation. Genetics iyag120 (2026). 10.1093/genetics/iyag120.

4. Q. Fan, D. Jespersen, Proteases and the ubiquitin-proteasome system: Understanding protein degradation under heat stress in plants. Environmental and Experimental Botany 237, 106174 (2025).

5. X. Li, et al., Stress granules sequester autophagy proteins to facilitate plant recovery from heat stress. Nat Commun 15, 10910 (2024).

6. N. Ohama, H. Sato, K. Shinozaki, K. Yamaguchi-Shinozaki, Transcriptional Regulatory Network of Plant Heat Stress Response. Trends in Plant Science 22, 53–65 (2017).

7. K. Yamada, et al., Cytosolic HSP90 regulates the heat shock response that is responsible for heat acclimation in Arabidopsis thaliana. J Biol Chem 282, 37794–37804 (2007).

8. E. Vierling, The Roles of Heat Shock Proteins in Plants. Annual Review of Plant Physiology and Plant Molecular Biology 42, 579–620 (1991).

9. F. McLoughlin, et al., Class I and II small heat-shock proteins protect protein translation factors during heat stress. Plant Physiol. 172, 1221–1236 (2016).

10. E. Llamas, et al., The intrinsic chaperone network of Arabidopsis stem cells confers protection against proteotoxic stress. Aging Cell 20, e13446 (2021).

11. R. S. Boston, P. V. Viitanen, E. Vierling, Molecular chaperones and protein folding in plants. Plant Mol Biol 32, 191–222 (1996).

12. M. P. Mayer, B. Bukau, Hsp70 chaperones: Cellular functions and molecular mechanism. Cell Mol Life Sci 62, 670 (2005).

13. A. Nishizawa-Yokoi, et al., HsfA1d and HsfA1e Involved in the Transcriptional Regulation of HsfA2 Function as Key Regulators for the Hsf Signaling Network in Response to Environmental Stress. Plant and Cell Physiology 52, 933–945 (2011).

14. J. Lämke, K. Brzezinka, I. Bäurle, HSFA2 orchestrates transcriptional dynamics after heat stress in Arabidopsis thaliana. Transcription 7, 111–114 (2016).

15. F. Schramm, et al., The Heat Stress Transcription Factor HsfA2 Serves as a Regulatory Amplifier of a Subset of Genes in the Heat Stress Response in Arabidopsis. Plant Mol Biol 60, 759–772 (2006).

16. N. Andrási, A. Pettkó-Szandtner, L. Szabados, Diversity of plant heat shock factors: regulation, interactions, and functions. Journal of Experimental Botany 72, 1558–1575 (2021).

17. A. Guihur, M. E. Rebeaud, P. Goloubinoff, How do plants feel the heat and survive? Trends in Biochemical Sciences 47, 824–838 (2022).

18. H.-C. Liu, H.-T. Liao, Y.-Y. Charng, The role of class A1 heat shock factors (HSFA1s) in response to heat and other stresses in Arabidopsis. *Plant*, Cell & Environment 34, 738–751 (2011).

19. T. Yoshida, et al., Arabidopsis HsfA1 transcription factors function as the main positive regulators in heat shock-responsive gene expression. Mol Genet Genomics 286, 321–332 (2011).

20. N. Ohama, et al., The Transcriptional Cascade in the Heat Stress Response of Arabidopsis Is Strictly Regulated at the Level of Transcription Factor Expression. Plant Cell 28, 181–201 (2016).

21. W. Busch, M. Wunderlich, F. Schöffl, Identification of novel heat shock factor-dependent genes and biochemical pathways in Arabidopsis thaliana. The Plant Journal 41, 1–14 (2005).

22. S. Cortijo, et al., Transcriptional Regulation of the Ambient Temperature Response by H2A.Z Nucleosomes and HSF1 Transcription Factors in Arabidopsis. Molecular Plant 10, 1258–1273 (2017).

23. D. Ogawa, K. Yamaguchi, T. Nishiuchi, High-level overexpression of the Arabidopsis HsfA2 gene confers not only increased thermotolerance but also salt/osmotic stress tolerance and enhanced callus growth. Journal of Experimental Botany 58, 3373–3383 (2007).

24. Y. Charng, et al., A Heat-Inducible Transcription Factor, HsfA2, Is Required for Extension of Acquired Thermotolerance in Arabidopsis. Plant Physiol 143, 251–262 (2007).

25. A. Nishizawa, et al., Arabidopsis heat shock transcription factor A2 (HSFA2) as a key regulator in response to several types of environmental stress. The Plant Journal 48, 535–547 (2006).

26. S. Alamos, A. Reimer, K. K. Niyogi, H. G. Garcia, Quantitative imaging of RNA polymerase II activity in plants reveals the single-cell basis of tissue-wide transcriptional dynamics. Nat. Plants 7, 1037–1049 (2021).

27. S. Hani, et al., Live single-cell transcriptional dynamics via RNA labelling during the phosphate response in plants. Nat. Plants 7, 1050–1064 (2021).

28. K. Jean-Baptiste, et al., Dynamics of Gene Expression in Single Root Cells of Arabidopsis thaliana. Plant Cell 31, 993–1011 (2019).

29. F. Federici, L. Dupuy, L. Laplaze, M. Heisler, J. Haseloff, Integrated genetic and computation methods for in planta cytometry. Nat Methods 9, 483–485 (2012).

30. J. Goedhart, et al., Structure-guided evolution of cyan fluorescent proteins towards a quantum yield of 93%. Nat Commun 3, 751 (2012).

31. M. Ikeda, N. Mitsuda, M. Ohme-Takagi, Arabidopsis HsfB1 and HsfB2b Act as Repressors of the Expression of Heat-Inducible Hsfs But Positively Regulate the Acquired Thermotolerance. Plant Physiol 157, 1243–1254 (2011).

32. D. T. Gillespie, Exact stochastic simulation of coupled chemical reactions. J. Phys. Chem. 81, 2340–2361 (1977).

33. D. T. Gillespie, Stochastic Simulation of Chemical Kinetics. Annu. Rev. Phys. Chem. 58, 35–55 (2007).

34. S. V. Kumar, P. A. Wigge, H2A.Z-containing nucleosomes mediate the thermosensory response in Arabidopsis. Cell 140, 136–147 (2010).

35. W. Sura, et al., Dual Role of the Histone Variant H2A.Z in Transcriptional Regulation of Stress-Response Genes. Plant Cell 29, 791–807 (2017).

36. M. Oliva, et al., An environmentally regulated dichotomy in cell lineages of the root. Nature 1–10 (2026). 10.1038/s41586-026-11179-y.

37. S. Fragkostefanakis, S. Röth, E. Schleiff, K.-D. Scharf, Prospects of engineering thermotolerance in crops through modulation of heat stress transcription factor and heat shock protein networks. *Plant*, Cell & Environment 38, 1881–1895 (2015).

38. O. Patange, et al., Escherichia coli can survive stress by noisy growth modulation. Nat Commun 9, 5333 (2018).

39. N. Q. Balaban, Bacterial Persistence as a Phenotypic Switch. Science 305, 1622–1625 (2004).

40. J.-W. Veening, W. K. Smits, O. P. Kuipers, Bistability, Epigenetics, and Bet-Hedging in Bacteria. Annu. Rev. Microbiol. 62, 193–210 (2008).

41. B. M. Martins, J. C. Locke, Microbial individuality: how single-cell heterogeneity enables population level strategies. Current Opinion in Microbiology 24, 104–112 (2015).

42. D. Fraser, M. Kærn, A chance at survival: gene expression noise and phenotypic diversification strategies. Molecular Microbiology 71, 1333–1340 (2009).

43. K. Aoki, F. Kragler, B. Xoconostle-Cázares, W. J. Lucas, A subclass of plant heat shock cognate 70 chaperones carries a motif that facilitates trafficking through plasmodesmata. Proceedings of the National Academy of Sciences 99, 16342–16347 (2002).

44. L. Yang, et al., Noncell-autonomous HSC70.1 chaperone displays homeostatic feedback regulation by binding its own mRNA. New Phytologist 237, 2404–2421 (2023).

45. G. M. Turco, et al., Molecular Mechanisms Driving Switch Behavior in Xylem Cell Differentiation. Cell Reports 28, 342–351.e4 (2019).

46. A. Angel, J. Song, C. Dean, M. Howard, A Polycomb-based switch underlying quantitative epigenetic memory. Nature 476, 105–108 (2011).

47. A. Angel, et al., Vernalizing cold is registered digitally at FLC. Proc Natl Acad Sci U S A 112, 4146–4151 (2015).

48. S. Berry, C. Dean, Environmental perception and epigenetic memory: mechanistic insight through FLC. The Plant Journal 83, 133–148 (2015).

49. J. Schindelin, et al., Fiji: an open-source platform for biological-image analysis. Nat Methods 9, 676–682 (2012).

50. B. Tang, et al., SALICYLIC ACID SENSOR1 reveals the propagation of an SA hormone surge during plant pathogen advance. Science 390, 188–194 (2025).

51. J. Rowe, et al., Next-generation ABACUS biosensors reveal cellular ABA dynamics driving root growth at low aerial humidity. Nat. Plants 9, 1103–1115 (2023).

52. J. H. Rowe, A. Rizza, A. M. Jones, “Quantifying Phytohormones in Vivo with FRET Biosensors and the FRETENATOR Analysis Toolset” in Environmental Responses in Plants: Methods and Protocols, P. Duque, D. Szakonyi, Eds. (Springer US, 2022), pp. 239–253.

