## Supplementary Information for "Regulatory architecture tunes gene expression noise in the plant heat shock response"

James Locke

##### **This file includes:**

**Supplemental Methods**

**Figures S1 to S28**

**Tables S1 to S5**

**Legends for Datasets S1 to S5**

**SI References**

##### **Other supporting materials for this manuscript include the following:**

**Supplementary Datasets S1 to S5**

### Supplemental Methods

#### 1. Generation of transgenic lines and mutant reporter backgrounds

##### **Reporter Constructs:**

The *pHSP70::2xmTurq-N7* transcriptional reporter line was generated in the Locke lab. The construct consists of 2104 bp of the promoter of *HSP70* (*AT3G12580*), a member of the Arabidopsis *HSP70* gene family, fused to double, nuclear localised mTurquoise fluorescent protein (2xmTurq-N7) followed by the *Agrobacterium tumefaciens* nopaline synthase gene terminator (NosT). The transcriptional fusion construct was generated using the Gateway cloning system (Invitrogen). The entry clones were designed and constructed for all three fragments in VectorNTI. The primers with the attachment site for BP-clonase reaction were also designed for three inserts in VectorNTI. The *attB* sites were added to each fragment by performing PCR (the primer sequences for the *HSP70* promoter are described in Supplementary Table S1). mTurquoise and NosT were cloned into pDONRp1-p2R and pDONRp2R-p3, respectively. The presence of the fragments and their correct direction were confirmed by Sanger sequencing of each vector. After that, LR reaction was performed using LR Clonase II Plus enzyme into the pB7M34GW destination vector (Basta-resistance) to fuse all three fragments in the expression clone.

*pHSP70::1xYFP-N7*, *pHSFA2::1xYFP-N7* transcriptional reporters constructs were prepared following the same method as *pHSP70::2xmTurq-N7* using the primers described in Supplementary Table S1.

##### **Reporter line generation:**

The *pHSP70::2xmTurq-N7* construct was transformed into a homozygous *35S::H2B-RFP* line in the Col-0 background (1) and positive transformants were screened for using BASTA selection. T2 segregation analysis was performed to find lines with a single locus of insertion and homozygous T4 lines were obtained. qPCR-based copy-number analysis performed by idnaGenetics, Norwich, UK, indicated that line *pHSP70::2xmTurq-N7* 1-1 contains approximately nine tandem copies of the transgene at the insertion locus, whereas line 3-14 contains approximately three copies. *pHSP70::1xYFP-N7*, *pHSFA2::1xYFP-N7*, transcriptional reporters constructs were transformed into the *pHSP70::2xmTurq-N7* transcriptional reporter line.

##### **Crosses into mutant backgrounds**

***hsfa2-1 pHSP70::2xmTurq-N7 UBQ10::mScarlet-N7***

*hsfa2-1* (SALK\_008978C) mutant seeds (2) were obtained from Nottingham Arabidopsis Stock Centre (NASC) and homozygous mutants (confirmed by PCR-based genotyping) were crossed with *pHSP70::2xmTurq-N7 35S::H2B-RFP*. We screened an F2 derived from selfed F1 plants on BASTA to select for plants carrying the *pHSP70::2xmTurq-N7* construct and genotyped F2 plants to find homozygous *hsfa2-1* mutants (see Supplementary Table S2 for primers). We screened the F3 seeds of these plants on BASTA to obtain lines that were homozygous for *pHSP70::2xmTurq-N7*. We observed silencing in the expression of the *35S::H2B-RFP* and therefore F4 plants were transformed with *UBQ10::mScarlet-N7* and T2 seeds containing the red nuclear marker were used for imaging.

##### ***hsfb2b-1 hsfb1-1 pHSP70::2xmTurq-N7 UBQ10::mScarlet-N7***

*hsfb2b-1* (SALK\_047291) and *hsfb1-1* (SALK\_104713C) mutant lines were obtained from NASC. SALK\_047291 contains 3 T-DNA insertions in separate loci, one of which disrupts *HSFB2b*. We genotyped plants that were segregating for the 3 T-DNA insertions to find individuals that were homozygous wild-type for the two off-target insertions and homozygous mutant for *hsfb2b-1*. We then crossed one of these plants with *hsfb1-1* mutants.

We screened an F2 derived from selfed F1 plants and obtained the homozygous *hsfb2b-1 hsfb1-1* double mutant (3) which we used for crossing with the *pHSP70::2xmTurq-N7* marker line. Because silencing of the *35S::H2B-RFP* marker had been observed in crosses involving SALK lines, we used plants carrying *UBQ10::mScarlet-N7* as an alternative red nuclear marker. Specifically, we crossed the *hsfb2b-1 hsfb1-1* double mutant with *pHSP70::2xmTurq-N7 35S::H2B-RFP* T1 plants that had been transformed with the *UBQ10::mScarlet-N7* construct. In the F1 of this cross, we screened plants on Hygromycin plates to select for the presence of the *UBQ10::mScarlet-N7* construct. We sowed an F2 derived from the selfed F1 plants on BASTA to screen for *pHSP70::2xmTurq-N7*, visually selected plants carrying the *UBQ10::mScarlet-N7* construct using a Leica stereofluorescence microscope and then performed PCR-based genotyping to find a *hsfb2b-1 hsfb1-1* double mutant carrying the *pHSP70::2xmTurq-N7*. We sowed the F3 of this homozygous *hsfb2b-1 hsfb1-1* mutant plant on BASTA and Hyg to select for *pHSP70::2xmTurq-N7* and *UBQ10::mScarlet-N7* respectively. Survivors carrying both transgenes were then selfed and screened in the F4 for homozygosity of the transgenes. Experiments were performed on F4 plants that were determined to be homozygous for both markers.

##### ***hsfa1abde pHSP70::2xmTurq-N7 UBQ10::mScarlet-N7***

The *hsfa1abde* mutant is in a mixed Ws/Col-0 background (4). To account for possible background effects on the expression of *pHSP70::2xmTurq-N7*, we obtained both quadruple mutants and triple mutant comparison lines from segregating F3 or F4 families derived from individual F2 plants. We did this for families derived from two different F2 plants (Figure S26).

Family 43 was obtained from F2 plant 43, which was homozygous for *pHSP70::2xmTurq-N7* (determined by screening the F3 on BASTA plates and finding all seedlings survived) and was *hsfa1a* +/- *bde* (i.e. homozygous mutant for *hsfa1 b d* and *e* genes and heterozygous for *hsfa1a*). In the F3 of this family, through PCR-based genotyping, we obtained plants that were homozygous *hsfa1abde* mutants and those that were homozygous *hsfa1bde* mutants and used F4 seeds from these plants to compare *pHSP70::2xmTurq-N7* gene expression in lines with a similar Col/Ws background but differing in the presence of the WT *HSFA1a* gene (Figure S25, Figure S26 *hsfa1abde* 43 vs *hsfa1 bde* 43). The *hsfa1abde* mutant from this family was used in Figure 4.

Family 176 was descended from an F2 plant with the genotype *hsfa1ab d* +/- *e* +/- . We screened the F3 on BASTA plates and found the line was homozygous for *pHSP70::2xmTurq-N7*. In the F3, we obtained a plant that was *hsfa1abd e* +/- . In an F4 family derived from this plant, we obtained plants that were homozygous *hsfa1abde* and homozygous *hsfa1abd* mutants, and used these for further analyses. These lines were used as an independent family to test whether reduced *pHSP70::2xmTurq-N7* induction in *hsfa1abde* was reproducible and not explained by family-specific background effects. (Figure S26 *hsfa1 abde* 176 versus *hsfa1 abd* 176).

Expression of the *35S::H2B-RFP* nuclear marker was weak and patchy in F2 and later generations of the *hsfa1* crosses, consistent with silencing of the 35S promoter. This could be because the *hsfa1 d* and *e* mutant alleles are SALK lines and contain a 35S promoter on the T-DNA which is reported to cause silencing of other 35S promoters in the same background (5). We thus transformed the homozygous mutants obtained with *UBQ10::mScarlet-N7* and used T2 seeds from this transformation containing the red nuclear marker for imaging. Experiments were performed using F6 plants for lines descended from F2 plant 43 line and F7 plants for lines descended from F2 plant 176.

#### ***arp6-2 pHSP70::2xmTurq-N7 35S::H2B-RFP***

*arp6-2* (SAIL\_236\_C07) mutants (6) were obtained from NASC and crossed with WT *pHSP70::2xmTurq-N7 35S::H2B-RFP* plants. In an F2 derived from selfed F1 plants, we noticed an aberrant flower phenotype with an almost complete absence of petals that segregated independently of the T-DNA insertion in *arp6*, as assessed by PCR-based genotyping (see Supplementary Table S2 for primers). We thus selected for plants that were homozygous *arp6* mutants and lacked the aberrant flower phenotype. We screened the F3 seeds of these plants using a Leica stereofluorescence microscope to identify F3 families that were homozygous for *pHSP70::2xmTurq-N7*, and carrying the *35S::H2B-RFP* construct and we selected those families where none of the plants had the aberrant flower phenotype, consistent with removal of the unlinked mutation causing that phenotype. We used the F4 seeds from these families for further analysis. The *35S::H2B-RFP* was not silenced in this background.

### 2. Statistical Analysis

#### Quantification of confocal cell-to-cell variability

For imaging analyses, nuclear fluorescence measurements were summarised separately for each root at each imaging time point. Mean fluorescence and the sample standard deviation were calculated across segmented nuclei within each root, and cell-to-cell variability was quantified as the coefficient of variation (CV = SD/mean). Each root therefore contributed one value for mean fluorescence and one for CV at each time point. Fold changes in mean fluorescence were calculated relative to the pre-stress value for the same seedling. Correlations between fluorescent reporters were quantified using Pearson correlation coefficients calculated across nuclei within individual roots at each imaging time point. Sample sizes and numbers of independent imaging experiments are provided in the corresponding figure legends.

#### Mean-adjusted comparison of HSP70 reporter variability between WT and *hsfa2*

Because CV depends on mean expression and *hsfa2* altered mean reporter fluorescence, mean adjusted variability was analysed over expression ranges represented by both genotypes. For each experiment and imaging time point, the analysis interval was defined as the intersection of the observed ranges of root mean fluorescence in WT and *hsfa2*. Roots outside this interval were excluded from the mean adjusted comparison, and experiments lacking within experiment overlap did not contribute to that time point.

The primary analyses were performed separately for the 4 hours (T1) and 24 hours after heat stress (T2) imaging time points, pooling roots across the 30 min, 1 h and 2 h heat-shock treatments. Linear models of the form

$$\log(\text{CV}) \sim \log(\text{mean fluorescence}) \times \text{genotype} + \text{experiment}$$

were fitted to the retained roots. Log mean fluorescence was centred on the mean of the retained observations. WT was used as the reference genotype and experiment was included as a fixed blocking factor. Because each experiment corresponded to a single heat-shock duration, experiment also accounted for treatment-specific experimental offsets. The genotype-by-mean interaction allowed the relationship between mean expression and variability to differ between genotypes. The genotype effect was tested by comparing the model above with a model containing only mean fluorescence and experiment (Supplementary Table S4).

#### 3. Single Nuclei Isolation and RNA sequencing

##### Preparation of nuclei and 10X snRNA-seq library preparation

Plants were grown vertically under neutral day (12 h light, 12 h dark) at 22°C in a growth chamber (Panasonic). One and a half hours after dawn, at least 200 11-day-old *Arabidopsis thaliana* (Col-0) seedlings were collected and placed in a microscope slide. To avoid root breakage and dryness, roots were saturated with liquid ½ MS and covered with a coverslip. Each slide was then placed inside a round petri dish and wrapped with parafilm to avoid water leaking into the sample. Petri dishes for heat shock were placed in a water bath at 35°C and control samples in a water bath at 22°C.

After 55 minutes, slides were removed from the water baths and carefully removed from the coverslips. For fresh samples roots were excised from cotyledons with a razorblade and chopped to very small pieces for immediate nuclei extraction. For frozen samples, roots were excised from cotyledons and snap frozen in liquid nitrogen and kept at -70°C until extraction.

To each sample was added 3 mL of Honda buffer (2.5% Ficoll 400, 5% Dextran T40, 0.4 M sucrose, 10 mM MgCl<sub>2</sub>, 1 µM DTT, 0.5% Triton X-100, 1 tablet/50 mL cComplete Protease Inhibitor Cocktail (Roche), 0.4 U µL<sup>-1</sup> RiboLock (Thermo Fisher Scientific), 25 mM Tris-HCl, pH 7.4). The contents were then transferred to a 70-µm mesh filter on top of a 5 cm petri dish using wide bore filtered pipette tips (ART, Thermo Fisher Scientific). With a sterile micropestle, the contents that remained inside the filter were homogenised and washed with Honda to a final volume of 6 mL. Petri dishes were then placed in an orbital shaker at 4°C 100 rpm for exactly 15 minutes, to release nuclei. The resulting suspension was filtered through a 70-µm, 35-µm and a 20-µm strainer, and centrifuged at 1000 *g* for 6 min at 4°C. The pellet was resuspended carefully in 500 µL staining buffer (12 µM PI, 0.4 U µL<sup>-1</sup> Ambion RNase Inhibitor (Thermo Fisher Scientific), 0.2 U µL<sup>-1</sup> SUPERaseIn RNase Inhibitor (Thermo Fisher Scientific) in PBS). Nuclei were sorted by gating on the PI peaks using a BD Influx System (BD Biosciences) with a 100 µm nozzle at 20 psi pressure. 15,000 - 50,000 nuclei per sample were collected into 200 µL of landing buffer (4% Ultra-pure BSA in PBS, 2 U µL<sup>-1</sup> Ambion RNase Inhibitor, 1 U µL<sup>-1</sup> SUPERaseIn RNase Inhibitor) and centrifuged at 1500 *g* for 6 min at 4°C. The nuclear pellet was resuspended in 43 µL of landing buffer and loaded to a 10X Chromium chip for library preparation using 10x Single Cell 3' v4 (GEM-X) reagents according to the manufacturer's instructions. Libraries were sequenced using Illumina Novaseq X 25B.

##### snRNA-seq data analysis

Fastq files were processed with Cell Ranger v5 (7) mapping to the *A. thaliana* genome using the Araport11 annotation (8). Quality control was performed for each sample, where nuclei were considered viable if their gene feature number > 500 and their percentage of mitochondrial gene expression < 5%. Adaptive thresholds were used for maximum read counts and chloroplast gene expression percentages across samples. Genes expressed in < 0.5% of nuclei were removed. DoubletFinder v2.0.6 (9) was used to detect and remove multiplet nuclei which were estimated based on the expected multiplet rates provided by the Chromium Single Cell 3' Reagent Kits User Guide. Mitochondrial and chloroplast genes were also removed prior to downstream analyses.

The final number of nuclei that passed quality control were 3886, 5179 and 11340 for 22°C samples, and 8261, 6853, and 10292 for 35°C samples, for replicate 1 (frozen), 2 and 3 (fresh) respectively.

Seurat v5.3.1 (10) was used to cluster nuclei and assign cell-type annotations across an integrated transcriptomic map including both control and heat-stress nuclei. We first removed 75 of the top heat-responsive genes determined by *Blair et al.* (2019) (11) in bulk *A. thaliana* seedlings from the remaining gene feature list to reduce the contribution of heat-stress variation. Nuclei expression were normalized using SCTransform (N variable features = 2000) (12) and were decomposed using PCA (including the top 50 most variable principal components), following which they were integrated across samples using CCAIntegration. Integrated nuclei were clustered using the Leiden algorithm with a resolution of 0.75. Using control nuclei only, genes with enriched expression within each cluster were manually inspected (defined using a minimum threshold of > 5% cluster expression and $\log_{2}FC > 1$, Adj. $P < 0.05$). These genes were used as markers to provide cell-type annotations for nuclei clusters based on their correspondence with previous single-cell marker lists (13) and experimental localization studies (14).

*HSP70* expression ($\log_{10}k$) values were correlated with other genes within nuclei for each condition and across each cell-type, as well as across the whole root. Each comparison was made using Pearson correlation in nuclei with non-zero expression for both paired genes. A sparse subset of 15 global heat-shock genes were selected, which included *HSP70* and genes which displayed high expression correlation (Pearson correlation coefficient, $r > 0.5$) with *HSP70* at 35C across all nuclei cell-types.

To quantify *HSP70*-associated activity across the root, UCell (15) was used to calculate a rank-based score for each nuclei across individual cell-types. We then defined each nuclei as ON or OFF for heatshock activity using threshold based on this score. For control nuclei, this included the top 5% and remaining 95% for ON and OFF nuclei, as most nuclei displayed low scores. In the nuclei exposed to 35C however, most displayed high *HSP70*-associated UCell scores ($> 0.98$), thus the bottom 5% nuclei were defined as OFF and the remaining 95% were defined as on.

Differential gene expression between ON and OFF groups were performed for each cell-type using the pseudobulked nuclei expression aggregated within each biological replicate. EdgeR (16) was used to normalize pseudobulked counts, while limma-voom (17, 18) was used to compare gene expression between ON and OFF groups using a variance-adjusted linear model, with biological replicates incorporated as covariates. Significant genes were defined using a standard significance threshold (Benjamini-Hochberg adjusted P-value < 0.05) and a logFC > 1.5 for upregulated genes and logFC < -1.0 for downregulated genes. Differential gene lists were annotated with *A. thaliana* biological process terms using topGO (19), which were condensed using rrvgo (20). GO terms were considered significant with elimFisher P-value < 0.05.

##### 4. Stochastic Modelling of the heat stress network

The network is dynamically updated with the Gillespie Algorithm (21). We compute the rates of relevant cellular events, such as protein expression, dilution, misfolding and refolding, complex formation and dissociation, and complex transfer reactions. Given a vector of non-negative integers corresponding to the amounts of proteins and protein complexes:

$$[A1, A2, B, P70, O, M, C\{A1 \bullet P70\}, C\{P70 \bullet M\}],$$

With:

• A1: HSFA1
• A2: HSFA2
• P70: HSP70/90
• B: HSFB
• O: proteome of the cell
• M: misfolded proteins,
• C{•}: protein complex between indicated proteins

We compute these rates as follows:

$$\text{Expression}(A1) = \text{leakage\_A1} + a1 * A1^n / (h1^n + A1^n)$$

$$\text{Decay}(A1) = d * A1$$

$$\text{Expression}(P70) = \text{leakage\_P70} + a2 * A1^n / (h2^n + A1^n + B^n) + a3 * A2^n / (h3^n + A2^n + B^n)$$

$$\text{Decay}(P70) = d * P70$$

$$\text{Expression}(A2) = \text{leakage\_A2} + a4 * A1^n / (h4^n + A1^n + B^n)$$

$$\text{Decay}(A2) = d * A2$$

$$\text{Expression}(B) = \text{leakage\_B} + a5 * A1^n / (h5^n + A1^n + B^n)$$

$$\text{Decay}(B) = d * B$$

$\text{Expression}(O) = a7$
$\text{Decay}(O) = d * O$
$\text{Misfolding}(O) = d4 * O$

$\text{Decay}(M) = d5 * M$

$\text{Complex formation}\{A1 \bullet P70\} = c1 * A1 * P70$
$\text{Complex dissociation}\{A1 \bullet P70\} = d1 * C\{A1 \bullet P70\}$
$\text{Decay}(C\{A1 \bullet P70\}) = d * C\{A1 \bullet P70\}$

$\text{Complex formation}\{P70 \bullet M\} = c3 * P70 * M$
$\text{Complex dissociation}\{P70 \bullet M\} = d3 * C\{P70 \bullet M\}$
$\text{Decay}(C\{P70 \bullet M\}) = d * C\{P70 \bullet M\}$
$\text{Refolding} = a6 * C\{P70 \bullet M\}$

$\text{Complex displacement}(C\{A1 \bullet P70\} \rightarrow C\{P70 \bullet M\}) = c2 * C\{A1 \bullet P70\} * M$
$\text{Complex displacement}(C\{P70 \bullet M\} \rightarrow C\{A1 \bullet P70\}) = c4 * C\{P70 \bullet M\} * A1$

The model simulates one cell of the plant for a total of 1000 AUT. We let the system equilibrate for 600 AUT, after which we apply Heat Stress (HS) for 30 AUT, and finally we let the system relax back to its non-stressed state for the remaining time. During HS, we increase the misfolding rate 5 fold (from 0.01 to 0.05). Parameter values were chosen by iterative manual search using the experimental results as qualitative guidance (Supplementary Table S3). We ran 250 repetitions with these parameters to estimate the distribution of simulated single-cell behaviours. This allowed us to obtain statistics that can be compared to experimental data. To generate the statistics in Supplementary Figures S16-S19, we selected two pre-heat-stress time points (400, 600) and four post-heat-stress time points (630, 650, 700, 800). For Figure 4 we sampled on timepoint 630. For each target time, we sampled each simulation replicate at the recorded time point closest to the target and extracted the corresponding HSP70 value. Across-replicate variability was quantified as the coefficient of variation (SD/mean), and per-time distributions were visualized by overlaying replicate points on boxplots (median and interquartile range).

A

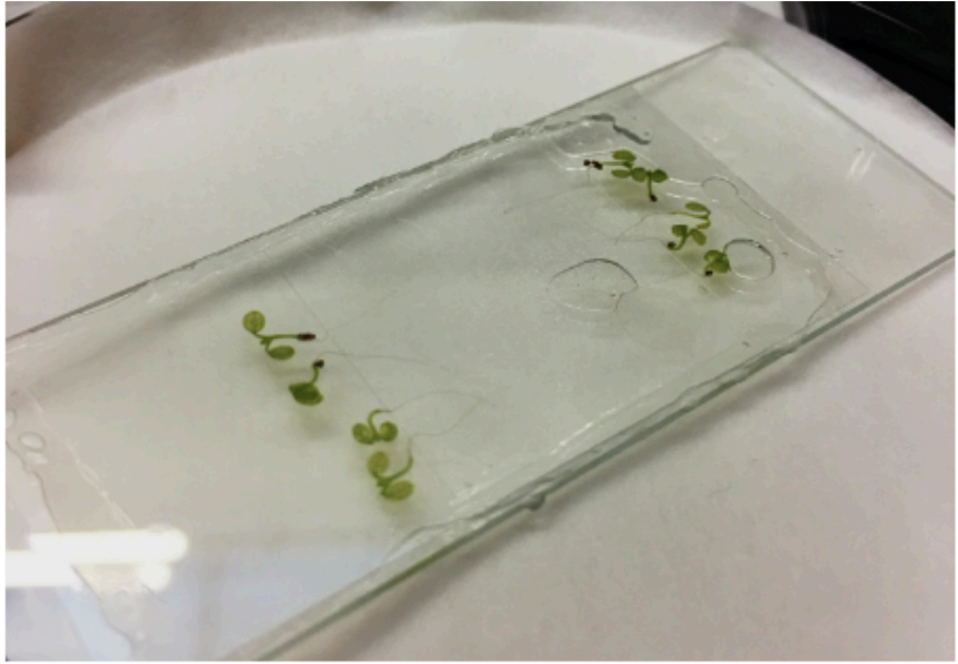

B

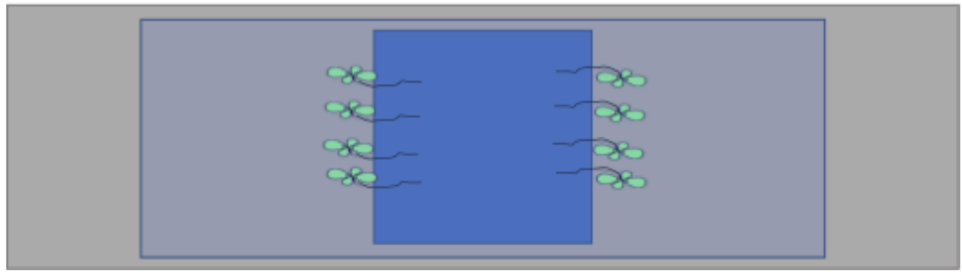

**Fig. S1. Experimental imaging setup.**

**(A)** Photograph of the sealed imaging chamber used for long-term live imaging of Arabidopsis seedlings. Seedlings were mounted on liquid half MS growth medium in a chamber that maintained humidity and nutrient access during imaging.

**(B)** Schematic of the imaging setup. Seedlings were positioned around a central imaging region to allow repeated imaging of roots while minimising disturbance during heat stress and recovery experiments.

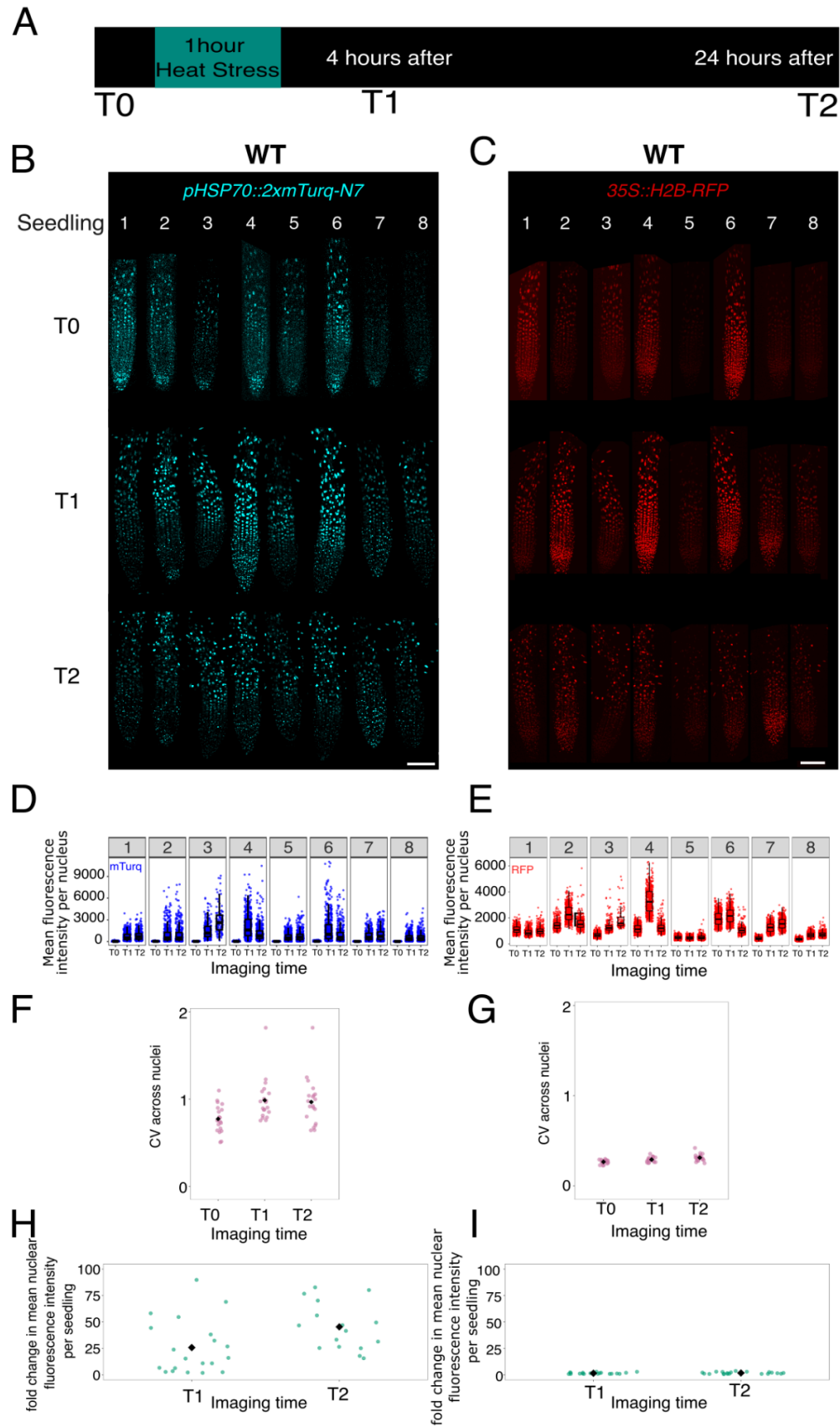

**Fig. S2. Independent 1 h heat stress repeat acquired on a second microscope confirms heterogeneous *HSP70* induction.**

**(A)** Imaging scheme. WT seedlings were imaged before heat stress (T0), then re-imaged 4 h (T1) and 24 h (T2) after a 1 h heat stress treatment.

**(B, C)** Representative confocal images from an independent repeat experiment acquired on a Zeiss 880 microscope, rather than the Nikon AXR microscope used for the other imaging datasets, showing *pHSP70::2xmTurq-N7* reporter signal (B) and the constitutive nuclear marker *35S::H2B-RFP* (C) in the same roots at T0, T1 and T2. Images are shown with adjusted display ranges for visualisation: mTurquoise2, T0 root 1, 250–300; T0 root 2, 200–300; remaining T0 roots, 150–300; T1 and T2, 150–6000; RFP, 200–5000.

**(D, E)** Quantification of nuclear fluorescence intensity for the *HSP70* reporter (D) and *35S::H2B-RFP* control (E). Each numbered panel represents one seedling; points represent individual nuclei.

**(F, G)** Coefficient of variation (CV) of nuclear fluorescence intensity across nuclei within individual seedlings for the *HSP70* reporter (F) and *35S::H2B-RFP* control (G) at each imaging time point.

**(H, I)** Fold change in mean nuclear fluorescence intensity per seedling at T1 and T2 relative to T0 for the *HSP70* reporter (H) and *35S::H2B-RFP* control (I). For (F-I), n = 20 seedlings from 3 independent 1 h heat stress experiments.

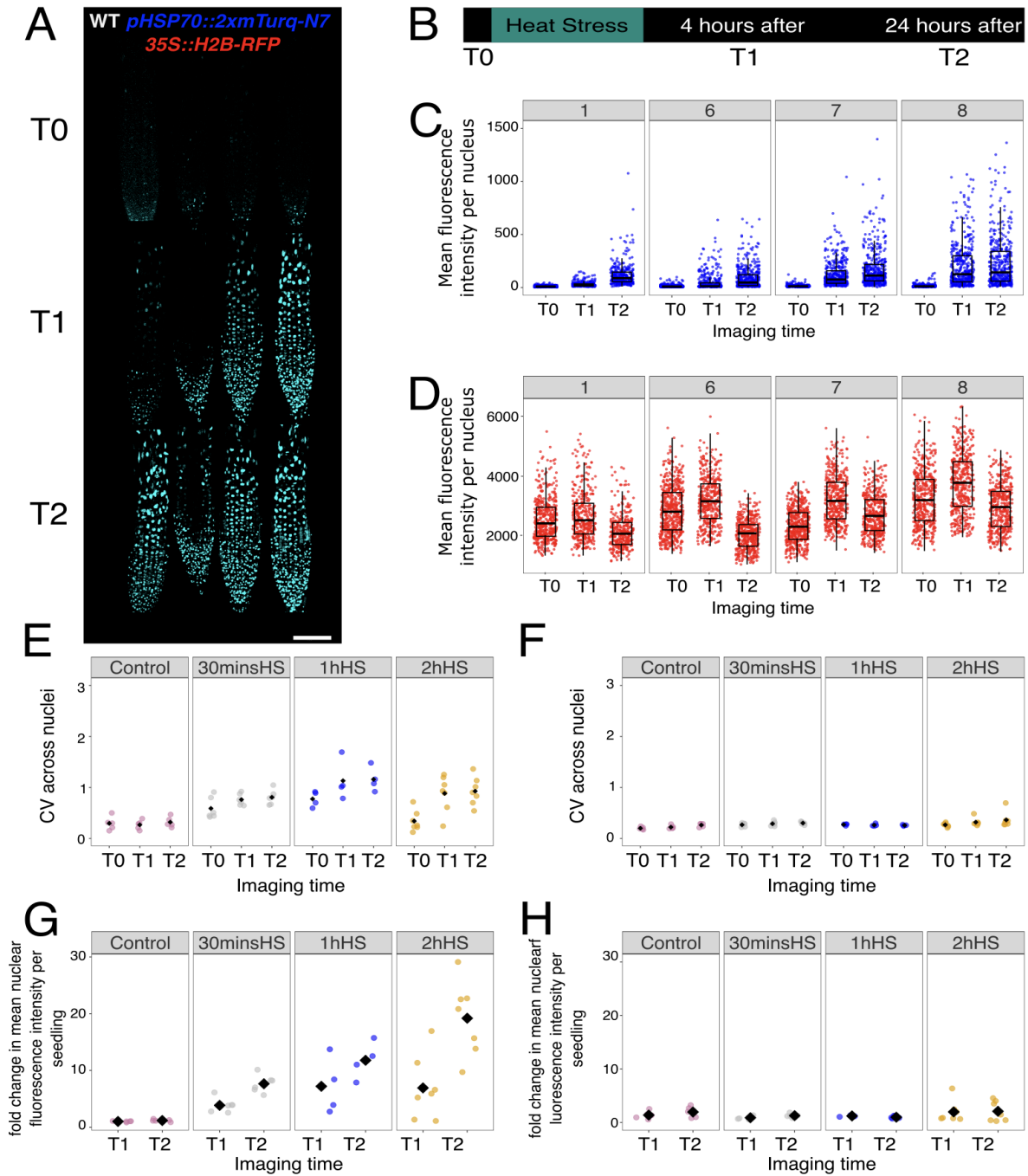

**Fig. S3. A second independent *HSP70* reporter insertion line shows heterogeneous induction after heat stress.**

**(A)** Representative confocal images of roots from an independent *pHSP70::2xmTurq-N7* reporter line carrying the constitutive nuclear marker *35S::H2B-RFP*, imaged before heat stress (T0), 4 h after (T1), and 24 h after 1 h heat stress (T2). Images are shown with adjusted display ranges for visualisation: mTurquoise2, root 1 at T0, 200–250; all other images, 180–300.

**(B)** Experimental timeline. Seedlings were imaged before heat stress, exposed to 35 °C for 30 min, 1 h, or 2 h, and re-imaged 4 h and 24 h later.

**(C, D)** Quantification of nuclear fluorescence intensity for the *pHSP70::2xmTurq-N7* reporter **(C)** and *35S::H2B-RFP* control **(D)** in the four seedlings shown in A, at T0, T1, and T2. Each point represents one nucleus.

**(E, F)** Cell-to-cell variability, measured as the coefficient of variation across nuclei within each seedling, for the *pHSP70::2xmTurq-N7* reporter **(E)** and *35S::H2B-RFP* control **(F)** under control conditions and after 30 min, 1 h, or 2 h heat stress. This independent reporter line recapitulates the heterogeneous increase in *HSP70* promoter activity after heat stress.

**(G, H)** Fold change in mean nuclear fluorescence intensity per seedling relative to the pre-stress time point for the *pHSP70::2xmTurq-N7* reporter **(G)** and *35S::H2B-RFP* control **(H)** under control conditions and after 30 min, 1 h, or 2 h heat stress .

For (E–H), each point represents one seedling and black diamonds indicate the mean. Sample sizes were: no-heat control, n = 5 seedlings; 30 min heat stress, n = 6 seedlings; 1 h heat stress , n = 5 seedlings; 2 h heat stress , n = 8 seedlings, each from one imaging experiment per stress duration.

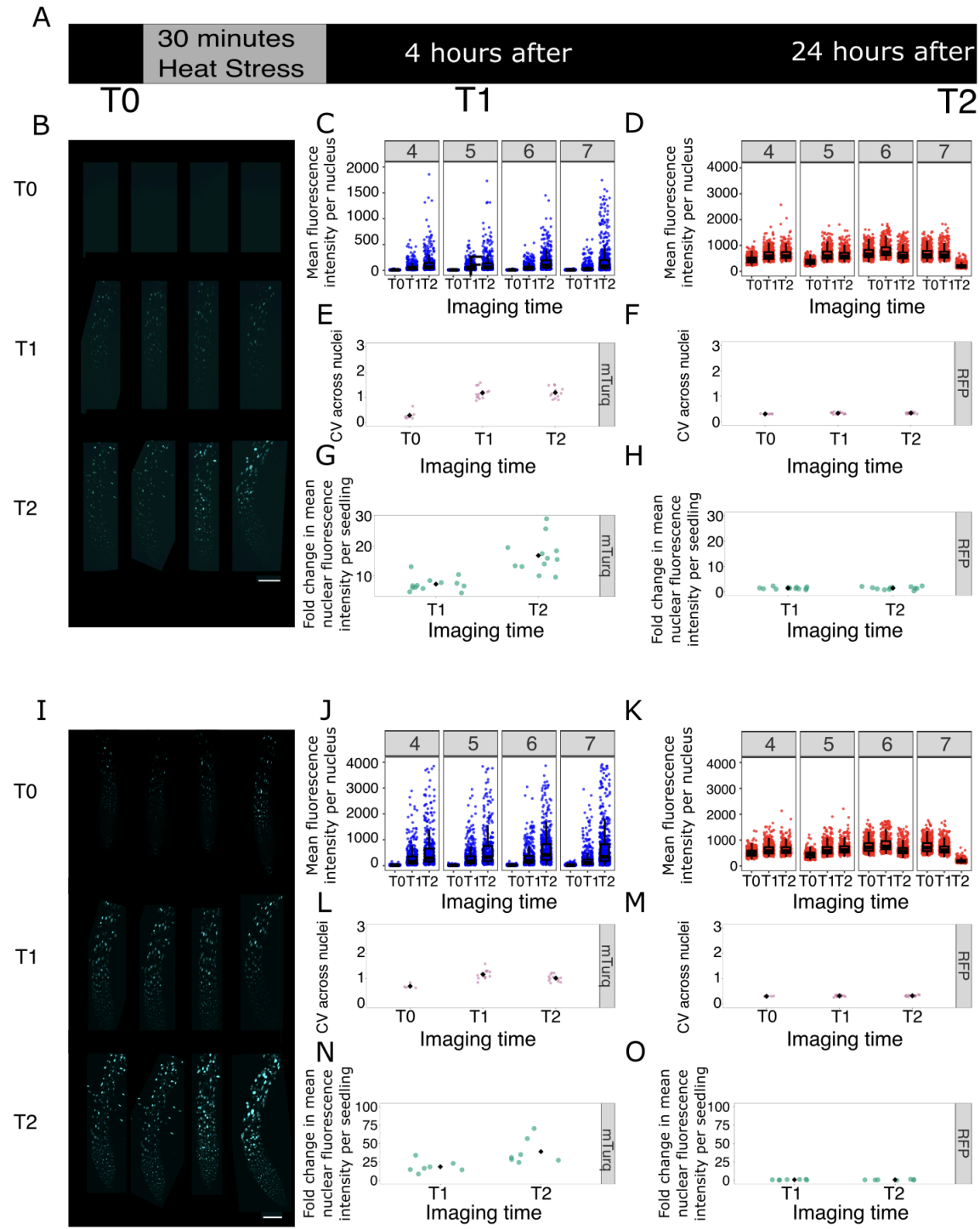

**Fig. S4. Detailed 30 min heat stress analysis and CFP acquisition setting comparison.**

**(A)** Imaging scheme. Seedlings were imaged before heat stress (T0), exposed to 35 °C for 30 min, and re-imaged 4 h (T1) and 24 h (T2) after heat stress.

**(B–H)** WT 30 min heat shock dataset acquired using low CFP acquisition settings. This is the 30 min dataset summarised in Fig. 1 and replotted as the WT comparator for the *hsfa2* 30 min analysis in Fig. 5 and Fig. S22. Data in (E–H) are plotted from $n = 12$ seedlings from 2 independent experiments.

**(B)** Representative confocal images of roots expressing *pHSP70::2xmTurq-N7* before and after heat stress.

**(C, D)** Quantification of nuclear fluorescence intensity for *pHSP70::2xmTurq-N7* **(C)** and *35S::H2B-RFP* **(D)** in individual seedlings. Each point represents one nucleus.
**(E, F)** Cell-to-cell variability, measured as the coefficient of variation (CV) across nuclei within each seedling, for *pHSP70::2xmTurq-N7* **(E)** and *35S::H2B-RFP* **(F)**.
**(G, H)** Fold change in mean nuclear fluorescence intensity per seedling at T1 and T2 relative to T0 for *pHSP70::2xmTurq-N7* **(G)** and *35S::H2B-RFP* **(H)**.

**(I–O)** Acquisition-setting comparison for the 30 min heat stress experiment. The same seedlings used for the low CFP measurements in B–H were additionally imaged using the standard CFP acquisition settings used for the 1 h and 2 h heat stress experiments. In one experiment, standard CFP images were acquired after heat stress only, at T1 and T2. In the second experiment, standard CFP images were acquired at all three time points, T0, T1 and T2.

**(I)** Representative confocal images of roots expressing *pHSP70::2xmTurq-N7* before and after heat stress.

**(J, K)** Quantification of nuclear fluorescence intensity for *pHSP70::2xmTurq-N7* **(J)** and *35S::H2B-RFP* **(K)**.

**(L, M)** Cell-to-cell variability, measured as the coefficient of variation (CV) across nuclei within each seedling, for *pHSP70::2xmTurq-N7* **(L)** and *35S::H2B-RFP* **(M)**.

**(N, O)** Fold change in mean nuclear fluorescence intensity per seedling at T1 and T2 relative to T0 for *pHSP70::2xmTurq-N7* **(N)** and *35S::H2B-RFP* **(O)**.

The standard CFP settings caused saturation in induced *pHSP70::2xmTurq-N7* images after 30 min heat stress, so the low-CFP dataset was used for Fig. 1 and for the WT comparison in the *hsfa2* 30 min analysis. Because CFP acquisition settings differ between datasets, absolute *pHSP70::2xmTurq-N7* fluorescence values should not be directly compared between **(B–H)** and **(I–O)**; fold changes were calculated within seedlings relative to their matched pre-stress values. Both acquisition settings showed heterogeneous *HSP70* induction after 30 min heat stress.

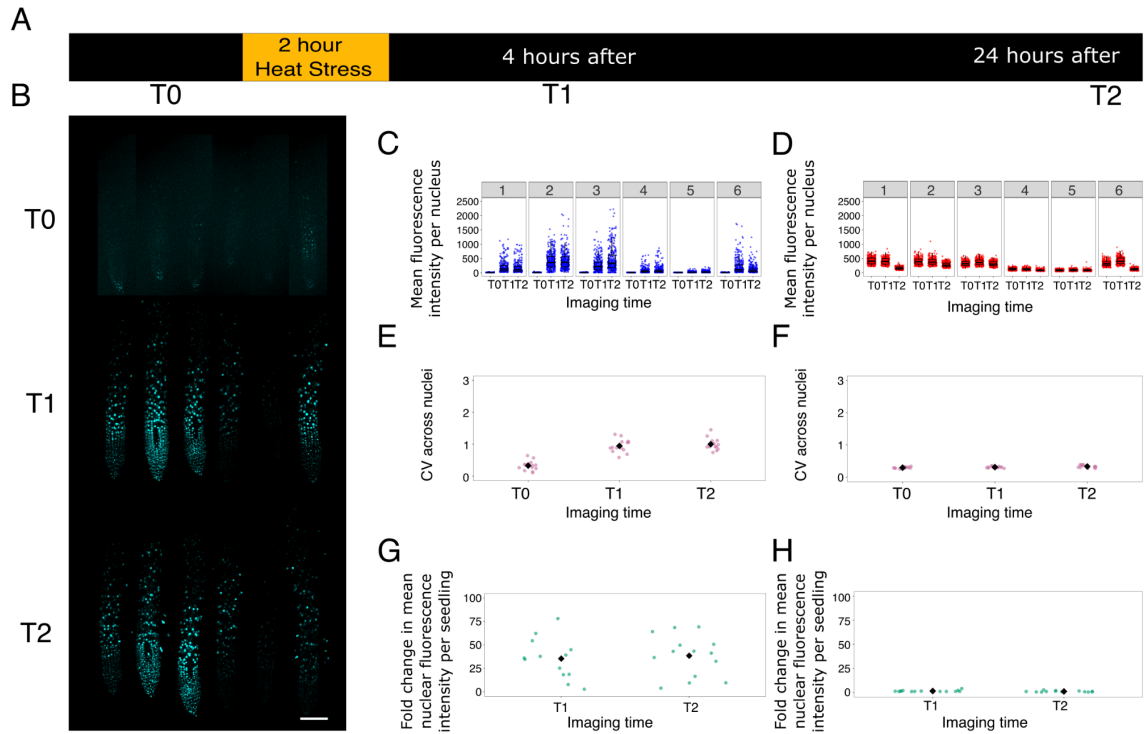

**Fig. S5. 2 hour heat stress experiment increases *HSP70* expression without corresponding changes in the constitutive nuclear marker.**

**(A)** Imaging scheme. WT seedlings were imaged before heat stress (T0), then re-imaged 4 h (T1) and 24 h (T2) after a 2 h heat stress treatment.

**(B)** Representative confocal images from an independent repeat experiment showing *pHSP70::2xmTurq-N7* reporter signal in the same roots at T0, T1 and T2. (Display range - CFP Before: 200-250, after 4 hours and 24 hours: 100-1500)

**(C, D)** Quantification of nuclear fluorescence intensity for *pHSP70::2xmTurq-N7* **(C)** and *35S::H2B-RFP* **(D)** in individual seedlings. Each point represents one nucleus. Boxplots summarise the distribution of pixel intensities, with points showing individual nuclei.

**(E, F)** Coefficient of variation (CV) of nuclear fluorescence intensity across nuclei within each seedling, for *pHSP70::2xmTurq-N7* **(E)** and *35S::H2B-RFP* **(F)** at each imaging timepoint.

**(G, H)** Fold change in mean nuclear fluorescence intensity per seedling at T1 and T2 relative to T0 for *pHSP70::2xmTurq-N7* **(G)** and *35S::H2B-RFP* **(H)**.

The WT 2 h dataset shown here is summarised in Fig. 1 and replotted as the WT comparator for the *hsfa2* 2 h analysis in Fig. 5 and Fig. S23. Data shown in (E-H) are plotted from n = 15 seedlings from 2 independent experiments.

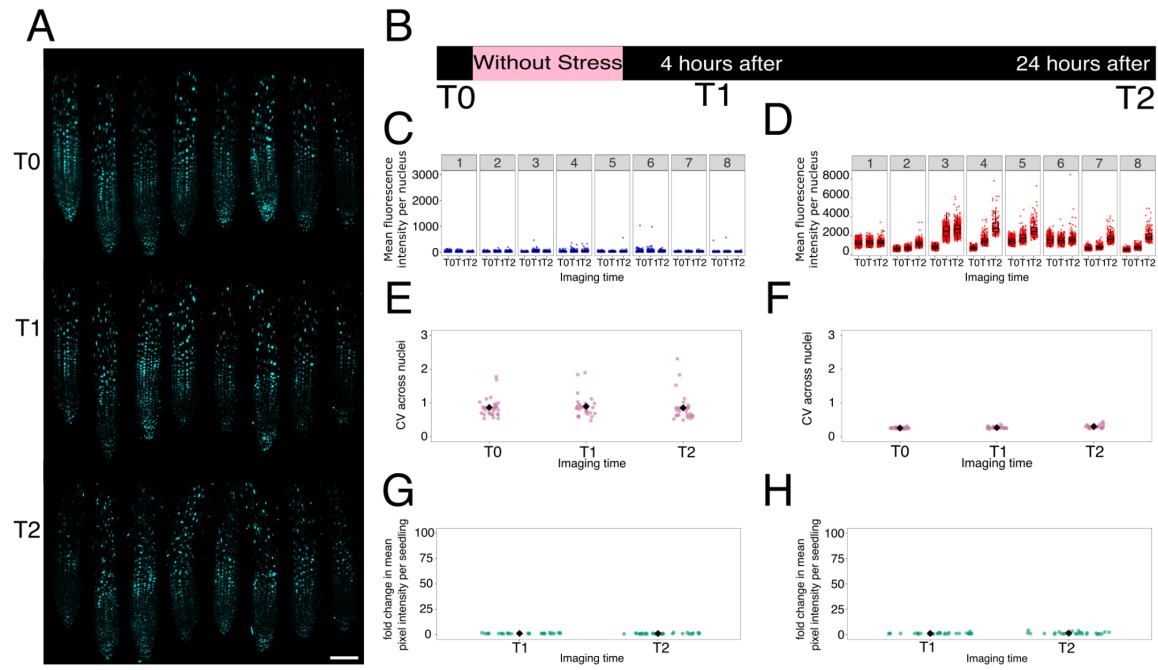

**Fig. S6. Control time-course experiments without heat stress show stable *HSP70* and constitutive *H2B* signals.**

**(A)** Representative confocal images from a control time-course experiment performed without heat stress, showing *pHSP70::2xmTurq-N7* signal in WT roots at T0, T1 and T2 (Display range : 120-200).

**(B)** Imaging scheme. Seedlings were imaged at the same time intervals used in the heat stress experiments, but without heat treatment.

**(C, D)** Quantification of pixel intensity in individual seedlings from a representative control experiment for *pHSP70::2xmTurq-N7* **(C)** and *35S::H2B-RFP* **(D)**. Each numbered panel represents one seedling. Boxplots summarise the distribution of pixel intensities, with points showing individual nuclei.

**(E, F)** Coefficient of variation (CV) of nuclear fluorescence intensity across nuclei within individual seedlings for *pHSP70::2xmTurq-N7* **(E)** and *35S::H2B-RFP* **(F)**, summarised across the same control experiments. In the absence of heat stress, both *pHSP70::2xmTurq-N7* and *35S::H2B-RFP* signals remained broadly stable in mean intensity and heterogeneity over time.

**(G, H)** Fold change in mean pixel intensity per seedling at T1 and T2 relative to T0 for *pHSP70::2xmTurq-N7* **(G)** and *35S::H2B-RFP* **(H)**, summarised across independent control experiments.

Fold-change and CV summaries in (E–H) are from $n = 30$ plants from 4 independent control experiments. Images were acquired on a Zeiss 880 microscope.

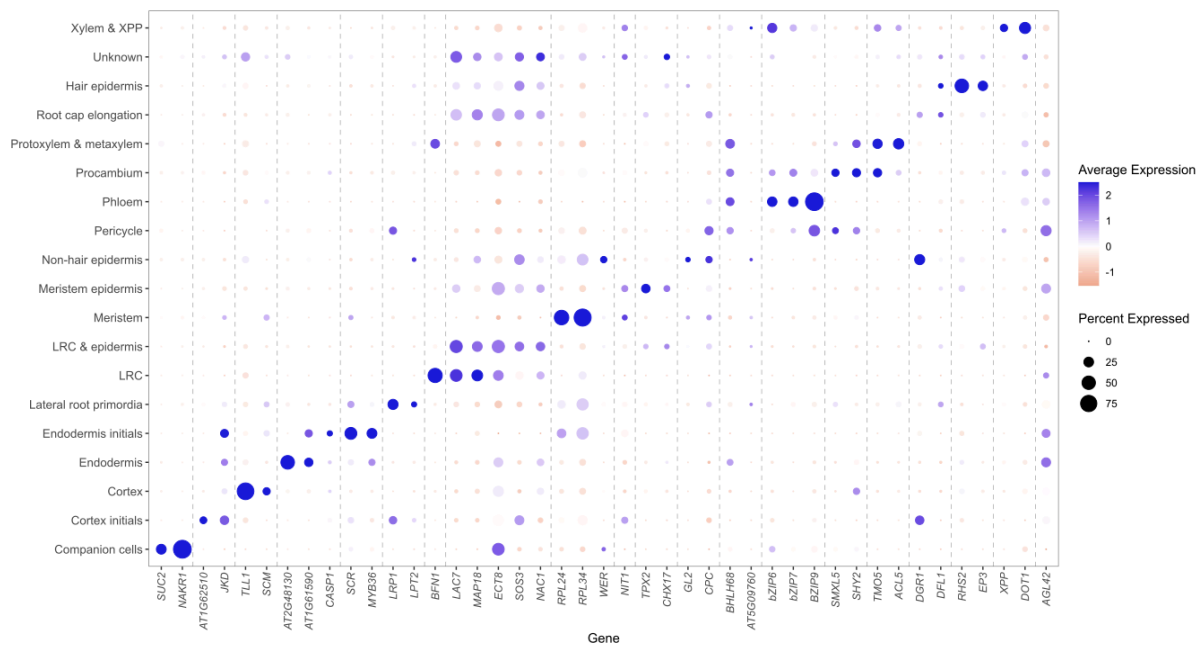

**Fig. S7. Marker gene expression supports root cell-type annotations in the integrated snRNA-seq dataset.**

Dot plot showing expression of selected marker genes across annotated root cell types in the integrated snRNA-seq dataset. Dot size indicates the percentage of nuclei in each annotated cell type expressing the marker gene, and colour indicates scaled average expression. Marker gene expression supports the assignment of major root cell populations shown in Fig. 3A.

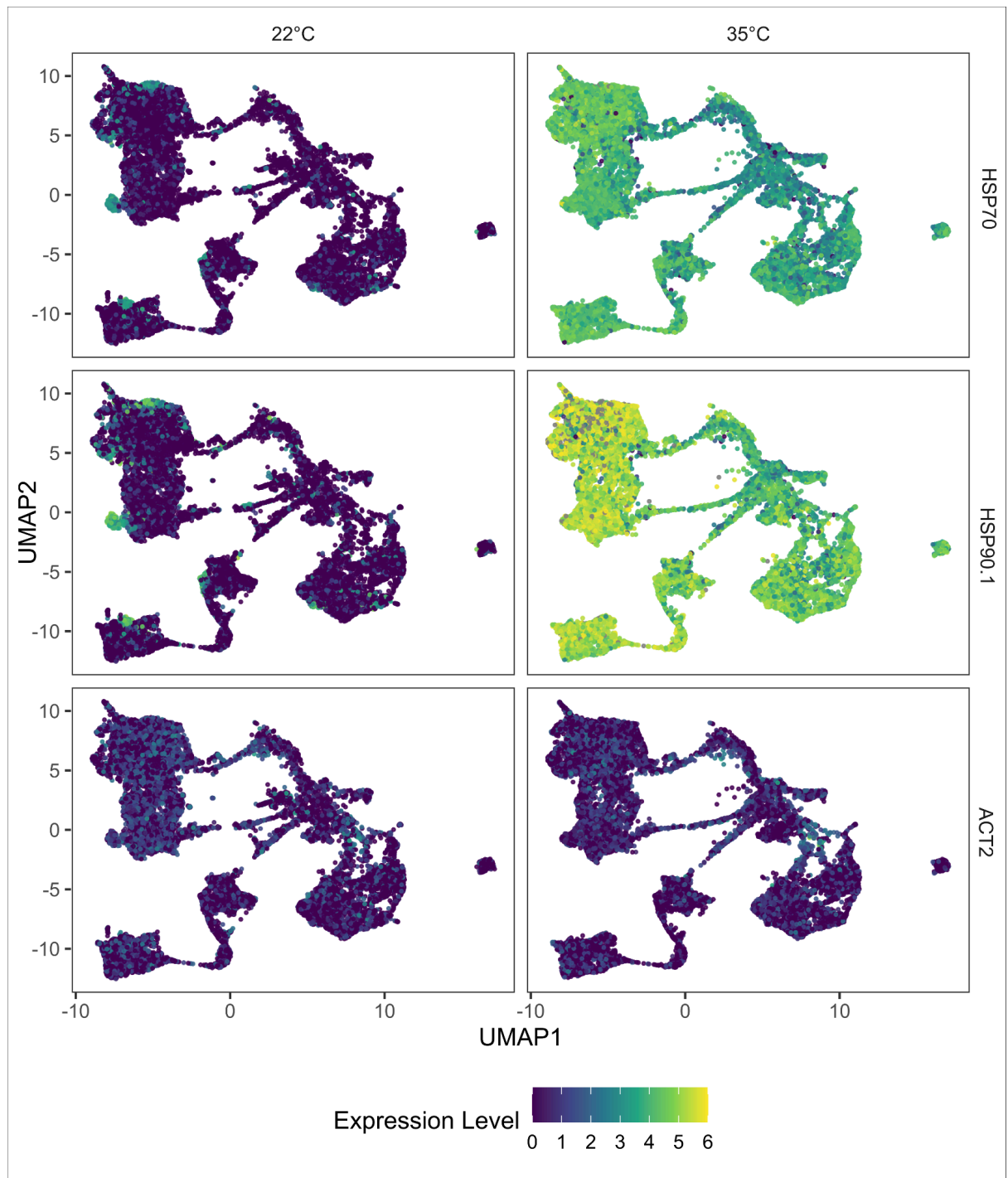

**Fig. S8. UMAPs showing expression of *HSP70*, *HSP90* and *ACT2* in single nuclei from control and heat stress root samples.** Expression of *HSP70* /AT3G12580, *HSP90*/AT5G52640, and *ACT2*/AT3G18780 is shown for control seedlings at 22 °C and seedlings exposed to 35 °C heat stress. Each point represents a nucleus; colour intensity indicates normalised expression level.

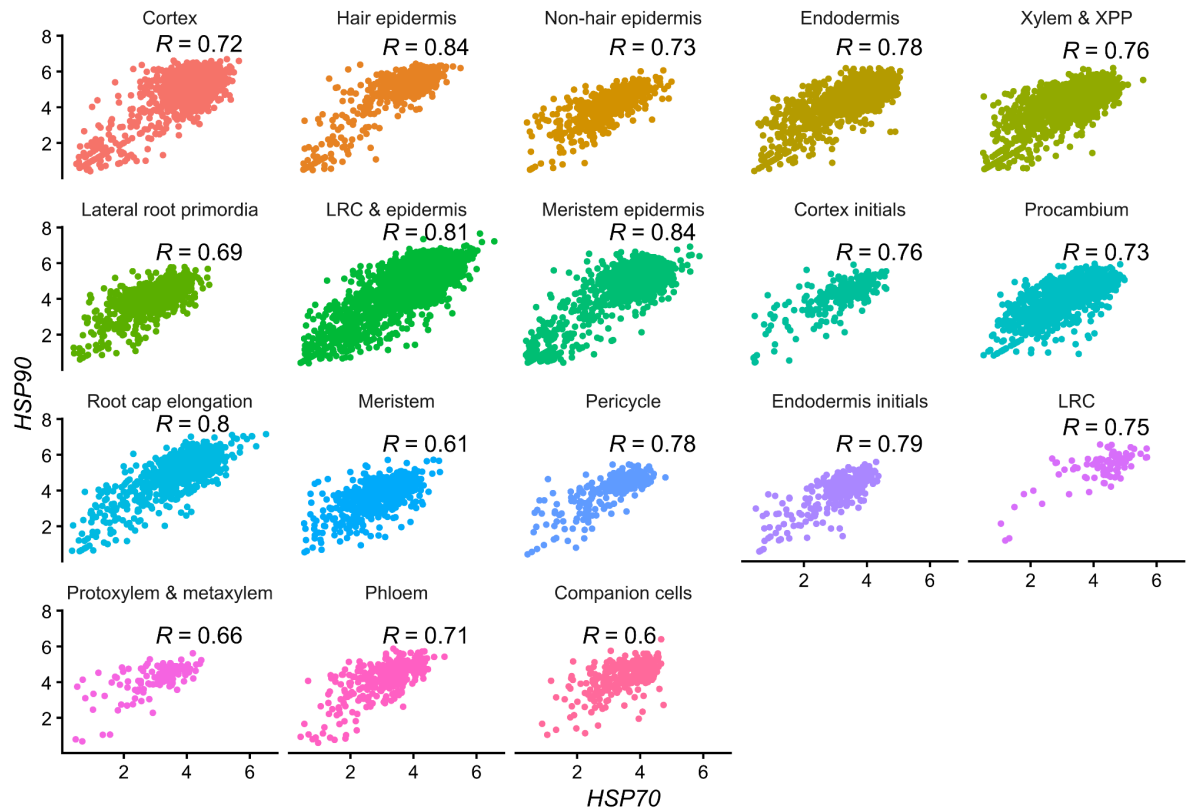

**Fig. S9. *HSP70* and *HSP90* expression covary across root cell types after heat stress.** Scatter plots show expression of *HSP70* and *HSP90* in individual nuclei from the 35 °C fresh heat stress samples, non-zero Seurat-normalized (LogNormalize, scale factor 10K) counts, separated by annotated cell type. Each point represents a nucleus. Pearson correlation coefficients (*R*) are indicated for each cell type.

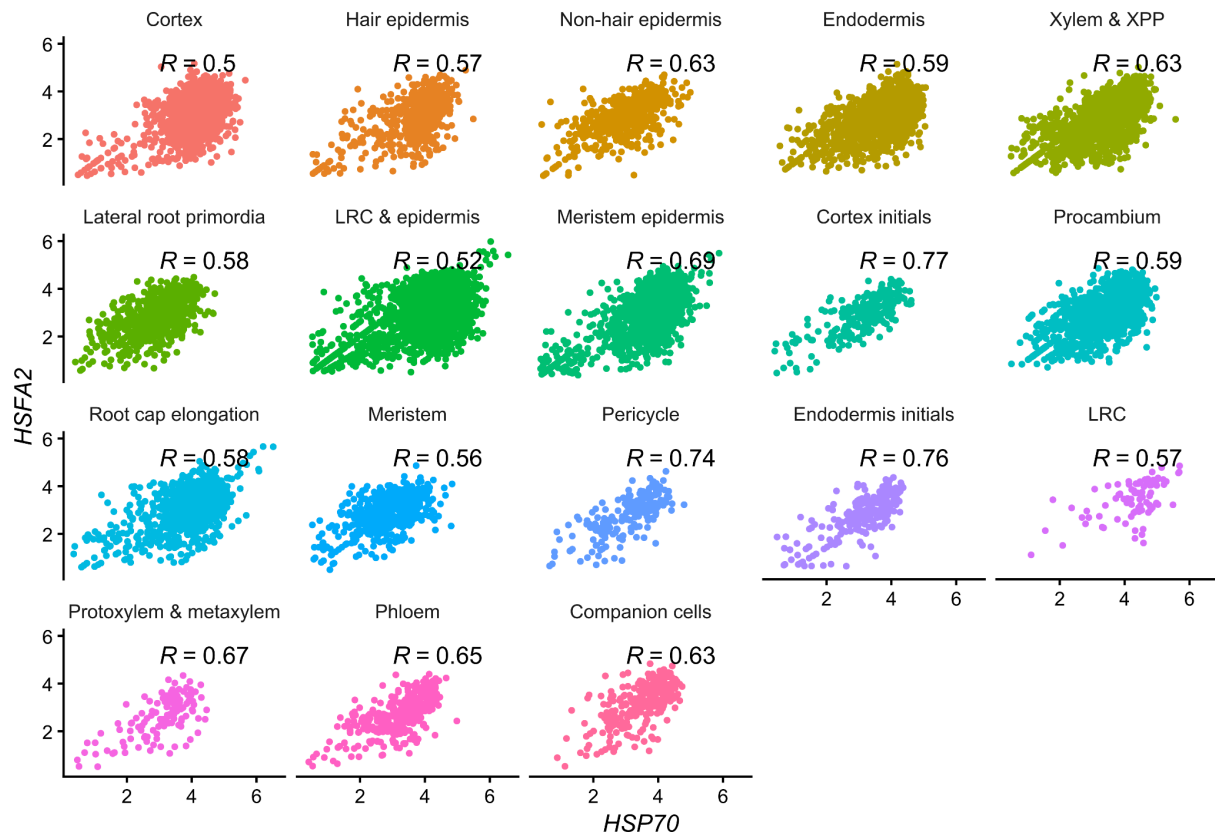

**Fig. S10. *HSP70* and *HSFA2* expression covary across root cell types after heat stress.** Scatter plots show expression of *HSP70* and *HSFA2* in individual nuclei from the 35 °C fresh heat stress samples, separated by annotated cell type, non-zero Seurat-normalized (LogNormalize, scale factor 10K) counts. Each point represents a nucleus. Pearson correlation coefficients (R) are indicated for each cell type.

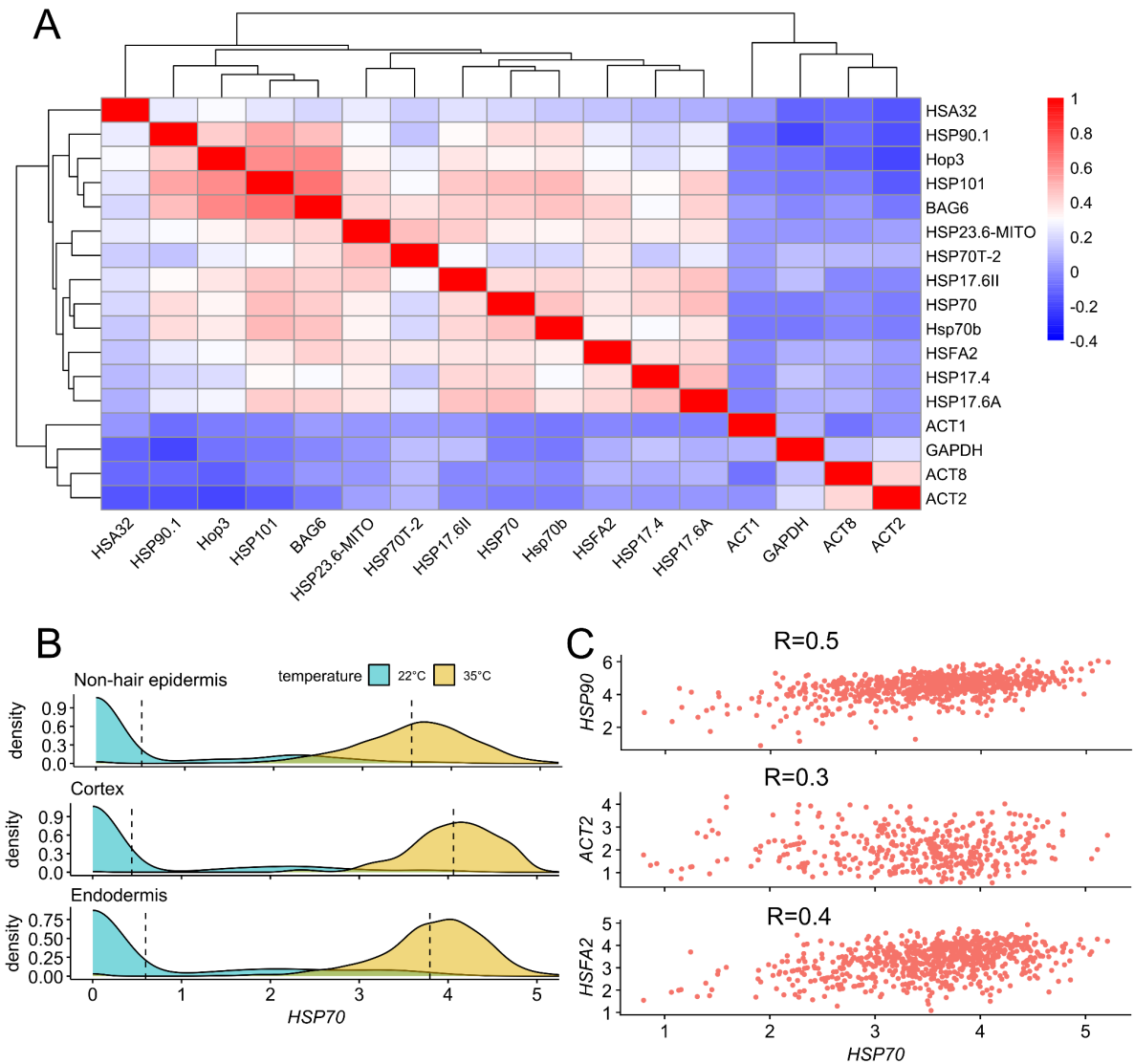

**Fig. S11. Liquid nitrogen frozen snRNA-seq replicate supports heterogeneous and coordinated heat shock gene activation.**

**(A)** Pairwise gene-gene correlation heatmap for heat shock responsive genes and control genes across nuclei from the 35 °C frozen sample. Heat shock genes show coordinated positive covariation, whereas control genes show weaker correlations.

**(B)** Distribution of *HSP70* expression in selected cell types in control (22 °C) and heat stress (35 °C) frozen samples.

**(C)** Single-nucleus expression correlations in the 35 °C frozen sample between *HSP70* and selected genes. Single-nucleus expression correlations in the 35 °C frozen sample between *HSP70* and selected genes. *HSP70* expression was positively correlated with *HSP90* ($R = 0.5$) and *HSFA2* ($R = 0.4$), with a somewhat lower correlation with *ACT2* ($R = 0.3$). Pearson correlation coefficients ($R$) are indicated.

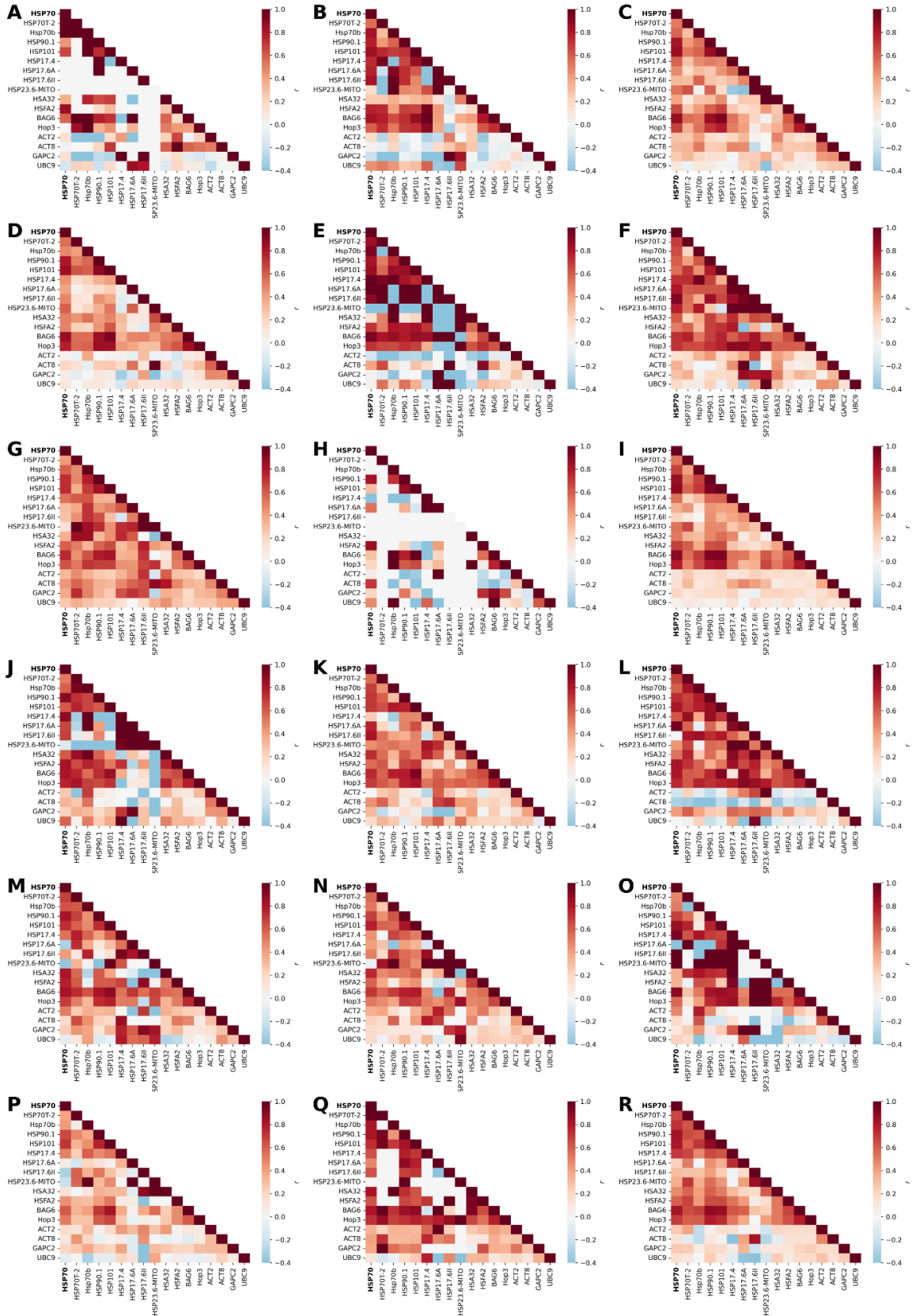

Fig. S12. Pairwise gene-gene correlation heatmap for heat shock responsive

**genes and control genes across nuclei from different cell types in 22 °C fresh samples.** A) Companion cells, B) Cortex initials, C) Cortex, D) Endodermis, E) Endodermis initials, F) Lateral root primordia, G) LRC & epidermis, H) LRC, I) Meristem epidermis, J) Meristem, K) Non-hair epidermis, L) Pericycle, M) Phloem, N) Procambium, O) Protoxylem & metaxylem, P) Root cap elongation, Q) Hair epidermis, R) Xylem & XPP

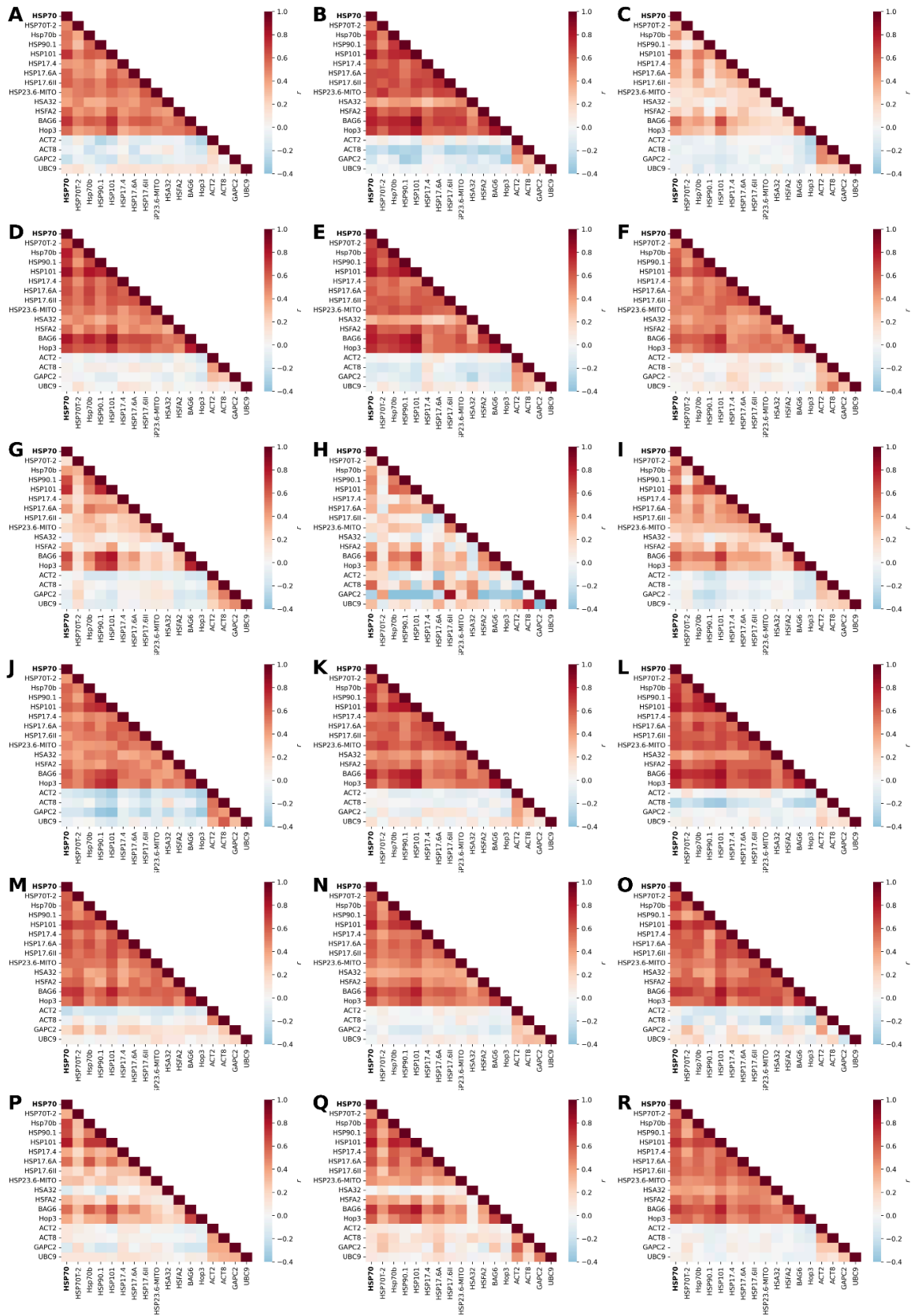

**Fig. S13. Pairwise gene-gene correlation heatmap for heat shock responsive genes and control genes across nuclei from different cell types in 35 °C fresh samples. A) Companion cells, B) Cortex initials, C) Cortex, D) Endodermis, E) Endodermis initials, F) Lateral root primordia, G) LRC & epidermis, H) LRC, I)**

Meristem epidermis, J) Meristem, K) Non-hair epidermis, L) Pericycle, M) Phloem, N) Procambium, O) Protoxylem & metaxylem, P) Root cap elongation, Q) Hair epidermis, R) Xylem & XPP

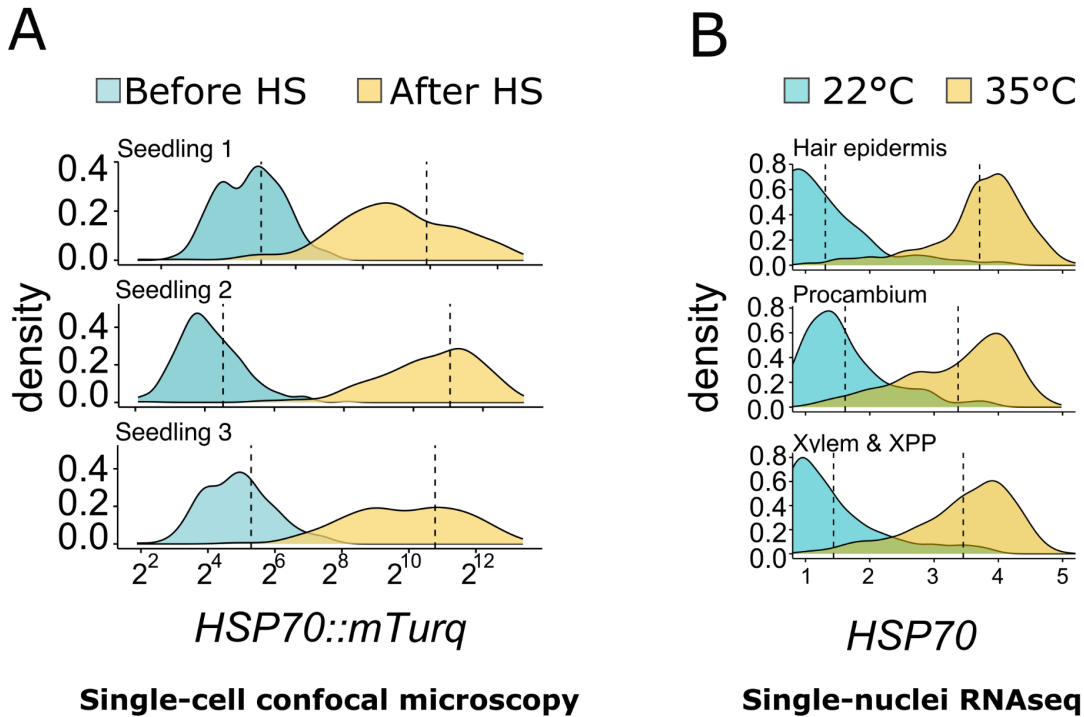

**Fig. S14. Confocal reporter imaging and snRNA-seq show qualitatively similar heterogeneous *HSP70* response distributions.**

**(A)** Distribution of nuclear *pHSP70::2xmTurq-N7* reporter fluorescence before heat stress and 4 h after a 1 h heat stress at 35 °C, measured by confocal microscopy. Each row shows one representative root.

**(B)** Distribution of endogenous *HSP70* expression in selected root cell types from control seedlings at 22 °C and seedlings exposed to 35 °C heat stress, measured by snRNA-seq.

Although the two measurements report different molecular outputs, accumulated reporter fluorescence in A and endogenous transcript abundance in B, both show a heat-induced shift towards higher *HSP70* signal while retaining broad cell-to-cell heterogeneity. Data in A are replotted from Fig. S2. Data in B are from the fresh tissue snRNA-seq datasets.

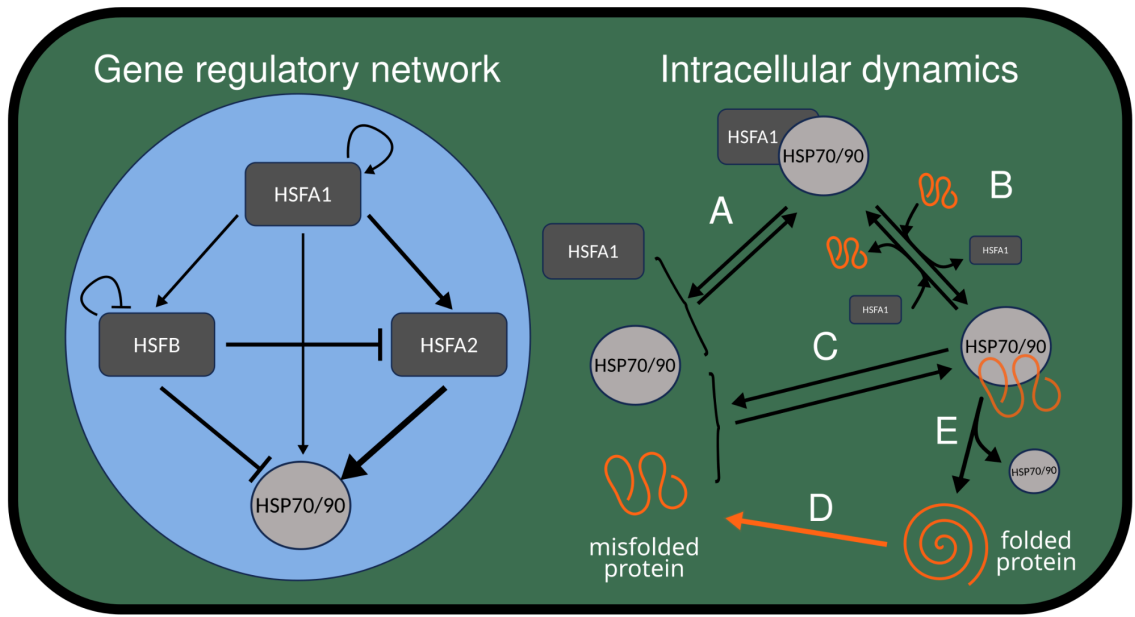

**Fig. S15. Detailed regulatory interactions and intracellular reactions represented in the stochastic model.**

Left, gene regulatory network included in the model. HSFA1 activates expression of HSFA2, HSFB and HSP70/HSP90, while HSFA2 further activates HSP70/HSP90 expression and HSFB represses heat shock gene expression. Right, intracellular chaperone dynamics represented in the model.

**(A)** In the absence of heat stress, HSFA1 forms a complex with HSP70/HSP90, limiting HSFA1 activity.

**(B, C)** During heat stress, misfolded proteins compete for HSP70/HSP90 binding, promoting release of HSFA1 and formation of HSP70/HSP90-misfolded protein complexes.

**(D)** Folded proteins misfold at a low rate under control conditions and at a higher rate during heat stress.

**(E)** HSP70/HSP90 promotes refolding of misfolded proteins and is released after refolding.

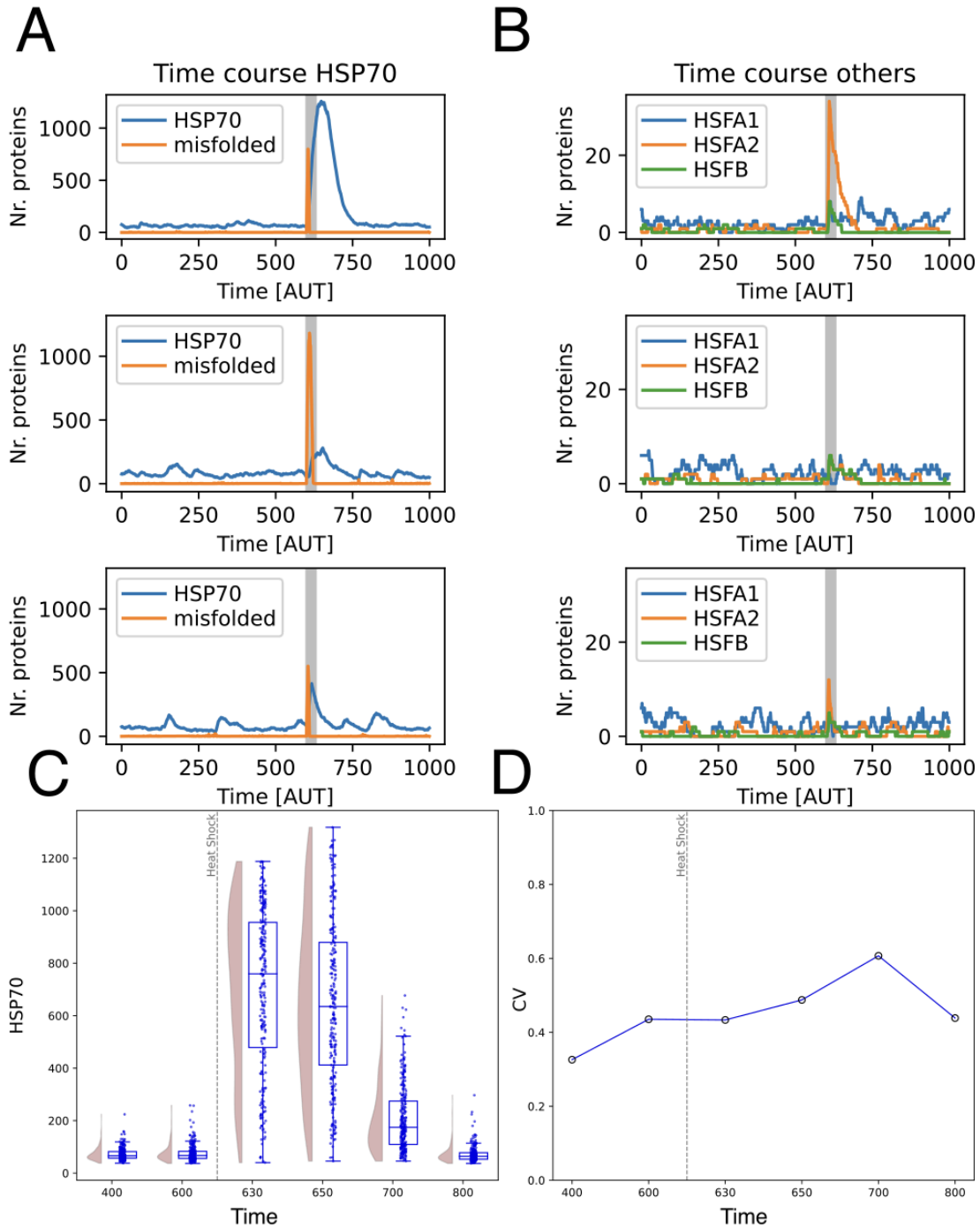

**Fig. S16. Stochastic simulations of the WT model reveal transient increases in HSP70 abundance and heterogeneity after heat stress.**

(A, B) Representative stochastic simulation trajectories over 1000 arbitrary time units (AUT). (A) HSP70 and misfolded protein levels. (B) HSFA1, HSFA2 and HSFB levels. The grey shaded region indicates the heat stress period. The three rows show example trajectories illustrating variable single-cell responses in the WT model. (C) Distribution of simulated HSP70 levels across cells at the indicated time points before and after heat shock. Violin plots show the full distribution, boxplots show the median and interquartile range, and points show individual simulated cells.

**(D)** Coefficient of variation (CV) of HSP70 levels across simulated cells at the same time points as in C. Heterogeneity increases after heat shock, peaks during the post-stress response, and then declines as HSP70 levels relax towards baseline.

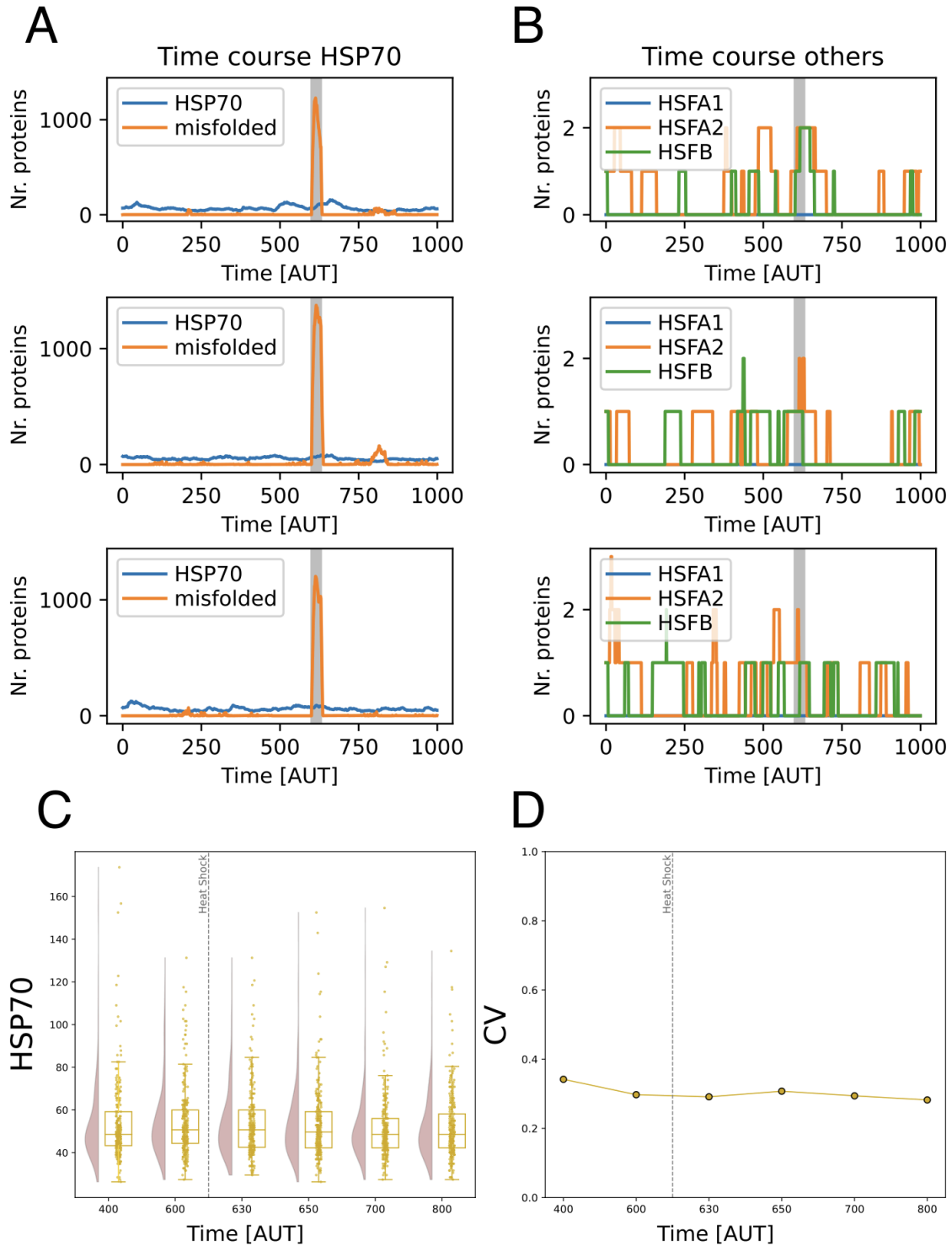

**Fig. S17. Stochastic simulations of the *hsfA1* mutant model show little change in HSP70 abundance or heterogeneity after heat stress.**

(A, B) Representative stochastic simulation trajectories over 1000 AUT for the *hsfA1* mutant model. (A) HSP70 and misfolded protein levels. (B) HSF1, HSF2 and HSF3 levels. Grey shading indicates the heat stress period.

(C) Distributions of simulated HSP70 levels across cells at selected timepoints before and after heat stress. (D) The corresponding CV values. Unlike the WT model, the *hsfA1* mutant shows little change in HSP70 levels or heterogeneity after heat stress.

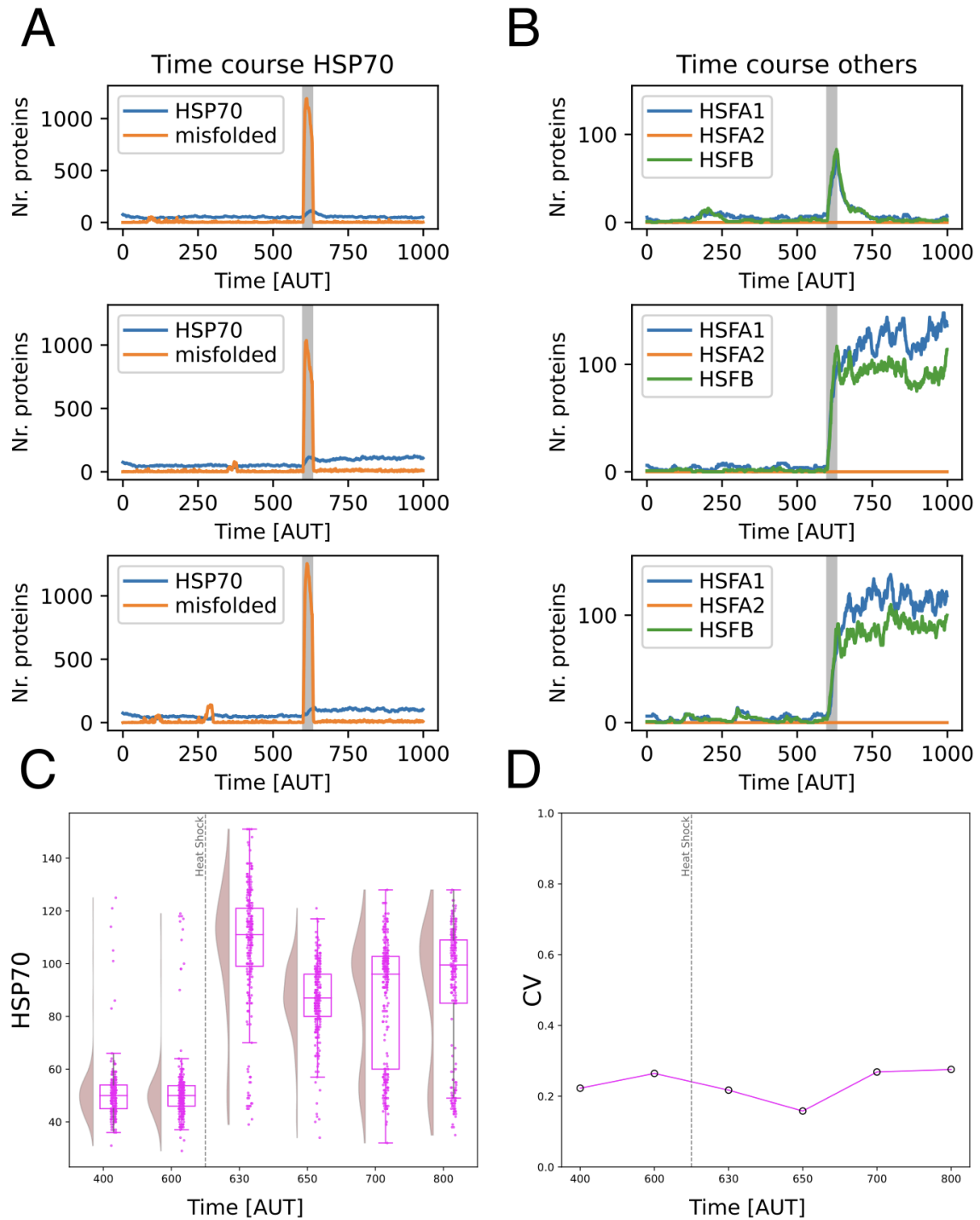

**Fig. S18. Stochastic simulations of the *hsfA2* mutant model show a modest HSP70 response with limited changes in heterogeneity after heat stress.**

(A, B) Representative stochastic simulation trajectories over 1000 arbitrary time units (AUT) for the *hsfA2* mutant model. (A) HSP70 and misfolded protein levels. (B) HSFA1, HSFA2 and HSFB levels. The grey shaded region indicates the heat stress period. The three rows show example trajectories illustrating the range of single-cell responses in the mutant model.

(C) Distribution of simulated HSP70 levels across cells at the indicated timepoints before and after heat shock (400, 600, 630, 650, 700 and 800 AUT). Violin plots show the full distribution, boxplots show the median and interquartile range, and points show individual simulated cells. The dashed vertical line marks the onset of heat stress.

**(D)** Coefficient of variation (CV) of HSP70 levels across simulated cells at the same timepoints. In the *hsfA2* mutant model, HSP70 levels increase after heat stress but remain lower and less heterogeneous than in the WT model.

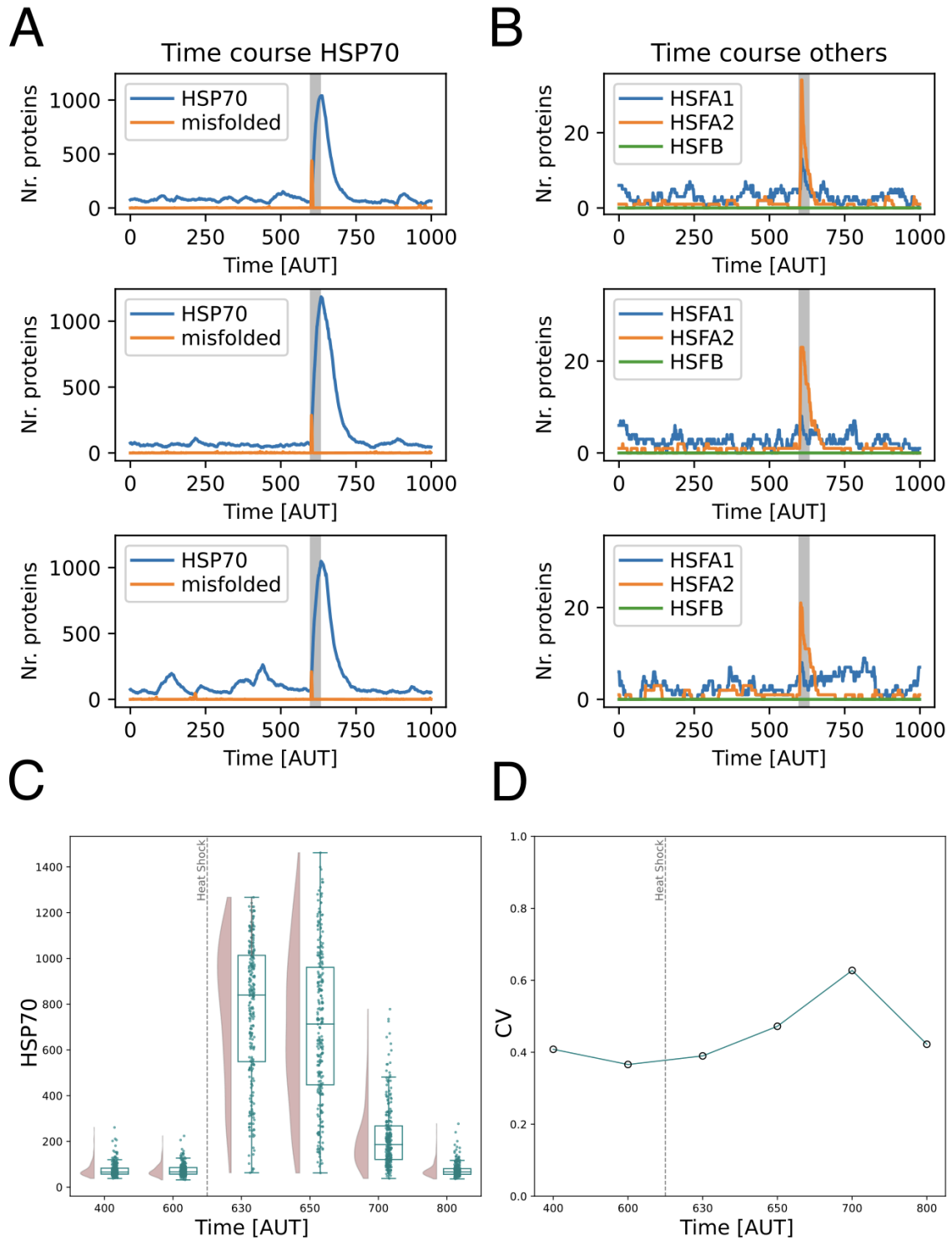

**Fig. S19. Stochastic simulations of the *hsfB* mutant model show HSP70 induction dynamics broadly similar to WT.**

**(A, B)** Representative stochastic simulation trajectories over 1000 arbitrary time units (AUT) for the *hsfB* mutant model. **(A)** HSP70 and misfolded protein levels. **(B)** HSFA1, HSFA2 and HSFB levels. The grey shaded region indicates the heat stress period. The three rows show example trajectories illustrating variable single-cell responses in the mutant model.

**(C)** Distribution of simulated HSP70 levels across cells at the indicated timepoints before and after heat shock (400, 600, 630, 650, 700 and 800 AUT). Violin plots show the full distribution, boxplots show the median and interquartile range, and points show individual simulated cells. The dashed vertical line marks the onset of heat stress.

**(D)** Coefficient of variation (CV) of HSP70 levels across simulated cells at the same timepoints. The *hsfB* mutant model shows a post-stress increase in *HSP70* levels and heterogeneity that is broadly similar to the WT model.

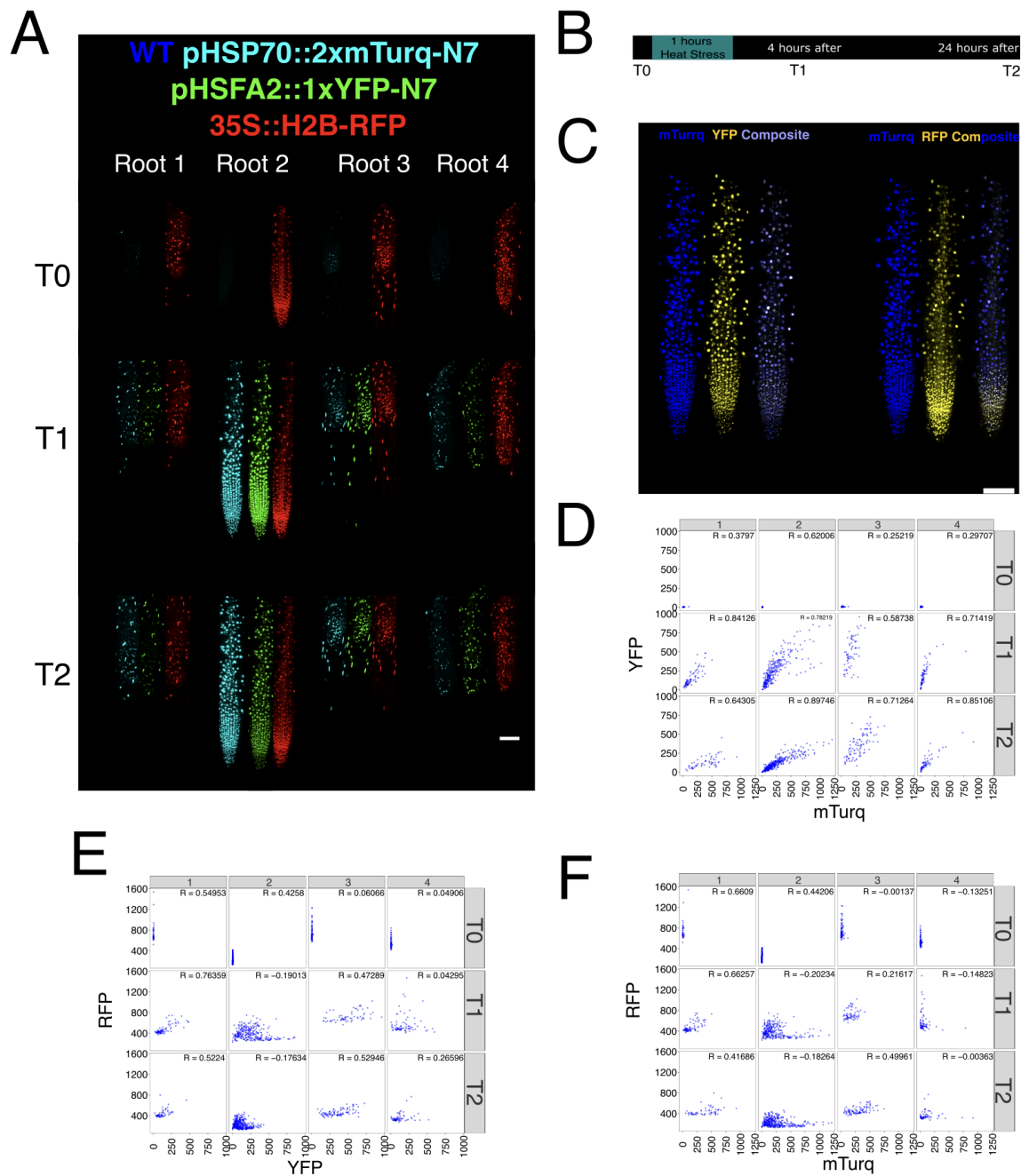

**Fig. S20. *HSFA2* and *HSP70* reporters show coordinated induction across individual nuclei after heat stress.**

**(A)** Representative confocal images of four independently transformed T1-generation seedlings carrying *pHSFA2::1xYFP-N7* in a *pHSP70::2xmTurq-N7*; *35S::H2B-RFP* background. Seedlings were imaged before heat shock (T0), 4 h after a 1 h heat shock at 35 °C (T1 imaging time point), and 24 h after heat shock (T2). Each seedling represents an independent transformation event. Display range: 180-800.

**(B)** Experimental timeline. Seedlings were imaged before heat shock, exposed to 35 °C for 1 h, and re-imaged 4 h and 24 h later.

**(C)** Representative false-coloured images showing overlap between *pHSP70::2xmTurq-N7* and *pHSFA2::1xYFP-N7* signals, and between

*pHSP70::2xmTurq-N7* and *35S::H2B-RFP*, at the 4 h time point. Display ranges: single channels, 205-800; composite images, 205-2000.

**(D)** Pairwise correlations between nuclear fluorescence intensities of *pHSP70::2xmTurq-N7* and *pHSFA2::1xYFP-N7* for the four independently transformed T1-generation seedlings shown in A, at T0, T1 and T2.

**(E, F)** Pairwise correlations between the constitutive nuclear marker *35S::H2B-RFP* and *pHSFA2::1xYFP-N7* (E), or *pHSP70::2xmTurq-N7* (F), for the same seedlings and time points. Pearson correlation coefficients (R) are indicated in each panel.

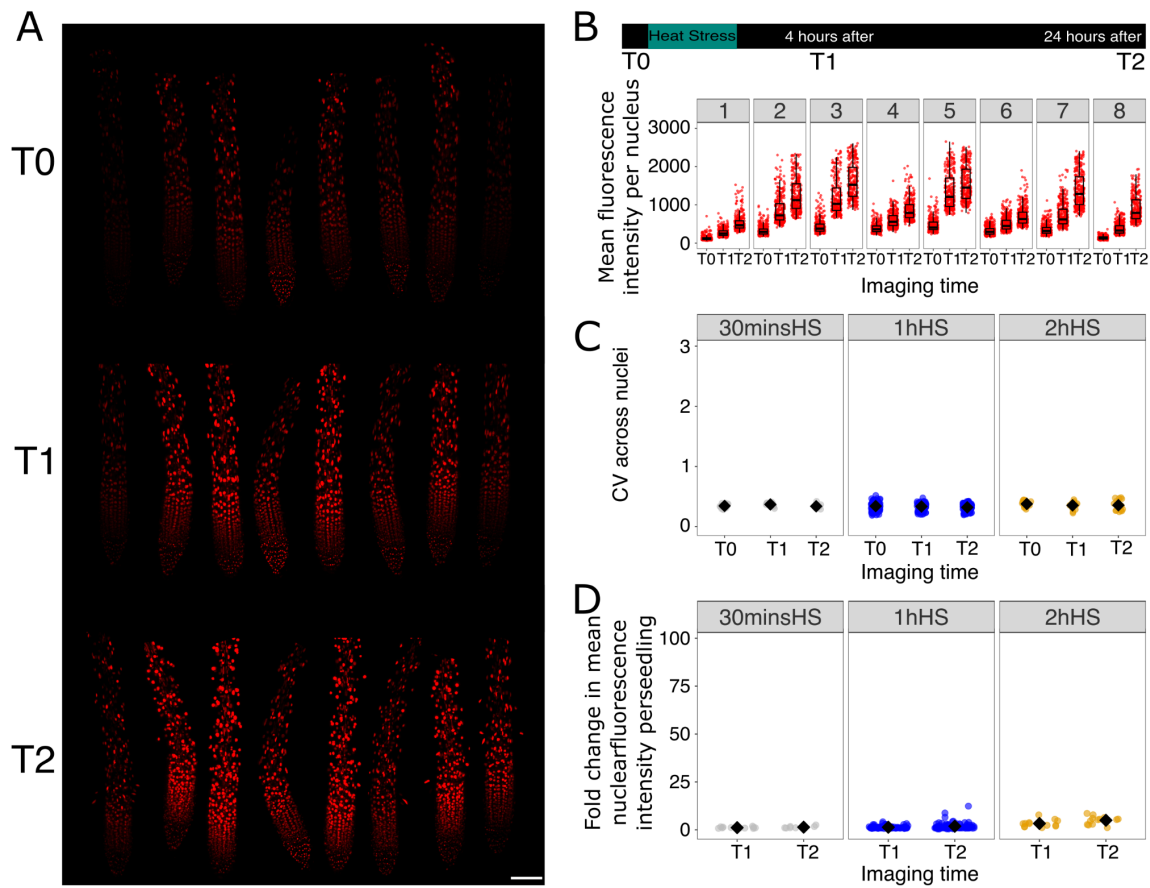

**Fig. S21. *UBQ10::mScarlet-N7* provides an alternative constitutive nuclear marker for *HSP70* reporter imaging experiments**

**(A)** Representative confocal images of *hsfa2* root tips expressing *pHSP70::2xmTurq-N7* and *UBQ10::mScarlet-N7* at T0, T1 and T2.

**(B)** Quantification of nuclear mScarlet intensity in individual seedlings at T0, T1 and T2. Each numbered panel represents one seedling; points show individual nuclei.

**(C)** Coefficient of variation of nuclear mScarlet intensity across nuclei within individual seedlings for the indicated heat stress treatments.

**(D)** Fold change in mean nuclear mScarlet intensity per seedling at T1 and T2 relative to T0. Although *UBQ10::mScarlet-N7* showed some variation across time points, these changes were modest compared with the heat stress-dependent changes observed for the HSP70 reporter and did not reproduce the strong increase in HSP70 heterogeneity.

Data for C and D include *UBQ10::mScarlet-N7* imaging datasets used in the *HSP70* reporter analyses: n = 13, 96 and 11 seedlings from 2, 12 and 2 independent imaging experiments for the 30 min, 1 h and 2 h heat stress treatments, respectively. Each point represents one seedling; black diamonds indicate the mean.

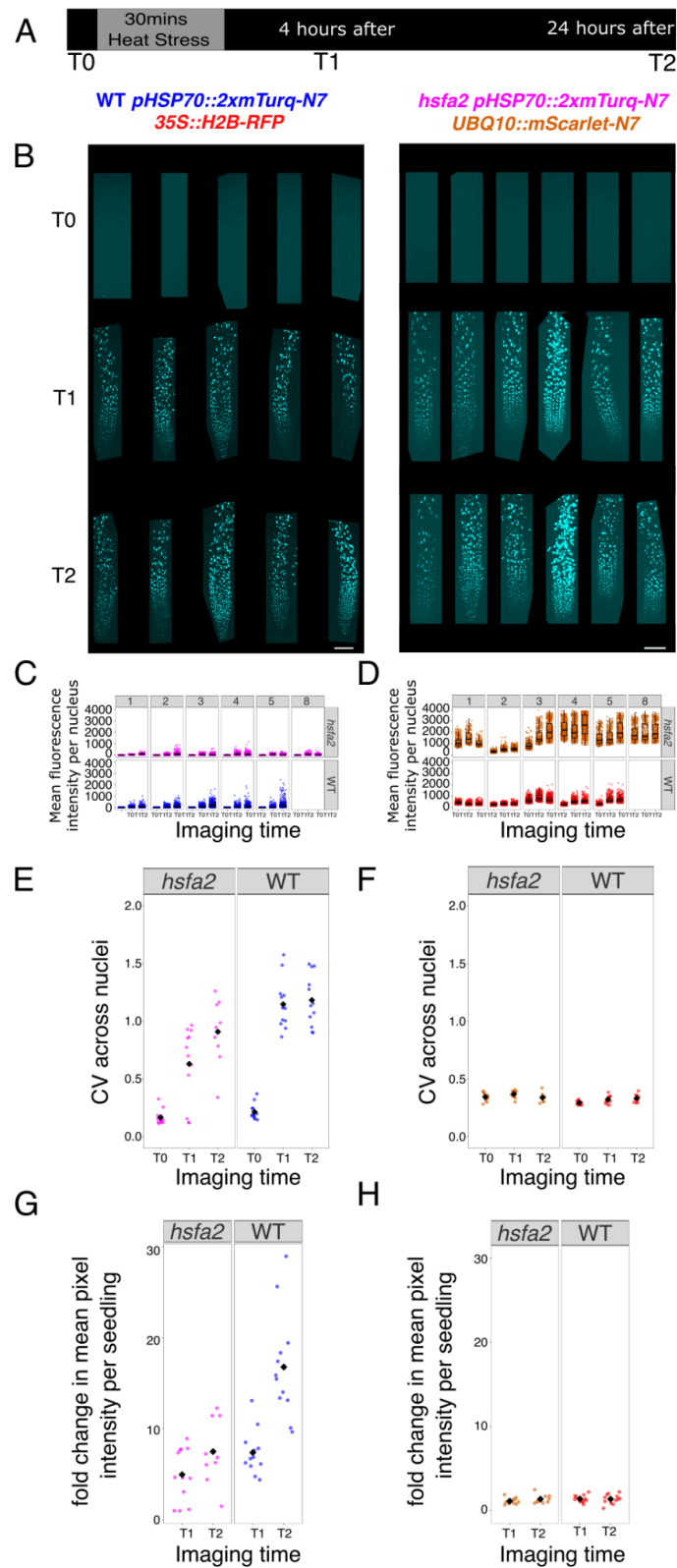

**Fig. S22. *HSFA2* promotes basal *HSP70* reporter expression and broadens the distribution of responses after a 30 min heat shock.**

**(A)** Experimental timeline. Seedlings were imaged before heat stress (T0), exposed to 35 °C for 30 min, and re-imaged 4 h (T1) and 24 h (T2) after heat treatment.

**(B)** Representative confocal images of roots from WT *pHSP70::2xmTurq-N7*;

35S::*H2B-RFP* seedlings and *hsfa2 pHSP70::2xmTurq-N7; UBQ10::mScarlet-N7* seedlings at T0, T1, and T2. In the WT background, low basal *HSP70* reporter activity is detected before stress and heat shock induces a broad, heterogeneous response across nuclei. In the *hsfa2* background, basal reporter activity is highly reduced and the induced response is weaker and more compressed. Display range: WT - T0, 150-350; T1&T2, 150-500 and *hsfA2* - 150-350.

**(C, D)** Quantification of nuclear fluorescence intensities for the *pHSP70::2xmTurq-N7* reporter and constitutive nuclear markers in *hsfa2* and WT seedlings across time points. Each point represents one nucleus.

**(E, F)** Cell-to-cell variability, measured as the coefficient of variation across nuclei within each seedling, for the *pHSP70::2xmTurq-N7* reporter **(E)** and the constitutive nuclear markers **(F)** in *hsfa2* and WT backgrounds. Each point represents one seedling; black diamonds indicate the mean. Loss of *HSFA2* reduces both basal reporter activity and the variability of the heat-induced response. **(G, H)** Fold change in mean nuclear fluorescence intensity per seedling relative to T0 for the *pHSP70::2xmTurq-N7* reporter **(G)** and the constitutive nuclear markers **(H)** in *hsfa2* and WT backgrounds. Each point represents one seedling; black diamonds indicate the mean.

The WT and *hsfa2* 30 min datasets shown here were acquired using the low CFP acquisition settings described in Fig. S4. The WT 30 min dataset is summarised in Fig. 1 and Fig. S4 and replotted as the WT comparator for the *hsfa2* 30 min analysis in Fig. 5. The *hsfa2* 30 min dataset shown here is summarised in Fig. 5. For (E-H), data are pooled from biological repeats: WT, n = 12 seedlings from 2 independent imaging experiments, *hsfa2*, n = 13 seedlings from 2 independent imaging experiments (10 seedlings for T2 timepoint).

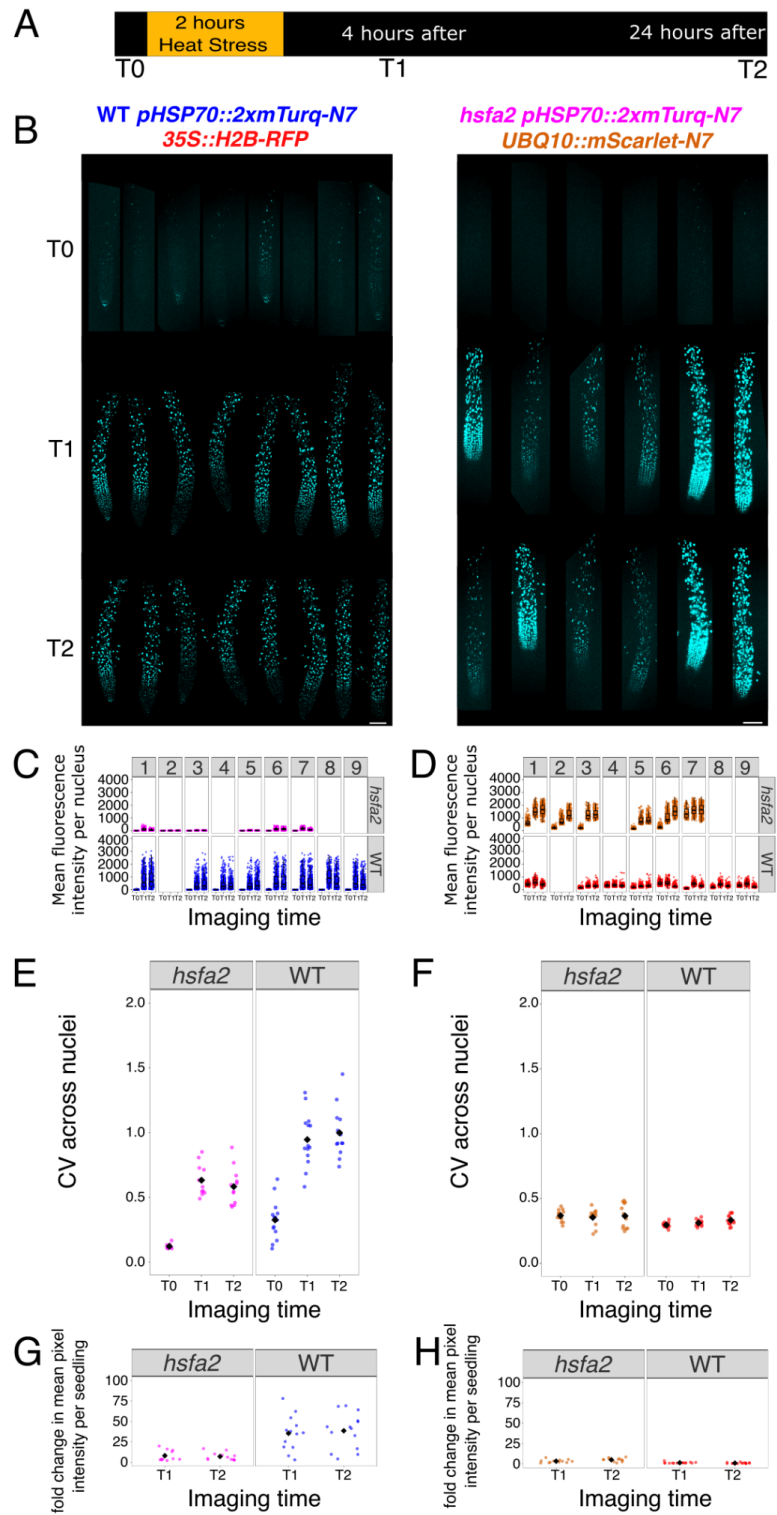

**Fig. S23. *HSFA2* promotes basal *HSP70* reporter expression and broadens the distribution of responses after a 2 h heat shock.**

**(A)** Experimental timeline. Seedlings were imaged before heat stress (T0), exposed to 35 °C for 2 h, and re-imaged 4 h (T1) and 24 h (T2) after heat treatment.

**(B)** Representative confocal images of roots from WT *pHSP70::2xmTurq-N7*; *35S::H2B-RFP* seedlings and *hsfa2-1* *pHSP70::2xmTurq-N7*; *UBQ10::mScarlet-N7* seedlings at T0, T1, and T2. In the WT background, low basal *HSP70* reporter activity is detected before stress and heat shock induces a broad, heterogeneous response across nuclei. In the *hsfa2* background, basal reporter activity is reduced and the induced response is weaker and more compressed. Display range: WT - T0, 200-250; T1&T2, 200-2000 and *hsfA2* - 200-250.

**(C, D)** Quantification of nuclear fluorescence intensities for the *pHSP70::2xmTurq-N7* reporter and constitutive nuclear markers in *hsfa2-1* and WT seedlings across time points. Each point represents one nucleus.

**(E, F)** Cell-to-cell variability, measured as the coefficient of variation across nuclei within each seedling, for the *pHSP70::2xmTurq-N7* reporter **(E)** and the constitutive nuclear markers **(F)** in *hsfa2-1* and WT backgrounds. Each point represents one seedling; black diamonds indicate the mean. Loss of *HSFA2* reduces both basal reporter activity and the variability of the heat-induced response.

**(G, H)** Fold change in mean nuclear fluorescence intensity per seedling relative to T0 for the *pHSP70::2xmTurq-N7* reporter **(G)** and the constitutive nuclear markers **(H)** in *hsfa2-1* and WT backgrounds. Each point represents one seedling; black diamonds indicate the mean.

The WT 2 h dataset is summarised in Fig. 1 and Fig. S5 and replotted as the WT comparator for the *hsfa2* 2 h analysis in Fig. 5. The *hsfa2* 2 h dataset shown here is summarised in Fig. 5. For (E-H), data are pooled from biological repeats: WT, n = 15 seedlings from 2 independent imaging experiments, *hsfa2*, n = 11 seedlings from 2 independent imaging experiments.

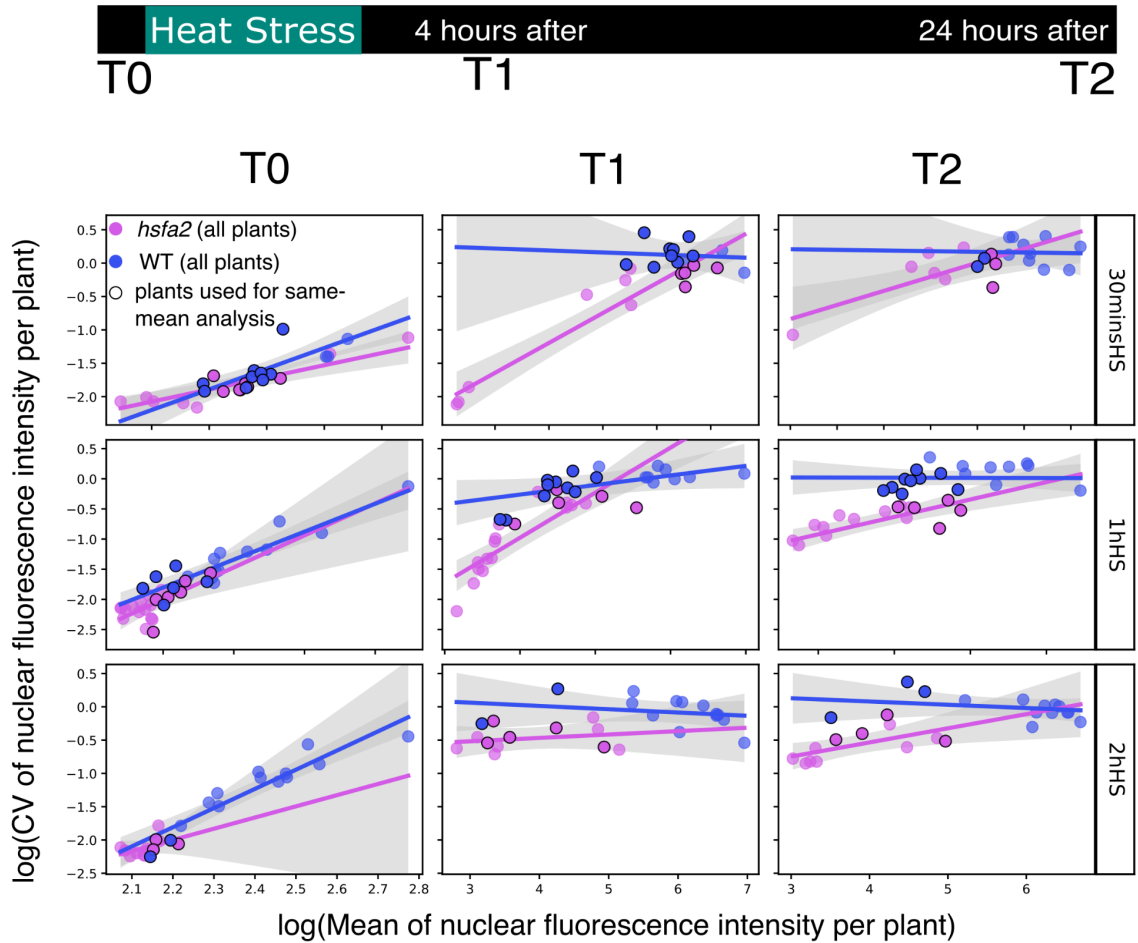

**Fig. S24. Mean-adjusted HSP70 reporter variability in WT and *hsfa2*.**

Blue and magenta lines show genotype-specific regressions fitted across all roots within each heat-treatment and imaging-time panel, with grey ribbons indicating 95% confidence intervals; experiment was included as a blocking factor in these descriptive fits. Black outlines indicate roots that fall within the WT-*hsfa2* mean fluorescence overlap range in their own experiment and therefore contribute to the mean-adjusted analysis. The treatment-specific panels are shown to visualise the data structure and genotype-specific mean-CV relationships; the primary statistical analysis pooled roots across the 30 min, 1 h and 2 h heat-shock treatments and tested genotype effects separately at 4 h and 24 h after heat treatment. Each point represents one root, with mean nuclear CFP fluorescence plotted against the coefficient of variation (CV) across nuclei. All roots are shown.

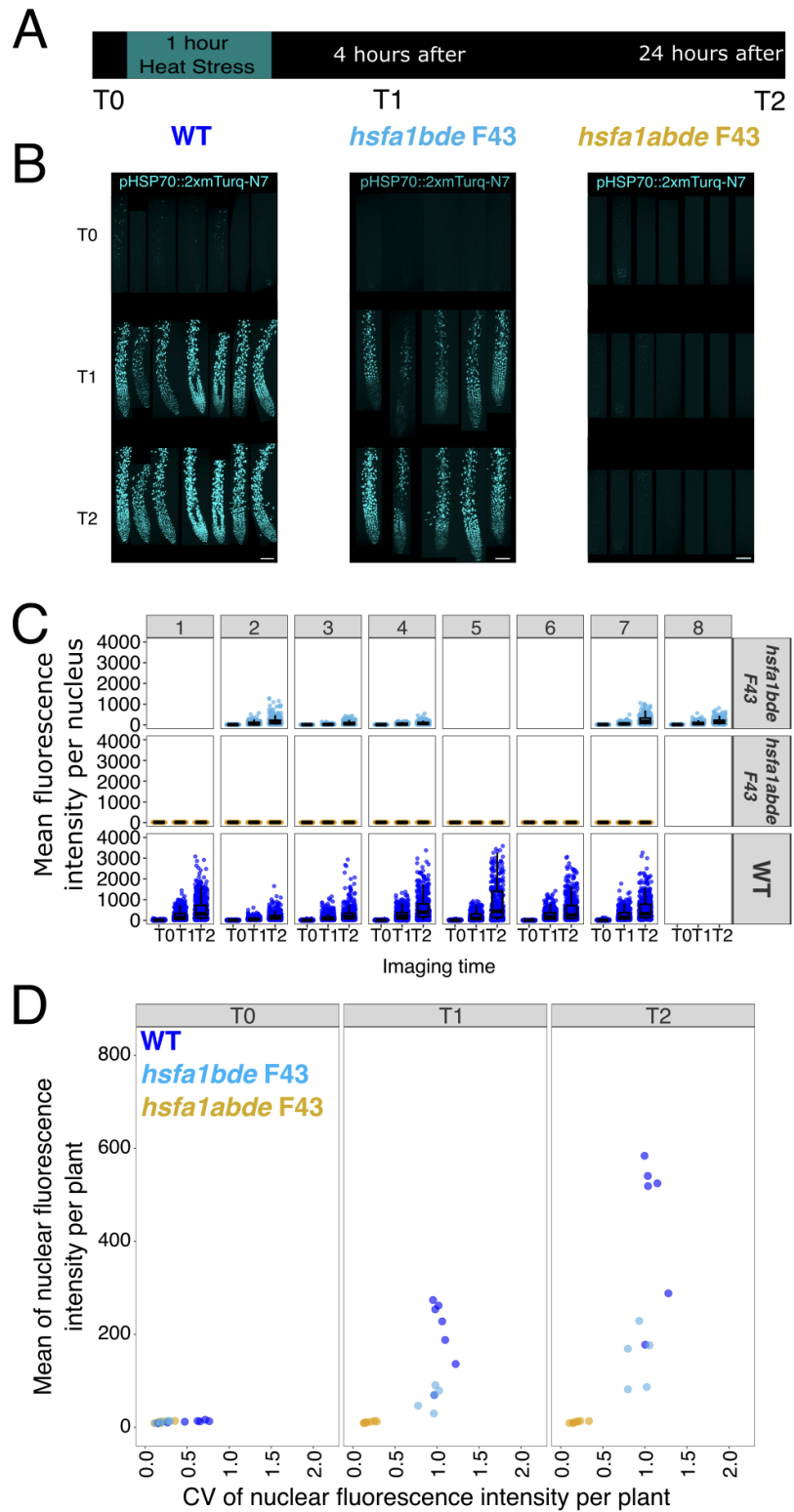

**Fig. S25. Sibling *hsfa1* triple- and quadruple-mutant lines support strongly reduced *HSP70* induction in *hsfa1abde*.**

**(A)** Imaging scheme. Seedlings were imaged before heat shock (T0), exposed to 35 °C for 1 h, and re-imaged 4 h (T1) and 24 h (T2) after heat shock.

**(B)** Representative confocal images of root tips expressing *pHSP70::2xmTurq-N7* in WT, *hsfa1bde* F43 and *hsfa1abde* F43 seedlings at T0, T1 and T2. F43 indicates sibling lines derived from the same segregating family, allowing comparison of related genetic backgrounds that differ in the presence or absence of a wild-type *HSFA1a* allele (Methods). Images are shown with the same display range: 180–400.

**(C)** Quantification of nuclear *HSP70* reporter fluorescence in individual seedlings at T0, T1 and T2. Each numbered panel represents one seedling; points represent individual nuclei.

**(D)** Mean nuclear *HSP70* reporter fluorescence plotted against the coefficient of variation across nuclei within individual seedlings. Each point represents one seedling. WT seedlings showed strong heat-induced reporter activation, *hsfa1bde* F43 seedlings retained reporter induction, and *hsfa1abde* F43 seedlings showed strongly reduced heat-induced reporter activation.

Data used in (B-D) is from WT, $n=7$; *hsfa1bde*, $n = 5$; *hsfa1abde*, $n = 7$ seedlings from 1 imaging experiment.

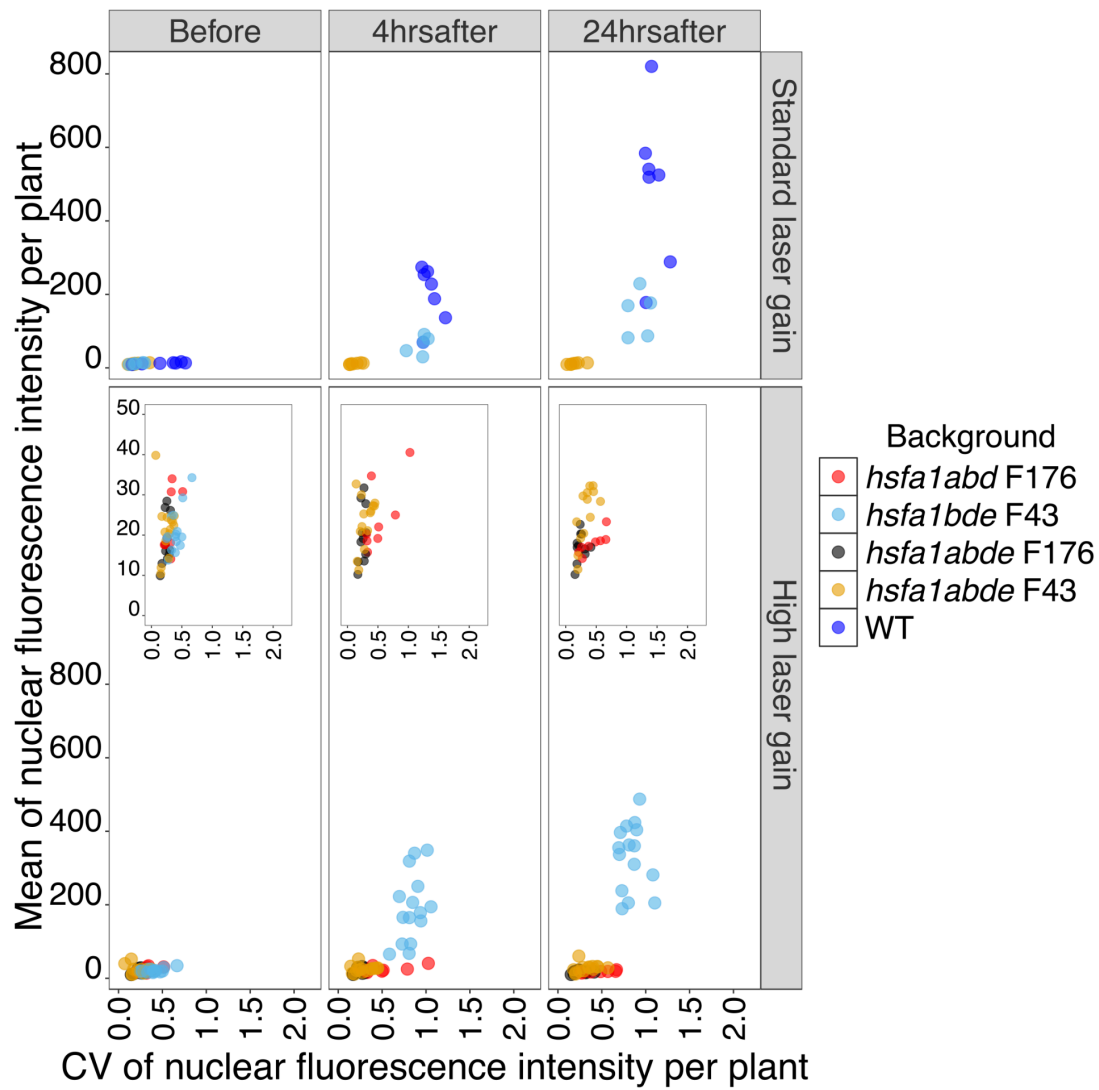

**Fig. S26. Independent *hsfa1abde* families and high-gain imaging support strongly reduced *HSP70* induction in *hsfa1abde*.**

Mean nuclear *HSP70* reporter fluorescence is plotted against the coefficient of variation across nuclei within individual seedlings before heat shock, 4 h after heat shock, and 24 h after heat shock. Top, data acquired using the standard gain used for the main *hsfa1* comparisons. Under these settings, *HSP70* reporter signal in *hsfa1abde* F43 and *hsfa1abde* F176 seedlings was close to background, whereas WT and *hsfa1bde* F43 seedlings showed clear heat-induced reporter activation. Bottom, data acquired using higher-gain settings to test whether low-level *HSP70* reporter signal could be detected in weakly expressing backgrounds. Insets show the expanded low-intensity range. High-gain imaging revealed faint reporter signal in *hsfa1abde* F43, *hsfa1abde* F176 and *hsfa1abd* F176 seedlings, but signal remained much lower than in WT and *hsfa1bde* F43 and showed little heat-induced increase. F43 and F176 indicate lines derived from two independent segregating families in the mixed Ws/Col-0 *hsfa1* background; these families were used to test whether reduced *HSP70* induction in *hsfa1abde* was reproducible and not explained by family-specific background effects.

Data used for standard laser-gain is from WT, $n=7$; *hsfa1bde*, $n=5$; *hsfa1abde*, $n=7$ seedlings from 1 experiment. Data used here for high laser-gain are from,

*hsfa1bde* family 43, n = 15 seedlings from 2 independent experiments ; *hsfa1abd* family 176, n = 8 seedlings from 2 independent experiments; *hsfa1abde* family 43, n = 15 seedlings from 2 independent experiments; *hsfa1abde* family 176, n = 11 seedlings from 2 independent experiments.

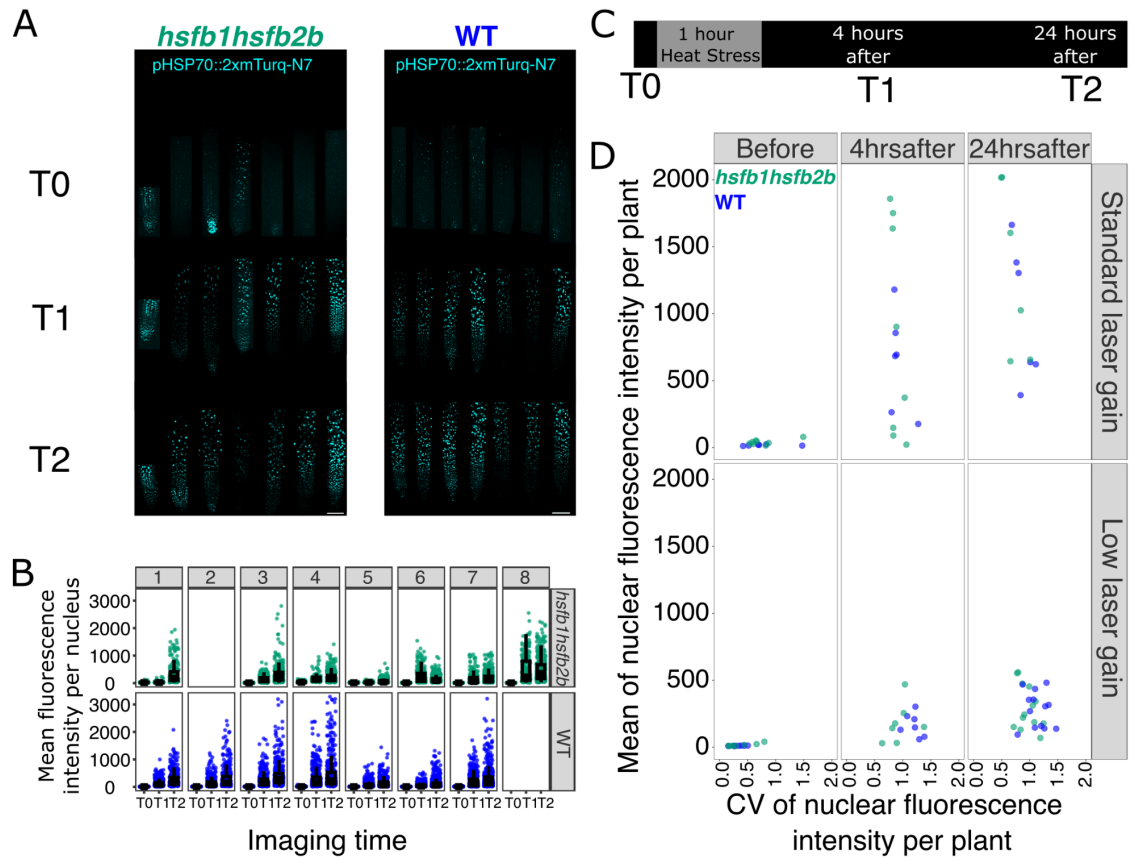

**Fig. S27. Loss of *HSFB1/HSFB2b* does not strongly alter heterogeneous *HSP70* activation after heat stress.**

**(A)** Representative confocal images of *hsfb1hsfb2b* and WT root tips expressing pHSP70::2xmTurq-N7 at T0 (before heat stress), T1 (4 h after heat stress), and T2 (24 h after heat stress). Basal reporter expression was detectable in both genotypes before heat stress, and reporter expression increased after a 1 h heat stress treatment. Display range: T0, 200-250; T1 and T2, 200-1500.

**(B)** Quantification of nuclear mTurquoise intensity in WT and *hsfB1hsfB2b* seedlings at T0, T1 and T2. Each point represents a single nucleus.

**(C)** Imaging scheme. Seedlings were imaged before heat stress (T0), then re-imaged 4 h (T1) and 24 h (T2) after a 1 h heat stress treatment.

**(D)** Mean pixel intensity plotted against the coefficient of variation (CV) of pixel intensity for individual seedlings at each time point. Top, images acquired using identical laser gain across samples. Bottom, the same roots imaged at lower laser gain to avoid possible saturation in strongly expressing samples. WT and *hsfB1hsfB2b* seedlings showed similar mean-CV relationships under both imaging conditions, indicating that loss of *HSFB1/HSFB2b* does not strongly affect *HSP70* expression heterogeneity.

Data used for (D) for low laser-gain are from, *hsfb1hsfb2b* n = 15 seedlings from 2 independent experiments ; WT, n = 13 seedlings from 2 independent experiments. For standard laser-gain data in (D), the same plants of one experiment with low laser-gain were reimaged, *hsfb1hsfb2b* n = 8 seedlings ; WT, n = 6 seedlings.

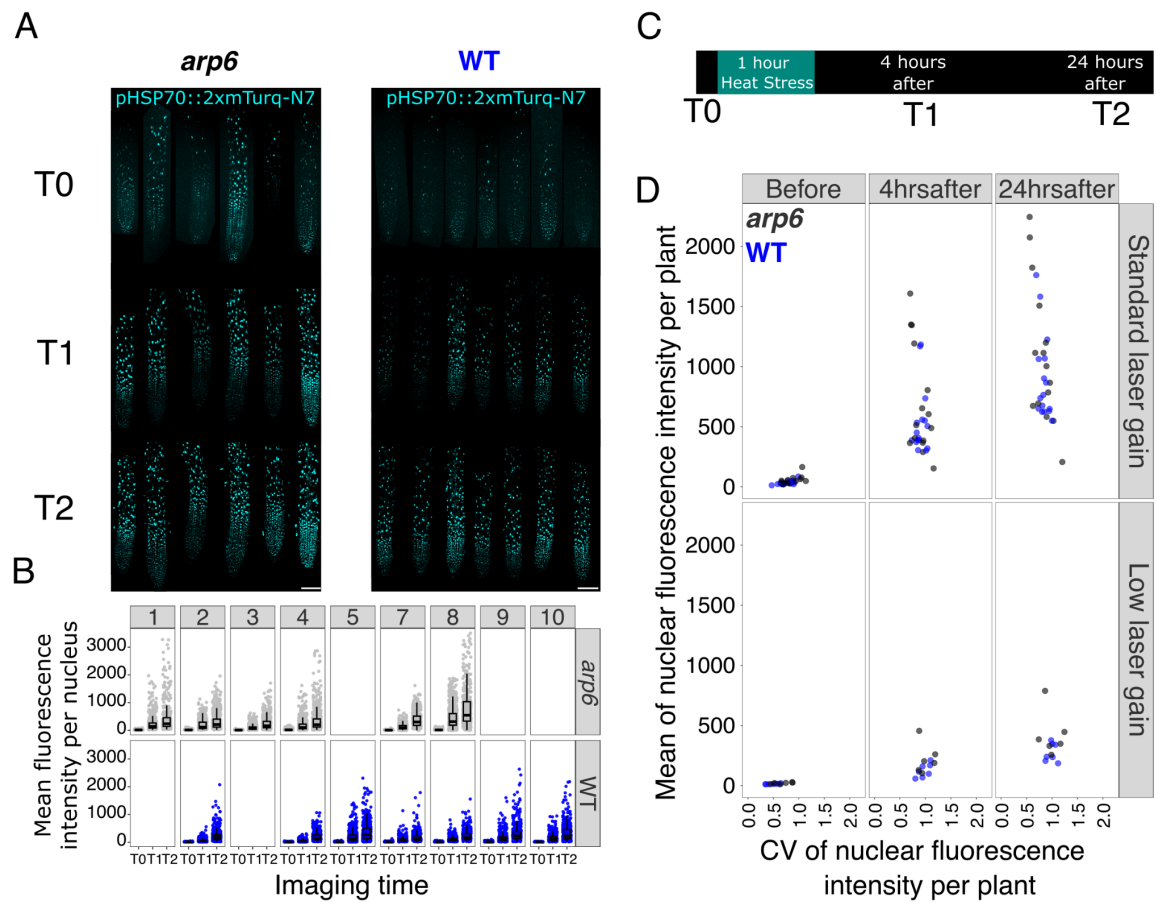

**Fig. S28. *arp6* elevates *pHSP70* expression without abolishing heat-shock heterogeneity.**

**(A)** Representative confocal images of *arp6* and WT root tips expressing *pHSP70::2xmTurq-N7* at T0 (before heat stress), T1 (4 h after heat stress), and T2 (24 h after heat stress). Display range: T0, 200-250; T1 and T2, 200-1000.

**(B)** Quantification of reporter pixel intensity in individual seedlings at T0, T1 and T2. Each numbered panel represents an individual seedling. Boxplots summarise the distribution of pixel intensities within each seedling, with points showing individual measurements. WT, blue; *arp6*, black.

**(C)** Imaging scheme. Seedlings were imaged before heat stress (T0), then re-imaged 4 h (T1) and 24 h (T2) after a 1 h heat stress treatment.

**(D)** Mean pixel intensity plotted against the coefficient of variation (CV) of pixel intensity for individual seedlings at each time point. Top, images acquired using standard laser gain. Bottom, images acquired using lower laser gain. WT, blue; *arp6*, black. *arp6* increased reporter expression, but substantial heterogeneity was retained after heat stress.

Data used for (D) standard laser-gain are from, *arp6* $n = 15$ seedlings from 2 independent experiments; WT, $n = 16$ seedlings from 2 independent experiments for standard laser. For low laser-gain data in (D), the same plants of one experiment with standard laser-gain were reimaged, *arp6* $n = 6$ seedlings; WT, $n = 7$ seedlings.

**Table S1.** Primers used for construct generation.

| Promoter | Fwd Primer | Rvs primer | Product size |
| --- | --- | --- | --- |
| HSP70 | attB4-pHSP70-F<br>GGGGACAACTTT<br>GTATAGAAAAGT<br>TGCTGATTAGTG<br>TGA CTGAACC | attB1R-pHSP70-R<br>GGGGACTGCTT<br>TTTTGTACAAAC<br>TTGCTATTAGAG<br>ATCAGAATTG | 2104 |
| HSFA2 | attB4-pHSFA2-F<br>GGGGACAACTTT<br>GTATAGAAAAGT<br>TGCCCAATCAAA<br>TCTCTCCTTCAC<br>G | attB1R-pHSFA2-R<br>GGGGACTGCTT<br>TTTTGTACAAAC<br>TTGCCTCAAATC<br>CATAAGCTCAGA<br>G | 716 |

**Table S2.** Primers used for genotyping.

| Gene | PCR detects presence of WT/ Mutant allele | Fwd primer | Rvs primer | Product size |
| --- | --- | --- | --- | --- |
| HSFA1a | WT | hsfA1a_F_GTCC<br>AGGTGATGAAA<br>ATGTGTTCAAG<br>CAA | hsfA1a_R_ATGA<br>AGAGCTCGCAA<br>GGACCTCCATT<br>GAA | ~2.2kb |
| HSFA1a | Mutant | hsfA1a_F_GTCC<br>AGGTGATGAAA<br>ATGTGTTCAAG<br>CAA | hsfA1a_TDNALB<br>_CATTTTATAATA<br>ACGCTGCGGA<br>CATCTAC | ~2.2kb |
| HSFA1b | WT | hsfA1b_F2_CCA<br>ATGAGGTCGTT<br>TCGTGG | hsfA1b_R2_GTC<br>CCACATTCTGC<br>AGTTGG | ~1kb |
| HSFA1b | Mutant | hsfA1b_TDNALB<br>_GATGCACTCG<br>AAATCAGCCAA<br>TTTGTAGAC | hsfA1b_R2_GTC<br>CCACATTCTGC<br>AGTTGG | ~750bp |
| HSFA1d<br>(SALK_02240<br>4C) | WT | hsfA1d_F_CCG<br>AAACATTTTCA<br>GGAGTTG | hsfA1d_R_ATGT<br>TGGGCACTATT<br>TGAAGC | 1206bp |
| HSFA1d<br>(SALK_02240<br>4C) | Mutant | LBb1.3<br>ATTTTGCCGAT<br>TTCGGAAC | hsfA1d_R_ATGT<br>TGGGCACTATT<br>TGAAGC | ~600bp |
| HSFA1e<br>(SALK_09494<br>3) | WT | hsfA1e_F_CGAA<br>CTTTGCAGGCT<br>TATTTG | hsfA1e_R_TCCA<br>TTATCTGTCTCG<br>CATCC | 1250bp |
| HSFA1e<br>(SALK_09494<br>3) | Mutant | LBb1.3<br>ATTTTGCCGAT<br>TTCGGAAC | hsfA1e_R_TCCA<br>TTATCTGTCTCG<br>CATCC | ~750bp |
| HSFA2<br>(SALK_00897<br>8C) | WT | hsfa2_F_AAGGT<br>TCCGAACCAA<br>GAAAAC | hsfa2_43.05_R_<br>TCCTTCCACGT<br>TACTTCAAGC | 1100bp |
| HSFA2<br>(SALK_00897<br>8C) | Mutant | LBb1.3<br>ATTTTGCCGAT<br>TTCGGAAC | hsfa2_43.05_R_<br>TCCTTCCACGT<br>TACTTCAAGC | ~750bp |
| HSFb2b-1<br>(SALK_047291) | WT-1 | HSFB2B-1_F_C<br>TGAATCGGCAA<br>GTGTTTTTC | HSFB2B-1_R_A<br>AGTAGTGGATG<br>TGGTGCTGG | 1155bp |
| HSFb2b-1<br>(SALK_047291) | Mutant-1 | LBb1.3<br>ATTTTGCCGAT<br>TTCGGAAC | HSFB2B-1_R_A<br>AGTAGTGGATG<br>TGGTGCTGG | ~700bp |
| HSFb2b-1<br>(SALK_047291) | WT-2 (off<br>target<br>insertion) | HSFB2B-1_2ndIn<br>sertionF_GTTG<br>ACGCTCCTTGT<br>TTTCAC | HSFB2B-1_2ndIn<br>sertionR_TTCGG<br>ACCTTAATTTTC<br>TTATATCG | 1184 |
| HSFb2b-1<br>(SALK_047291) | Mutant-2<br>(off target<br>insertion) | LBb1.3<br>ATTTTGCCGAT<br>TTCGGAAC | HSFB2B-1_2ndIn<br>sertionR_TTCGG<br>ACCTTAATTTTC<br>TTATATCG | ~700bp |
| HSFb2b-1<br>(SALK_047291) | WT-3 (off<br>target<br>insertion) | HSFB2B-1_3rdIn<br>sertionF_TCTAG<br>CCTCACCCAGC<br>AAGAAG | HSFB2B-1_3rdIn<br>sertionR_TGCAG<br>GTTGGTTTGA<br>GATAG | 1055bp |
| HSFb2b-1<br>(SALK_047291) | Mutant-3<br>(off target<br>insertion) | LBb1.3<br>ATTTTGCCGAT<br>TTCGGAAC | HSFB2B-1_3rdIn<br>sertionR_TGCAG<br>GTTGGTTTGA<br>GATAG | ~600bp |
| HSFb1-1<br>(SALK_104713C) | WT | HSFB1-1_F_CA<br>GACTTTGCTGC<br>TTTTCCAC | HSFB1-1_R_AC<br>TCCGATCTTCT<br>CCGTTCTC | 1238bp |
| HSFb1-1<br>(SALK_104713C) | Mutant | LBb1.3<br>ATTTTGCCGAT<br>TTCGGAAC | HSFB1-1_R_AC<br>TCCGATCTTCT<br>CCGTTCTC | ~700bp |
| arp6<br>(SAIL_236_C07) | WT | arp6-2-S<br>(5'-GACGTTATT<br>CCAGCCTGCA<br>GATTTA-3') | arp6 Rvs K2<br>CGAGCTGATCC<br>AAGTTCTTCG | ~750bp |
| arp6<br>(SAIL_236_C07) | Mutant | arp6-2-S<br>(5'-GACGTTATT<br>CCAGCCTGCA<br>GATTTA-3') | LB3<br>(5'-TAGCATCTG<br>AATTTTCATAACC<br>AATCTCGATACA<br>C-3') | ~700bp |

**Table S3.** Modelling parameters.

| Parameter | Value | Definition |
| --- | --- | --- |
| a1 | 5 | Max activation strength for A1 self-activation |
| a2 | 10 | A1-dependent activation strength for P70 |
| a3 | 75 | A2-dependent activation strength for P70 |
| a4 | 60 | A1-dependent activation strength for A2 |
| a5 | 15 | A1-dependent activation strength for B |
| a6 | 2 | Refolding rate coefficient (from $C\{P70 \cdot M\}$) |
| a7 | 300 | Basal production rate of O |
| h1 | 10 | Hill threshold for A1 self-activation |
| h2 | 5 | Hill threshold for A1→P70 regulation |
| h3 | 8 | Hill threshold for A2→P70 regulation |
| h4 | 12 | Hill threshold for A1→A2 regulation |
| h5 | 15 | Hill threshold for A1→B regulation |
| n | 2 | Hill coefficient (cooperativity exponent) |
| leakage_A1 | 0.1 | Basal expression ("leakage") of A1 |
| leakage_P70 | 1.75 | Basal expression ("leakage") of P70 |
| leakage_A2 | 0.01 | Basal expression ("leakage") of A2 |
| leakage_B | 0.01 | Basal expression ("leakage") of B |
| c1 | 10 | Association rate: $A1 + P70 \rightarrow C\{A1 \cdot P70\}$ |
| c3 | 15 | Association rate: $P70 + M \rightarrow C\{P70 \cdot M\}$ |
| c2 | 5 | Displacement rate: $C\{A1 \cdot P70\} + M \rightarrow C\{P70 \cdot M\}$ |
| c4 | 1.6 | Reverse displacement: $C\{P70 \cdot M\} + A1 \rightarrow C\{A1 \cdot P70\}$ |
| d1 | 1.35 | Dissociation rate of $C\{A1 \cdot P70\}$ |
| d3 | 2.5 | Dissociation rate of $C\{P70 \cdot M\}$ |
| d | 0.04 | Common decay/dilution rate |
| d5 | 0.1 | Decay rate of misfolded proteins pool (M) |
| d4 (normal) | 0.01 | Misfolding rate of O under normal conditions |
| d4 (heat) | 0.05 | Misfolding rate of O under heat shock |
| init_A1 | 1 | Initial value of A1 |
| init_P70 | 35 | Initial value of P70 |
| init_C_A1_P70 | 5 | Initial value of $C\{A1 \cdot P70\}$ |
| init_M | 0 | Initial value of M |
| init_O | 7500 | Initial value of O |
| init_C_P70_M | 35 | Initial value of $C\{P70 \cdot M\}$ |
| init_A2 | 1 | Initial value of A2 |
| init_B | 1 | Initial value of B |

**Table S4. Statistical analysis of CV difference between genotypes**

| Time point | n_roots | n_WT | n_hsfA2 | n_experiments | *hsfa2*_WT_CV_ratio_at_central_mean | CI95_low | CI95_high | p_genotype_at_central_mean | p_join_t_genotype_terms | p_mean_by_genotype_interaction |
| --- | --- | --- | --- | --- | --- | --- | --- | --- | --- | --- |
| T1 | 36 | 21 | 15 | 5 | 0.718 | 0.622 | 0.828 | 5.432x10<sup>-5</sup> | 3.92 x10<sup>-5</sup> | 0.00881 |
| T2 | 26 | 14 | 12 | 5 | 0.655 | 0.562 | 0.763 | 1.630x10<sup>-5</sup> | 6.89 x10<sup>-5</sup> | 0.714 |

**Table S5. Marker list**

| Cluster | Gene ID | Gene name | Annotation | Reference |
| --- | --- | --- | --- | --- |
| 1, 7, 23 | AT3G09220 | LAC7 | LRC & epidermis | (22) |
| 1, 7, 23 | AT5G44610 | MAP18 | LRC & epidermis | (23) |
| 1, 7, 23 | AT1G79270 | ECT8 | LRC & epidermis | (24) |
| 1, 7, 23 | AT5G24270 | SOS3 | LRC & epidermis | (25) |
| 1, 7, 23 | AT1G56010 | NAC1 | LRC & epidermis | (26) |
| 2 | AT3G44310 | NIT1 | Meristem epidermis | (27) |
| 2 | AT1G65970 | TPX2 | Meristem epidermis | (28) |
| 2 | AT4G23700 | CHX17 | Meristem epidermis | (29) |
| 3 | AT4G30450 | XPP | Xylem & XPP | (30) |
| 3 | AT2G36120 | DOT1 | Xylem & XPP | (31) |
| 4 | AT1G45201 | TLL1 | Cortex maturation | (32) |
| 4 | AT1G11130 | SCM | Cortex maturation | (32, 33) |
| 5 | AT5G57130 | SMXL5 | Procambium | (34) |
| 5 | AT1G04240 | SHY2 | Procambium | (31) |
| 8, 15 | AT2G48130 | AT2G48130 | Endodermis & CS | (35) |
| 8, 15 | AT1G61590 | AT1G61590 | Endodermis & CS | (35) |
| 8 | AT2G36100 | CASP1 | Endodermis & CS | (36) |
| 9 | AT5G12330 | LRP1 | Lateral root primordia | (37) |
| 9 | AT2G38530 | LTP2 | Lateral root primordia | (38) |
| 11 | AT1G80240 | DGR1 | Root cap elongation | (39) |
| 11 | AT5G54510 | DFL1/GH3.6 | Root cap elongation | (40) |
| 12 | AT1G79840 | GL2 | Non-hair | (41, 42) |
| 12 | AT2G46410 | CPC | Non-hair | (43) |
| 13 | AT2G36620 | RPL24A | Meristem | (44) |
| 13 | AT1G69620 | RPL34B | Meristem | (44) |
| 13 | AT5G14750 | WER | Meristem | (45) |
| 14 | AT1G12950 | RHS2 | Root-hair | (46) |
| 14 | AT3G54420 | EP3 | Root-hair | (47) |
| 16 | AT4G29100 | BHLH68 | Pericycle | (31) |
| 16 | AT5G09760 | AT5G09760 | Pericycle | (31) |
| 17 | AT2G22850 | bZIP6 | Phloem | (48) |
| 17 | AT4G37730 | bZIP7 | Phloem | (48) |
| 17 | AT5G24800 | BZIP9 | Phloem | (48) |
| 18 | AT1G62510 | AT1G62510 | Cortex initials | (35, 49) |
| 18 | AT5G03150 | JKD | Cortex initials | (45) |
| 19 | AT1G22710 | SUC2 | Companion cells | (50) |
| 19 | AT5G02600 | NAKR1 | Companion cells | (51) |
| 20 | AT3G54220 | SCR | Endodermis initials | (52) |
| 20 | AT5G57620 | MYB36 | Endodermis initials | (53, 54) |
| 21 | AT3G25710 | TMO5 | Protoxylem & metaxylem | (42, 55) |
| 21 | AT5G19530 | ACL5 | Protoxylem & metaxylem | (31, 56, 57) |
| 22 | AT1G11190 | BFN1 | LRC | (58) |

### Legends for Datasets

**Supplementary Dataset S1** DGE analysis results for 22 °C

**Supplementary Dataset S2** DGE analysis results for 35 °C

**Supplementary Dataset S3** Gene Ontology enrichment results for 22 °C

**Supplementary Dataset S4** Gene Ontology enrichment results for 35 °C

**Supplementary Dataset S5** Lists of genes recurrently upregulated in ON nuclei at 22 °C and 35 °C, and the 15-gene signature used for UCell scoring. Bold entries identify genes included in the UCell signature.

### SI References:

1. F. Federici, L. Dupuy, L. Laplace, M. Heisler, J. Haseloff, Integrated genetic and computation methods for in planta cytometry. *Nat Methods* **9**, 483–485 (2012).
2. Y. Charng, *et al.*, A Heat-Inducible Transcription Factor, HsfA2, Is Required for Extension of Acquired Thermotolerance in Arabidopsis. *Plant Physiol* **143**, 251–262 (2007).
3. M. Ikeda, N. Mitsuda, M. Ohme-Takagi, Arabidopsis HsfB1 and HsfB2b Act as Repressors of the Expression of Heat-Inducible Hsfs But Positively Regulate the Acquired Thermotolerance. *Plant Physiol* **157**, 1243–1254 (2011).
4. H.-C. Liu, H.-T. Liao, Y.-Y. Charng, The role of class A1 heat shock factors (HSFA1s) in response to heat and other stresses in Arabidopsis. *Plant, Cell & Environment* **34**, 738–751 (2011).
5. S. Mlotshwa, *et al.*, Transcriptional silencing induced by Arabidopsis T-DNA mutants is associated with 35S promoter siRNAs and requires genes involved in siRNA-mediated chromatin silencing. *The Plant Journal* **64**, 699–704 (2010).
6. R. B. Deal, M. K. Kandasamy, E. C. McKinney, R. B. Meagher, The Nuclear Actin-Related Protein ARP6 Is a Pleiotropic Developmental Regulator Required for the Maintenance of FLOWERING LOCUS C Expression and Repression of Flowering in Arabidopsis. *Plant Cell* **17**, 2633–2646 (2005).
7. G. X. Y. Zheng, *et al.*, Massively parallel digital transcriptional profiling of single cells. *Nat Commun* **8**, 14049 (2017).
8. C.-Y. Cheng, *et al.*, Araport11: a complete reannotation of the Arabidopsis thaliana reference genome. *The Plant Journal* **89**, 789–804 (2017).
9. C. S. McGinnis, L. M. Murrow, Z. J. Gartner, DoubletFinder: Doublet Detection in Single-Cell RNA Sequencing Data Using Artificial Nearest Neighbors. *Cell Systems* **8**, 329–337.e4 (2019).
10. Y. Hao, *et al.*, Dictionary learning for integrative, multimodal and scalable single-cell analysis. *Nat Biotechnol* **42**, 293–304 (2024).

11. E. J. Blair, *et al.*, Contribution of time of day and the circadian clock to the heat stress responsive transcriptome in Arabidopsis. *Sci Rep* **9**, 4814 (2019).
12. C. Hafemeister, R. Satija, Normalization and variance stabilization of single-cell RNA-seq data using regularized negative binomial regression. *Genome Biol* **20**, 296 (2019).
13. R. Shahan, *et al.*, A single-cell Arabidopsis root atlas reveals developmental trajectories in wild-type and cell identity mutants. *Developmental Cell* **57**, 543-560.e9 (2022).
14. T. A. Lee, *et al.*, A single-cell, spatial transcriptomic atlas of the Arabidopsis life cycle. *Nat. Plants* **11**, 1960–1975 (2025).
15. M. Andreatta, S. J. Carmona, UCell: Robust and scalable single-cell gene signature scoring. *Computational and Structural Biotechnology Journal* **19**, 3796–3798 (2021).
16. M. D. Robinson, D. J. McCarthy, G. K. Smyth, edgeR: a Bioconductor package for differential expression analysis of digital gene expression data. *Bioinformatics* **26**, 139–140 (2010).
17. C. W. Law, Y. Chen, W. Shi, G. K. Smyth, voom: Precision weights unlock linear model analysis tools for RNA-seq read counts. *Genome Biol* **15**, R29 (2014).
18. M. E. Ritchie, *et al.*, limma powers differential expression analyses for RNA-sequencing and microarray studies. *Nucleic Acids Res* **43**, e47 (2015).
19. A. Alexa, J. Rahnenführer, T. Lengauer, Improved scoring of functional groups from gene expression data by decorrelating GO graph structure. *Bioinformatics* **22**, 1600–1607 (2006).
20. S. Sayols, rrvgo: a Bioconductor package for interpreting lists of Gene Ontology terms. *MicroPubl Biol* **2023**, 10.17912/micropub.biology.000811.
21. D. T. Gillespie, Exact stochastic simulation of coupled chemical reactions. *J. Phys. Chem.* **81**, 2340–2361 (1977).
22. P. V. Turlapati, K.-W. Kim, L. B. Davin, N. G. Lewis, The laccase multigene family in Arabidopsis thaliana: towards addressing the mystery of their gene function(s). *Planta* **233**, 439–470 (2011).
23. X. Wang, *et al.*, Arabidopsis MICROTUBULE-ASSOCIATED PROTEIN18 Functions in Directional Cell Growth by Destabilizing Cortical Microtubules. *Plant Cell* **19**, 877–889 (2007).
24. Z. Cai, *et al.*, The m6A reader ECT8 is an abiotic stress sensor that accelerates mRNA decay in Arabidopsis. *Plant Cell* **36**, 2908–2926 (2024).

25. R. Quan, *et al.*, SCABP8/CBL10, a Putative Calcium Sensor, Interacts with the Protein Kinase SOS2 to Protect Arabidopsis Shoots from Salt Stress. *Plant Cell* **19**, 1415–1431 (2007).
26. Q. Xie, G. Frugis, D. Colgan, N. H. Chua, Arabidopsis NAC1 transduces auxin signal downstream of TIR1 to promote lateral root development. *Genes Dev* **14**, 3024–3036 (2000).
27. A. Doskočilová, *et al.*, NITRILASE1 regulates the exit from proliferation, genome stability and plant development. *New Phytologist* **198**, 685–698 (2013).
28. S. Fischer, P. Flis, F.-J. Zhao, D. E. Salt, Transcriptional network underpinning ploidy-related elevated leaf potassium in neo-tetraploids. *Plant Physiol* **190**, 1715–1730 (2022).
29. S. Chanroj, S. Padmanaban, D. D. Czerny, G.-Y. Jauh, H. Sze, K<sup>+</sup> transporter AtCHX17 with its hydrophilic C tail localizes to membranes of the secretory/endocytic system: role in reproduction and seed set. *Mol Plant* **6**, 1226–1246 (2013).
30. A.-K. Schürholz, *et al.*, A Comprehensive Toolkit for Inducible, Cell Type-Specific Gene Expression in Arabidopsis. *Plant Physiol* **178**, 40–53 (2018).
31. J. R. Wendrich, *et al.*, Vascular transcription factors guide plant epidermal responses to limiting phosphate conditions. *Science* **370**, eaay4970 (2020).
32. A. Farmer, S. Thibivilliers, K. H. Ryu, J. Schiefelbein, M. Libault, Single-nucleus RNA and ATAC sequencing reveals the impact of chromatin accessibility on gene expression in *Arabidopsis* roots at the single-cell level. *Molecular Plant* **14**, 372–383 (2021).
33. S.-H. Kwak, R. Shen, J. Schiefelbein, Positional signaling mediated by a receptor-like kinase in Arabidopsis. *Science* **307**, 1111–1113 (2005).
34. E.-S. Wallner, *et al.*, Strigolactone- and Karrikin-Independent SMXL Proteins Are Central Regulators of Phloem Formation. *Curr Biol* **27**, 1241–1247 (2017).
35. T. Denyer, *et al.*, Spatiotemporal Developmental Trajectories in the Arabidopsis Root Revealed Using High-Throughput Single-Cell RNA Sequencing. *Dev Cell* **48**, 840-852.e5 (2019).
36. C. N. Shulse, *et al.*, High-Throughput Single-Cell Transcriptome Profiling of Plant Cell Types. *Cell Rep* **27**, 2241-2247.e4 (2019).
37. D. L. Smith, N. V. Fedoroff, LRP1, a gene expressed in lateral and adventitious root primordia of arabidopsis. *Plant Cell* **7**, 735–745 (1995).
38. K. Kumari, *et al.*, Female gametophyte expressed Arabidopsis thaliana lipid transfer proteins AtLtpI.4 and AtLtpI.8 provide a link between callose homeostasis, pollen tube guidance, and fertilization success. [Preprint] (2021). Available at: https://www.biorxiv.org/content/10.1101/2021.01.13.426551v1 [Accessed 18 September 2026].

39. Y. Gao, A. A. Badejo, Y. Sawa, T. Ishikawa, Analysis of Two l-Galactono-1,4-Lactone-Responsive Genes with Complementary Expression During the Development of *Arabidopsis thaliana*. *Plant Cell Physiol* **53**, 592–601 (2012).
40. W. Xuan, *et al.*, Root Cap-Derived Auxin Pre-patterns the Longitudinal Axis of the *Arabidopsis* Root. *Current Biology* **25**, 1381–1388 (2015).
41. M. M. Lee, J. Schiefelbein, Cell Pattern in the *Arabidopsis* Root Epidermis Determined by Lateral Inhibition with Feedback. *Plant Cell* **14**, 611–618 (2002).
42. J.-Y. Lee, *et al.*, Transcriptional and posttranscriptional regulation of transcription factor expression in *Arabidopsis* roots. *Proceedings of the National Academy of Sciences* **103**, 6055–6060 (2006).
43. R. Tominaga-Wada, T. Wada, Regulation of root hair cell differentiation by R3 MYB transcription factors in tomato and *Arabidopsis*. *Front. Plant Sci.* **5** (2014).
44. D. Zheng, *et al.*, PlantscRNADB 4.0: Improved marker identification and annotation under a cell-type uniformity for plants. *Molecular Plant* **19**, 673–688 (2026).
45. K. Kajala, *et al.*, Innovation, conservation, and repurposing of gene function in root cell type development. *Cell* **184**, 3333–3348.e19 (2021).
46. S.-K. Won, *et al.*, cis-Element- and Transcriptome-Based Screening of Root Hair-Specific Genes and Their Functional Characterization in *Arabidopsis*. *Plant Physiol* **150**, 1459–1473 (2009).
47. P. A. Passarinho, A. J. Van Hengel, P. F. Fransz, S. C. de Vries, Expression pattern of the *Arabidopsis thaliana* AtEP3/AtchitIV endochitinase gene. *Planta* **212**, 556–567 (2001).
48. J.-Y. Kim, *et al.*, Distinct identities of leaf phloem cells revealed by single cell transcriptomics. *Plant Cell* **33**, 511–530 (2021).
49. I. Efroni, P.-L. Ip, T. Nawy, A. Mello, K. D. Birnbaum, Quantification of cell identity from single-cell gene expression profiles. *Genome Biol* **16**, 9 (2015).
50. K. Dasgupta, *et al.*, Expression of Sucrose Transporter cDNAs Specifically in Companion Cells Enhances Phloem Loading and Long-Distance Transport of Sucrose but Leads to an Inhibition of Growth and the Perception of a Phosphate Limitation1[W][OPEN]. *Plant Physiol* **165**, 715–731 (2014).
51. C. Zhang, R. A. Barthelson, G. M. Lambert, D. W. Galbraith, Global Characterization of Cell-Specific Gene Expression through Fluorescence-Activated Sorting of Nuclei. *Plant Physiol* **147**, 30–40 (2008).
52. K. Birnbaum, *et al.*, A Gene Expression Map of the *Arabidopsis* Root. *Science* **302**, 1956–1960 (2003).

53. T. Kamiya, *et al.*, The MYB36 transcription factor orchestrates Casparian strip formation. *Proceedings of the National Academy of Sciences* **112**, 10533–10538 (2015).
54. L. M. Liberman, E. E. Sparks, M. A. Moreno-Risueno, J. J. Petricka, P. N. Benfey, MYB36 regulates the transition from proliferation to differentiation in the Arabidopsis root. *Proc Natl Acad Sci U S A* **112**, 12099–12104 (2015).
55. B. De Rybel, *et al.*, A bHLH Complex Controls Embryonic Vascular Tissue Establishment and Indeterminate Growth in Arabidopsis. *Developmental Cell* **24**, 426–437 (2013).
56. N. K. Clay, T. Nelson, Arabidopsis thickvein mutation affects vein thickness and organ vascularization, and resides in a provascular cell-specific spermine synthase involved in vein definition and in polar auxin transport. *Plant Physiol* **138**, 767–777 (2005).
57. L. Muñiz, *et al.*, ACAULIS5 controls Arabidopsis xylem specification through the prevention of premature cell death. *Development* **135**, 2573–2582 (2008).
58. M. Huysmans, *et al.*, NAC Transcription Factors ANAC087 and ANAC046 Control Distinct Aspects of Programmed Cell Death in the Arabidopsis Columella and Lateral Root Cap. *Plant Cell* **30**, 2197–2213 (2018).
